# Nonlinear computation by a habenula-driven recurrent inhibitory network in the raphe

**DOI:** 10.1101/2022.08.31.506056

**Authors:** Michael B. Lynn, Sean Geddes, Mohamad Chahrour, Sébastien Maillé, Léa Caya-Bissonnette, Emerson Harkin, Érik Harvey-Girard, Samir Haj-Dahmane, Richard Naud, Jean-Claude Béïque

## Abstract

Serotonin (5-HT) neurons in the dorsal raphe nucleus (DRN) receive a constellation of long-range inputs, yet guiding principles of local circuit organization and underlying computations in this nucleus are largely unknown. Using inputs from the lateral habenula (LHb) to interrogate the processing features of the DRN, we uncovered 5-HT1A receptor-mediated recurrent connections between 5-HT neurons, refuting classical theories of autoinhibition. Cellular electrophysiology and imaging of a genetically encoded 5-HT sensor revealed that these recurrent inhibitory connections spanned the raphe, were slow, stochastic, strongly facilitating, and gated spike output. These features collectively conveyed highly non-linear dynamics to this network, generating excitation-driven inhibition and winner-take-all computations. *In vivo* optogenetic activation of LHb inputs to DRN, at frequencies where these computations are predicted to ignite, transiently disrupted expression of a reward-conditioned response in an auditory conditioning task. Together, these data identify a core computation supported by an unsuspected slow serotonergic recurrent inhibitory network.

## Introduction

The serotonergic dorsal raphe nucleus (DRN) is a heterogeneous midbrain nucleus which regulates mood and behavior through its widespread efferent innervation (Dorocic et al., 2014; Weissbourd et al., 2014). The coding features of DRN 5-HT neurons still remain largely elusive, although recent studies have shown that they respond to environmental variables such as reward and threat (Cohen et al., 2015; Li et al., 2016; Zhong et al., 2017; Seo et al., 2019), making them suited to supervise behavioral policies based on internal representations of environmental contexts. Moreover, the DRN has been linked to a diverse range of mood-related states such as aggression (Takahashi et al., 2015), patience (Miyazaki et al., 2014; Miyazaki et al., 2018; Miyazaki et al., 2020), and depressive and anxiety disorders (Graeff et al., 1996; Warden et al., 2012). DRN may additionally support cognitive flexibility or modify learning rate (Matias et al., 2017; Grossman et al., 2022). DRN microcircuits are composed of multiple intermingled cell types, including dopaminergic, glutamatergic, serotonergic and GABAergic neurons, organized into functionally distinct modules (Ren et al., 2018). However, the precise rules by which these circuits process incoming information to ultimately provide instructive cues to regulate behavior are largely unknown.

One major source of input to the DRN is from the lateral habenula (LHb), which provides monosynaptic excitatory drive to 5-HT neurons (Zhou et al., 2017). LHb neurons fire in response to punishments, reward omissions and graded threat information by phasic, high-frequency discharges (Matsumoto & Hikosaka, 2009; Proulx et al., 2014; Bromberg-Martin & Hikosaka, 2011; Lecca et al., 2020; Amo et al., 2014; Yang et al., 2018). Recent work has suggested that both the raphe and habenula regulate behavioral policy over long timescales by modifying internal representations (Marques et al., 2020). How incoming synaptic inputs from LHb are processed by DRN microcircuits into an internal state representation that persists over longer timescales to influence behavioral policy is currently unknown.

Here, to elucidate key computations performed by local DRN microcircuits on synaptic inputs, we combined optogenetic, electrophysiological, two-photon imaging, computational and behavioral approaches. Using long-range inputs from LHb to interrogate DRN functional circuitry, we uncovered an unexpected slow inhibitory conductance activated by local recurrent connections between 5-HT neurons. These recurrent connections, mediated by inhibitory 5-HT_1A_ receptors (5-HT_1A_Rs), were slow, highly facilitating, and strongly modulated spiking output of 5-HT neurons. We developed an experimentally-grounded model of the complex short-term dynamics of these recurrent connections, which predicts a sharp transition from net excitation to excitation-driven inhibition and winner-take-all dynamics between long-range inputs to ensembles in DRN. Key predictions of this model were validated in a simple auditory conditioning task. Together, these findings outline highly nonlinear computations supported by an unsuspected recurrent inhibitory network between 5-HT neurons in the DRN.

## Results

### Excitatory monosynaptic LHb inputs to DRN trigger a frequency-dependent indirect inhibition

As a first step towards uncovering organizing principles of microcircuitry within DRN, we activated LHb, a strong long-range input to DRN, and probed neural responses. We utilized a SERT-Cre::tdTom mouse line for whole-cell electrophysiological recordings from genetically identified 5-HT neurons, validating this approach with immunohistochemical staining (Fig. S1A-D; 98.4% overlap between TPH2 and tdTomato expression). Upon bilateral injection of AAV-GFP in LHb, we observed high LHb axon density in DRN (Fig. S2A) consistent with recent work (Zhou et al., 2017). Next, we injected AAV-ChR2(H134R)-mCherry bilaterally into LHb of SERT-Cre::tdTom mice (Fig. 1A and Fig. S1E, see Methods). Photostimulation induced strong and reliable action potentials in LHb neurons which were qualitatively similar to those following current step injections (Fig. S2B-C). Within the DRN, we targeted 5-HT neurons predominantly within the ventromedial DRN, an area exhibiting high LHb axonal density (Fig. S2A; Fig. S3; Zhou et al., 2017). Recording from identified DRN 5-HT neurons from this subregion in the voltage-clamp configuration (Fig. 1C-D), we photostimulated LHb axons with brief (1-3ms) pulses of blue light, triggering synaptic release of glutamate that activated AMPARs (Fig. 1E; V_m_=-70 mV) and NMDARs (Fig. 1F; V_m_=+40mV). LED-stimulated glutamate release was blocked by TTX and rescued by the K^+^ channel blocker 4-AP (Fig. 1G), indicating that LHb makes monosynaptic contacts with DRN 5-HT neurons. These excitatory inputs from LHb were capable of driving 5-HT neurons to spike (Fig. 1H). Thus, LHb afferents directly excite 5-HT neurons, consistent with previous reports (Zhou et al., 2017). While habenula-driven feedforward inhibition onto 5-HT neurons has been previously reported (Zhou et al., 2017; but see Dorocic et al., 2014), we observed little evidence of optically triggered delayed IPSCs in our recording conditions (only 1.7% of 5-HT neurons exhibited IPSCs, and this only upon 20 Hz LHb train photostimulation, n=116). Activation of ChR2-expressing local DRN GABAergic neurons (using a SOM-Cre:: tdTom mouse line) triggered IPSCs in GABAergic neurons (Fig. S4A-G), in agreement with previous work showing local feedforward circuitry in the raphe (Geddes et al., 2016; Zhou et al., 2017). However, optogenetic activation of LHb inputs produced primarily subthreshold excitation of GABAergic neurons (peak depolarization of ∼7mV; Fig. S4H-O), presumably limiting the ability of these long-range inputs to trigger disynaptic inhibition onto 5-HT neurons, at least in our recording conditions and locations. Thus, DRN 5-HT neurons within the ventromedial subdivision of DRN principally receive monosynaptic long-range excitatory input from LHb.

**Figure 1:**
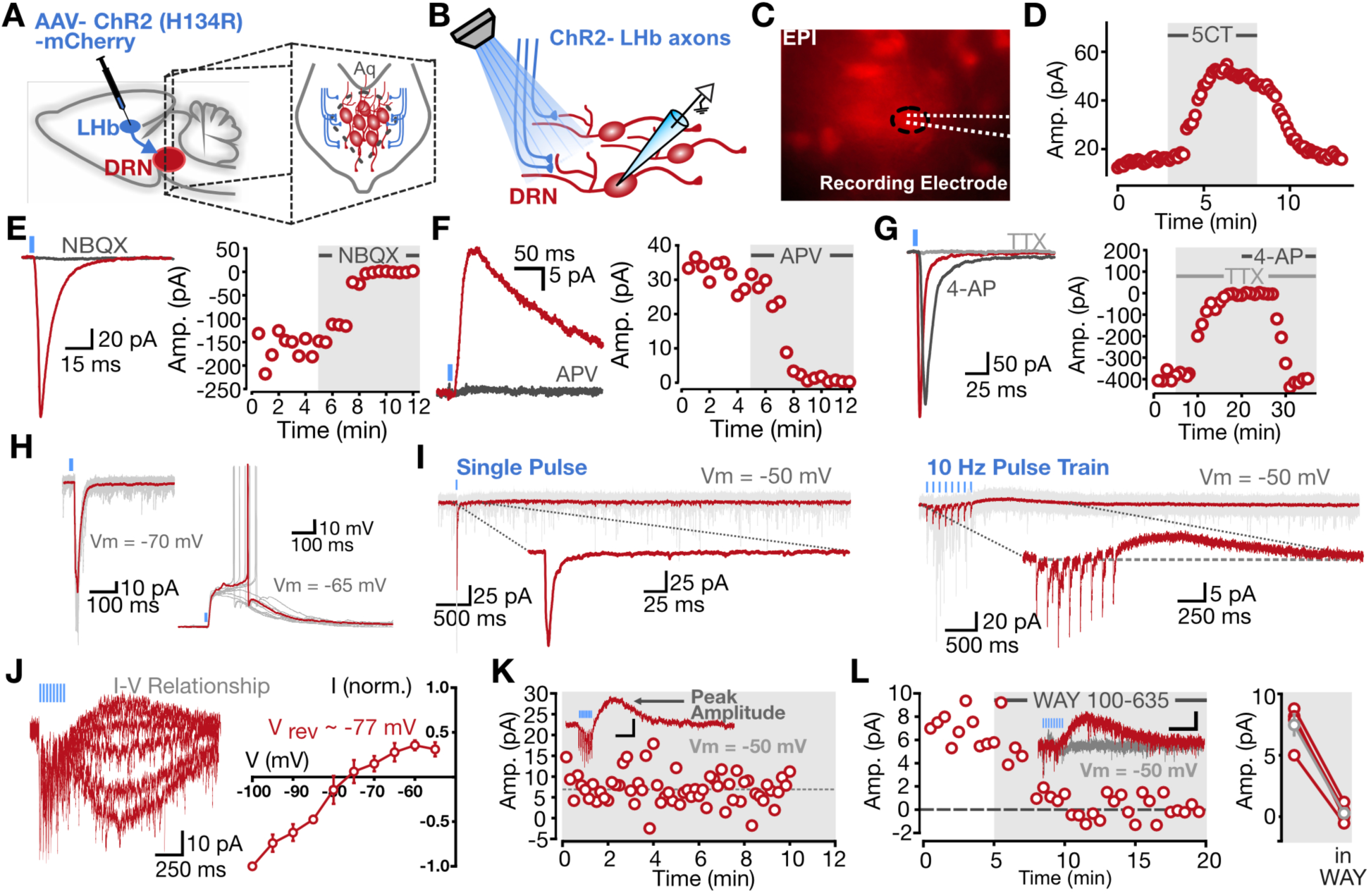
Habenula input to raphe triggers direct excitation and heterosynaptic inhibition in 5-HT neurons. **(A-B)** Schematic of experimental approach. **(C)** Epifluorescent image of tdTomato-positive 5-HT neurons. **(D)** Application of 5-CT (100nM) triggered a transient increase in holding potential. **(E)** Single pulses of light at Vm=-70mV triggered EPSCs (left) which were abolished by application of NBQX (10uM, right). **(F)** At Vm=+40mV, single pulses of light triggered EPSCs (left) which were abolished by application of APV (50uM, right). **(G)** Application of TTX (1uM) blocked EPSCs at Vm=-70mV, while subsequent application of 4-AP (1mM) recovered the current. **(H)** Voltage-clamp recordings of ChR2-EPSCs (Vm = −70 mV) and spiking behavior in response to presynaptic activation. **(I)** Representative 5-HT neuron responses to single pulses of light (left) and pulse trains at 10Hz (right). **(J)** Representative traces of the current-voltage (I-V) relationship of the delayed outward current (left) and group data (right, n=3 neurons, mean +/- SE) **(K)** Representative timecourse of peak amplitude stability over time. **(L)** Representative traces and timecourse showing the blockade of the outward current by WAY 100-635 (400 nM; left). Population data presented in a scatter-line plot (right, Pre-WAY, 7.47 +/- 0.8; Post-WAY, 0.28 +/- 0.4; n = 4 neurons; p = 0.001; paired two-sided t-test). All recordings performed at room temperature; see Fig. S10H-K for replication at 30-32C.

Populations of LHb neurons can discharge at high rates following negative reward prediction errors, depression or stress, or graded threat information (Matsumoto & Hikosaka, 2009, Yang et al., 2018, Dolzani et al., 2016, Amo et al., 2014; Groos & Helmchen, 2024; Andalman et al., 2019). To explore how such input patterns are parsed by DRN microcircuits, we delivered trains of optical stimuli at a range of frequencies consistent with previously reported *in vivo* phasic habenular responses (Amo et al., 2014; Groos & Helmchen, 2024). Train stimulation at 10Hz or 20Hz, surprisingly, triggered the appearance of a delayed outward current lasting hundreds of milliseconds, following the initial monosynaptic EPSCs (Fig. 1I). The overall kinetics of this protracted current, as well as its underlying rectifying I-V curve displaying a reversal potential at ∼-77mV (Fig. 1J), suggest the involvement of a G-protein inwardly rectifying potassium (GIRK) channel (Newberry & Nicoll, 1984). While several G-protein coupled receptor subtypes couple to GIRK channels, the inhibitory 5-HT receptor of the 5-HT_1A_ subtype are expressed at high level in DRN 5-HT neurons (Pazos & Palacios, 1985) and were thus reasonable candidates. Bath application of the 5-HT_1A_R antagonist WAY100-635 abolished the outward current triggered by burst activation of LHb axons in the DRN (Fig. 1K-L) without influencing the light-induced EPSCs (pre-WAY amplitude, 25.2 +-12.5pA; post-WAY amplitude, 17.9 +-6.9pA; n=3 neurons, p=0.48, paired t-test). Across all recordings from genetically identified 5-HT neurons, 33.9% showed no response to optogenetic LHb stimulation, 50.8% exhibited direct excitation only, 12.7% exhibited direct excitation and indirect 5-HT_1A_ inhibition, and 0.84% exhibited only indirect 5-HT_1A_ inhibition. Together, these results demonstrate that trains of LHb input to DRN 5-HT neurons trigger rapid monosynaptic glutamatergic-mediated excitation followed by a slower, 5-HT_1A_R-mediated inhibition.

### DRN 5-HT neurons are organized in a recurrent inhibitory network

We next sought to identify the cellular and network underpinnings of this non-canonical mixed excitation/inhibition response, initially focusing on the protracted 5-HT_1A_R-mediated inhibition. A dominant and well-accepted model posits that 5-HT1ARs are autoreceptors. In this model, individual 5-HT neurons, upon firing, limit their own excitability through the dendritic release of 5-HT (Maxwell et al., 1983; Ridet et al., 1994; Bunin & Wightmann, 1998), which activates these so-called inhibitory somatodendritic 5-HT_1A_ autoreceptors in an autocrine fashion (Gozlan et al., 1983; Weissmann-Nanopoulos et al., 1985; Pineyro & Blier, 1999). This canonical autoreceptor model is nonetheless difficult to reconcile with our observations that train activation of LHb inputs triggered 5-HT_1A_R-mediated outward currents in nonspiking, voltage-clamped 5-HT neurons (Fig. 1I,J), or in 5-HT neurons receiving no functional excitatory LHb inputs (Fig. S2D-E). We therefore next sought to experimentally disambiguate the autoreceptor hypothesis from the alternative hypothesis that DRN 5-HT neurons are organized in a recurrent inhibitory network architecture (Fig. 2A).

**Figure 2:**
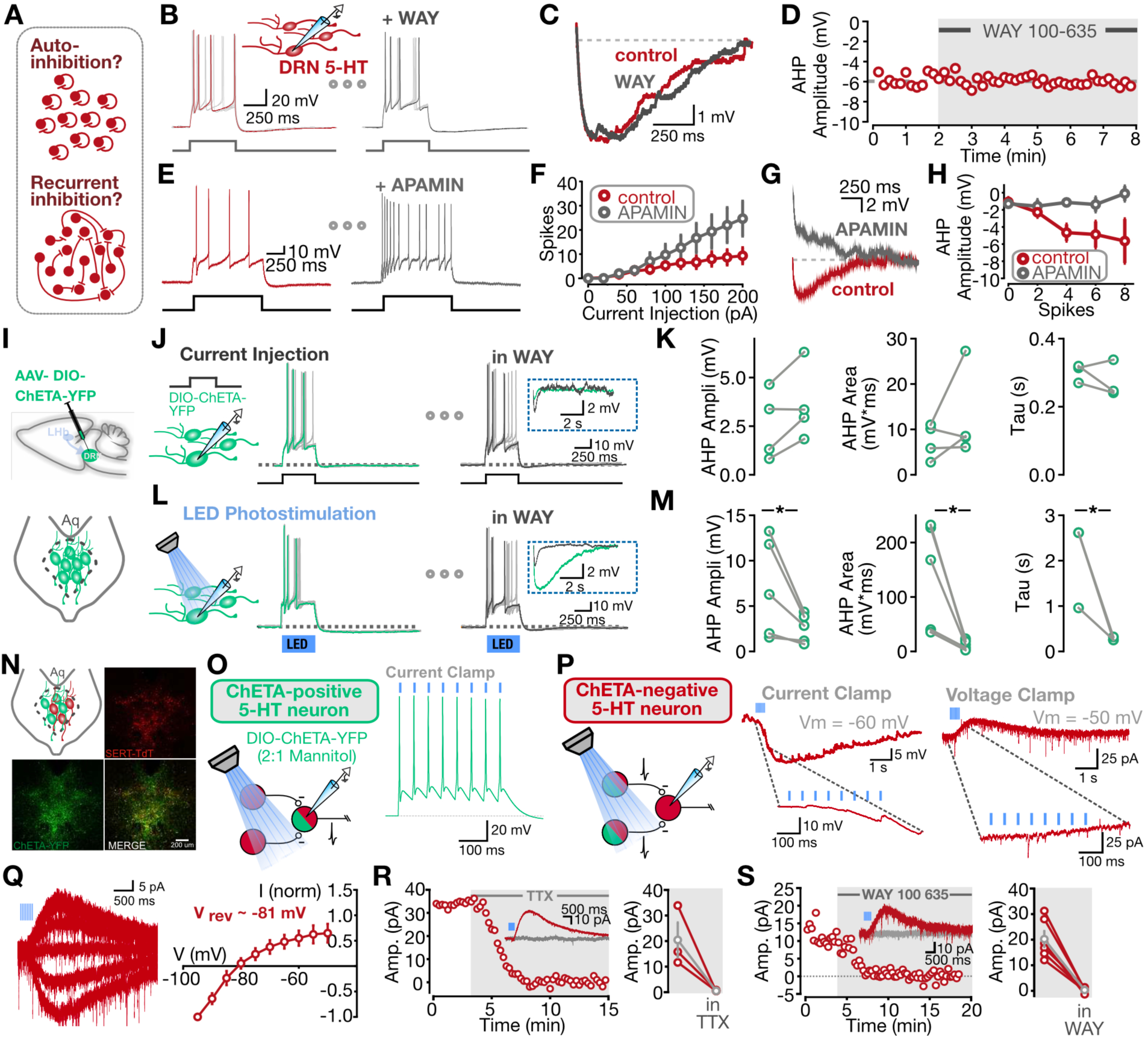
5-HT neurons in raphe are organized in a recurrent inhibitory network. **(A)** Two hypothesized mechanisms of 5-HT1AR-based signaling in raphe. **(B)** Representative traces of spiking in response to direct current injection before and after WAY 100-635 application. **(C)** Direct comparison of AHP before and after application of WAY 100-635. **(D)** Representative scatter plot showing no change in the AHP amplitude with bath application of WAY 100-635. **(E)** Representative traces of spiking in response to direct current injection before and after APAMIN application. **(F)** Relationship between current injection strength and evoked spikes, in the presence or absence of APAMIN (n=3 neurons, mean +/- SE) **(G)** Direct comparison of AHP before and after application of APAMIN. **(H)** Relationship between spike number and AHP amplitude, in the presence or absence of APAMIN (n=3 neurons, mean +/- SE). **(I)** Schematic of local optogenetic experimental approach. **(J)** Representative traces from CHETA-expressing 5-HT neurons showing current step-driven spiking before (green) and after (black) bath application of WAY100-635. Grey lines represent single sweeps from the same neuron. **(K)** Quantification of AHP statistics for control and WAY100-635 conditions (AHP amplitude, p=0.96, n=4 neurons; AHP area, p=0.85, n=4 neurons; AHP tau, p=0.09, n=3 neurons; paired two-sided t-tests). Grey lines represent single sweeps from the same neuron. **(L)** Representative traces from CHETA-expressing 5-HT neurons showing photostimulation-driven spiking before (green) and after (black) bath application of WAY100-635. **(M)** Quantification of AHP statistics for control and WAY100-635 conditions (AHP amplitude, p=0.03, n=5 neurons; AHP area, p=0.03, n=5 neurons; AHP tau, p=0.02, n=3 neurons; paired two-sided t-tests). **(N)** Schematic and confocal images of sparse local optogenetic approach (AAV-DIO-CHETA-YFP diluted in mannitol 2:1). **(O)** While recording from an opsin-expressing neuron, brief optical stimulation could reliably elicit spiking behavior. **(P)** While recording from a non-opsin-expressing serotonergic neuron, photostimulation at 20Hz produced an inhibitory current; note the absence of direct photocurrents. **(Q)** Representative traces of the current-voltage (I-V) relationship of the inhibitory current (left) and group data (right, n=5 neurons, mean +/- SE) **(R)** Representative traces and timecourse of TTX (1uM) bath application (left) and group data (right; n=3 neurons; grey average trace: mean +/- SE). **(S)** Representative traces and timecourse of WAY100-635 (400nM) bath application (left) and group data (right; n=6 neurons; grey average trace: mean +/- SE). All recordings performed at room temperature.

We first reasoned that if 5-HT_1A_Rs were indeed autocrine inhibitory autoreceptors, evoked spikes from a single 5-HT neuron should elicit a 5-HT1AR-mediated inhibition in the same isolated neuron, contributing to the post-spike afterhyperpolarization (AHP) commonly observed in 5-HT neurons (Vandermaelen & Aghajanian, 1983). Whole-cell recordings, however, revealed that the prominent post-spiking AHP triggered by current injection was insensitive to application of the 5-HT_1A_R antagonist WAY100-635 (Fig. 2B-D), yet fully abolished by application of the SK channel blocker apamin (Fig. 2E-H), demonstrating that the slow AHP in 5-HT neurons is mediated by SK channels, and not 5-HT_1A_Rs. The absence of autocrine effects on the AHP could be explained by the small number of action potentials emitted in current-clamp experiments. Voltage-clamp experiments further showed that the robust outward current that followed long, constant, depolarizing voltage steps in 5-HT neurons was also sensitive to apamin, but not to WAY100-635 (Fig. S5). Together, these experiments failed to support an autocrine function for 5-HT_1A_Rs on 5-HT neurons.

To begin exploring the alternative possibility of a recurrent network between 5-HT neurons in the DRN, we designed a series of targeted optogenetic strategies based on viral transduction of 5-HT neurons with a Cre-dependent AAV ChETA-YFP in SERT-Cre::tdTom mice (Fig. S5A, see Methods). *Ex vivo* slice recordings from DRN 5-HT neurons co-expressing tdTomato and YFP-tagged ChETA opsin (Fig. S7B-C) showed that LED stimulation induced robust inward photocurrents and depolarizations that induced spiking reliably up to ∼20 Hz (Fig. S7D-E). Train photostimulation of all transduced 5-HT neurons triggered a robust inhibitory 5-HT_1A_R conductance (Fig. S7F-G). We then reasoned that neuronal spiking induced by photostimulating ChETA-expressing 5-HT neurons should reveal an additive 5-HT_1A_R mediated component of the resultant AHP in the recorded neuron if DRN 5-HT neurons were indeed interconnected in a recurrent inhibitory network. To test this, we cross-calibrated single-cell direct current injections and LED intensity such that both methods independently triggered spiking of similar frequency (Fig. 2J and 2L). While the spiking triggered by direct current injections induced a strong and highly reliable AHP in these neurons (*i.e.,* tdTomato and YFP-tagged ChETA-positive) that was not affected by the 5-HT_1A_R antagonist WAY100-635 (Fig. 2K-L), the same spiking triggered by LED photostimulation of the entire DRN network induced a substantially larger AHP, and the additional component was blocked by WAY100-635 (Fig. 2M). These results fail to support the autocrine hypothesis of 5-HT_1A_R function, and rather suggest that 5-HT neurons are rather organized in a recurrent inhibitory network.

To more directly test the presence of recurrent connectivity in the raphe, we next developed an AAV-ChETA-YFP transduction strategy in the DRN of SERT-Cre::tdTom mice (Fig. 2N) to achieve sparse, mosaic-like, expression of ChETA-YFP in 5-HT neurons (see Methods). This allowed us to obtain visually targeted recordings either from ChETA-expressing 5-HT neurons (*i.e.,* YFP and tdTomato expressing neurons that exhibited reliable optically driven spiking; Fig. 2O) or from non ChETA-expressing 5-HT neurons (*i.e*., YFP-negative, tdTomato-positive 5-HT neurons). Targeted recordings from non ChETA-expressing 5-HT neurons revealed that high-frequency LED photostimulation (which directly drives the surrounding network but not the recorded neuron) induced delayed, strongly rectifying, outward currents (Fig. 2P-Q) which were blocked by TTX and by WAY 100-635 (Fig. 2R-S). Taken together, these optogenetic data strongly support the existence of an unsuspected recurrent inhibitory network in DRN, consisting of 5-HT neurons communicating via 5-HT_1A_Rs.

To further explore connectivity features between 5-HT neurons in the DRN, we performed small, visually guided injections of fluorescent dextran retrograde tracer into either dorsal DRN or ventral DRN in acute slices from SERT-Cre::tdTom mice (dorsal DRN injection diameter: 105.6 +-37.7um, ventral DRN injection diameter: 91.0 +-22.4um; p=0.69, independent t-test, n=3 injections each). We quantified the locations of neurons expressing both fluorescent dextran and tdTomato signals (Fig. S7A; see Methods; Trinh et al., 2016), revealing dual-labeled neurons several hundred microns outside of the injection site (Fig S7B). The locations of both dual-labeled neurons and single-labeled tdTomato-expressing 5-HT neurons, were extracted for each individual injection (Fig. S7C). Pooling across all injections revealed a spatial organization of labeled neurons relative to the injection center that was distinct for dorsal and ventral injections (Fig. S7D). Labeled neurons were found at distances up to ∼500 µm from the injection center, and the distribution of distances from the injection site was significantly different between dorsal and ventral injections (Fig. S7E; p=1.29*10^-3; Kolmogorov-Smirnov test), suggesting submodule-specific differences in spatial connectivity profiles. Furthermore, dorsal injections were most similar to unstructured connectivity (random sampling from all 5-HT neurons), while ventral injections had more local organization of connectivity (Fig. S7F; see Methods). As a final step, we computed a polar plot of the angle of labeled neurons relative to the injection center, which for dorsal injections revealed an enrichment in ventrally located labeled 5-HT neurons, and for ventral injections led to a more homogeneous distribution of labeled neurons in polar coordinates (Fig. S7G-H). Together, these experiments outline a set of spatial and anatomical features of the recurrent inhibitory network within DRN.

### A quantitative description of release dynamics at 5-HT_1A_R-mediated recurrent connections

We next parameterized key functional features of the serotonergic recurrent inhibitory network. Train optogenetic stimulation (20Hz, 8 pulses) of 5-HT neurons triggered a 5-HT_1A_R conductance that peaked approximately 400 ms following the end of the train (392.6 ± 106.9ms) and that slowly decayed (2.1 ± 0.46s). Varying the number of light pulses delivered at 20Hz revealed a relationship between pulse number and postsynaptic conductance that was almost linear, indicating that 5-HT_1A_R coupling to GIRK channels can follow multiple successive activation by endogenous 5-HT with limited desensitization (Fig. S8A-C). To infer presynaptic release dynamics, we next calculated the response variance as a function of pulse number, finding virtually no change in variance despite changes in amplitude (Fig. S8D). While such dynamics are in principle consistent with deterministic neurotransmitter release (*i.e.,* P_rel_ ∼1; see Methods for full details), artefactual inflation of presynaptic release probability has previously been reported for optogenetic synaptic terminal activation (Zhang & Oertner, 2007) (see Methods). These results nonetheless outline the timecourse of a GIRK- and 5-HT_1A_R -mediated synapse that quasi-linearly reports presynaptic 5-HT release over a physiological range (see below).

To monitor 5-HT presynaptic release dynamics in the DRN circumventing the confound brought about by optogenetic stimulation, we next recorded DRN 5-HT neurons while electrically stimulating nearby serotonergic fibers with a bipolar concentric electrode (Fig. 3A). A cocktail of synaptic blockers was bath-applied to isolate serotonergic transmission (Fig. 3A, see Methods). High-frequency electrical stimulation readily triggered a 5-HT_1A_R -mediated outward current with a similar time course to that induced by LED stimulation (Fig. 3B). The subset of 5-HT neurons displaying an electrically evoked outward current did not exhibit clear anatomical organizing principles within the ventral DRN (Fig. S3B; circles filled with red). We next delivered isolated electrical pulses under a low-intensity minimal stimulation regime (*i.e.,* activating small numbers of individual 5-HT synapses) while monitoring 5-HT_1A_R -mediated currents. These recordings showed instances of clear successes (∼5 pA) and failures (Fig. 3C) of 5-HT release, consistent with small numbers of 5-HT synapses each exhibiting a P_rel_ < 1 (Lynn et al., 2020). Fig. 3D depicts the distribution of event charges plotted alongside neuron-specific control distributions (obtained from integrating an equivalent duration of noise; see Methods). The bimodal event distribution, exhibiting a left peak overlapping with the noise distribution, is consistent with probabilistic 5-HT transmission.

**Figure 3:**
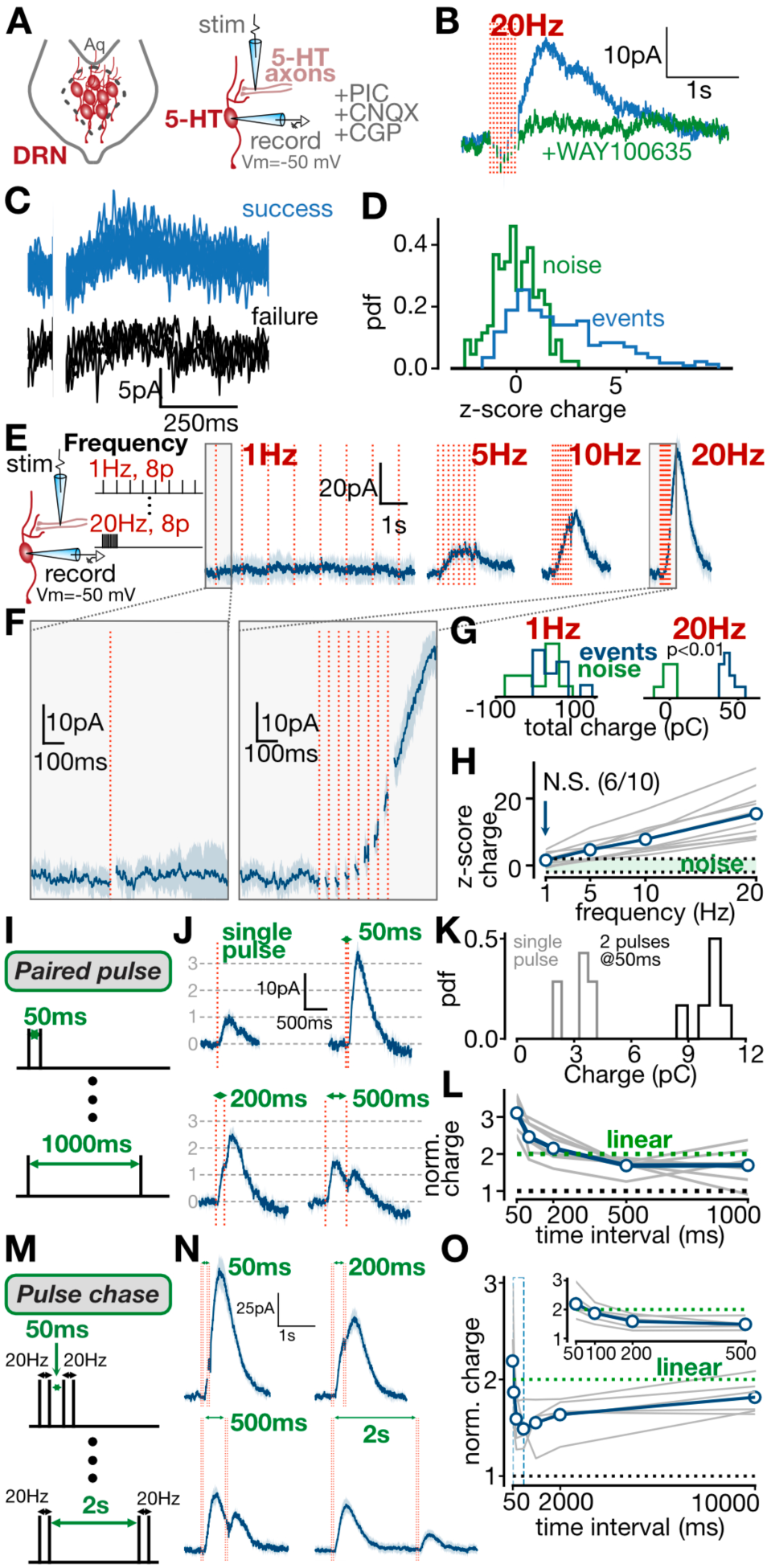
Serotonin release is probabilistic and strongly facilitating at inhibitory recurrent connections. **(A)** Experimental schematic. **(B)** Electrical stimulation at 20Hz evoked an outward current that is blocked by WAY100-635. **(C)** Representative traces depicting responses to low-amplitude electrical stimulation, showing sorted successes and failures from the same neuron. **(D)** Histograms of charge z-score, calculated for post-stimulus events (blue) and an equivalent period of non stimulus-driven noise from the same neuron (green) (n=424 events from 4 neurons). **(E)** Experimental setup for frequency-varying electrical stimulation experiments (left). Representative averaged traces from a single neuron depicting responses to electrical stimulation at various frequencies (right). **(F)** Expanded traces showing responses to 1Hz and 20Hz stimuli. **(G)** Charge histograms for stimulus-driven events and noise (one example neuron; 1Hz: p>0.05, 20Hz: p<0.01, two-sided t-tests). **(H)** Charge z-score as a function of frequency, with 95% noise C.I shown in green (n=10 neurons, two-sided t-tests for each neuron as in G). **(I)** Paired-pulse electrical stimulation paradigm. **(J)** Representative averaged traces from a single neuron depicting responses to electrical stimulation at various paired-pulse intervals. **(K)** Charge histograms for single-pulse responses and paired-pulse responses at 50ms for one example neuron. **(L)** Post-stimulus charge as a function of paired-pulse time interval (n=7 neurons). Green line depicts linear summation. **(M)** Pulse-chase electrical stimulation paradigm. **(N)** Representative averaged traces from a single neuron depicting responses to pulse-chase electrical stimulation protocol at various temporal intervals. **(O)** Post-stimulus charge as a function of pulse-chase time interval (n=6 neurons). Green line depicts linear summation. All recordings performed at 30-32C and [Ca2+]=2.5mM; see Fig. S10A-G for replication at lower external calcium concentrations.

We next examined the short-term dynamics of these 5-HT synapses by delivering electrical stimulation trains with a fixed pulse number (8) and variable frequency (Fig. 3E). A deterministic synapse should exhibit identical responses over a range of input frequencies if pulse number is fixed. While 1 Hz stimulation produced a notably weak response, higher frequencies of 10 Hz and 20 Hz elicited progressively larger inhibitory currents (Fig. 3E-F). To directly compare these protracted responses, which displayed peak amplitudes that were highly dependent on stimulus parameters, we employed a z-score based metric of total charge. Briefly, we computed the total charge evoked by each stimulus train, and compared this with a surrogate noise distribution constructed by performing the same integration on an equivalent duration of baseline recording (example neuron shown in Fig. 3G; two-sided t-test on trial distribution of charges for events and noise; See Methods). While low-frequency (1 Hz) stimuli did not produce responses significantly different from the surrogate noise distribution (Fig. 3H; 6/10 neurons with p>0.05 for event *vs* noise charges, two-sided t-test), high-frequency (20 Hz) stimuli produced progressively larger responses (Fig. 3H), consistent with short-term facilitation.

Altogether, we conclude that the protracted 5-HT1AR-mediated inhibition reflects the probabilistic release of 5-HT that exhibits substantial short-term facilitation.

To further quantify the timescale of these short-term synaptic dynamics, we delivered patterned electrical stimulation, starting with classic paired-pulse stimulation experiments. When paired-pulse stimuli were closely spaced (50-200 ms) we observed supralinear summation compared to isolated single pulses; in contrast, paired-pulse intervals of 500 ms and longer produced depressing responses (Fig. 3J-K). This short-term depression observed at timescales of seconds is consistent with previous reports on 5-HT_1A_R-mediated inhibition (Courtney & Ford, 2016). By comparing the actual evoked charge with the predicted linear summation of isolated single-pulse responses, we quantified the dynamics of these recurrent synapses across multiple timescales (Fig. 3L). Together, these results indicate that the short-term dynamics of these 5-HT synapses are dominated by strong short-term facilitation at short intervals, transitioning to mild depression at timescales > 500 ms.

In the experiments above, we aimed to approach conditions of minimal stimulation (i.e., successes and failures). We additionally repeated these experiments with stronger electrical stimulation (Fig. S9) and under low extracellular calcium conditions (Fig. S10A-G; [Ca^2+^]=1.5 mM), and found that 5-HT release dynamics still exhibited strong short-term facilitation, especially for short inter-pulse intervals in these conditions. Likewise, the recruitment of 5-HT_1A_R-mediated inhibition by high frequency optogenetic activation of LHb inputs to the DRN (Fig. 1I) still remained in these conditions ([Ca^2+^]=1.5 mM; 30-32 °C; Fig. S10H-K).

To examine meta-dynamic features of these 5-HT synapses, we delivered a second set of patterned electrical stimuli consisting of a paired-pulse stimulation (20 Hz; ISI of 50 ms) separated from another paired-pulse stimulation by some varying time interval (pulse-chase; Fig. 3M). These experiments revealed, unexpectedly, that the paired-pulse facilitation at 20 Hz is highly dependent on stimulus history over a timescale of seconds. When paired-pulse sets were separated by short intervals, facilitation was present for both sets, yielding a supralinear response to the patterned input (Fig. 3N). However, when separated by intervals of >500 ms, the second paired-pulse stimulus elicited notably smaller responses (less facilitation), indicating release meta-dynamics, or a history-dependent change in facilitation (Fig. 3N). Charge quantification metrics, similar to above (see Methods), revealed that this dependence on stimulation history lasted up to 10 seconds (Fig. 3O).

Any effort to identify the network computations sustained by this recurrent inhibitory network requires, as an obligatory intermediate step, an accurate quantitative model of the synaptic dynamics described above. We therefore next developed a mathematical model capturing these ground-truth short-term dynamics of 5-HT synapses. We first fitted the experimentally-derived 5-HT_1A_R -mediated inhibitory GIRK current with a simple summed exponential model (Fig. 4A). Next, to adequately capture the short-term dynamics of 5-HT synapses, we fitted our experimental data using a previously described spike-response plasticity model (SRP; Rossbroich et al., 2021; see Methods). Briefly, a presynaptic spiketrain is convolved with a kernel indicating the sign and timescale of synaptic dynamics, passed through a sigmoidal readout, and convolved at each presynaptic spike-time with a postsynaptic event kernel (here, describing a GIRK-mediated current). Using the paired-pulse electrical stimulation data as input, we used a least-squares approach to find a best fit, yielding an optimized short-term plasticity (STP) kernel that showed a short-timescale facilitation and long-timescale depression components (Fig. 4B). The resultant model effectively captured the dynamics of the paired-pulse experimental data across all timescales tested (Fig. 4C). Remarkably, despite not being fitted on this data, the model described the meta-facilitation dynamics observed in the pulse-chase experiments – in effect acting as a test set for the model (Fig. 4D). This likely reflects the fact that the kernel has a short window of facilitation, which afterwards tends towards depression if subsequent stimuli arrive too late (such as in the pulse-chase experiments with extended time intervals; see Methods). To provide further intuition about this model, we conducted simple simulations where presynaptic trains were delivered to single neurons, and the model returned the kinetics and response magnitude of our earlier electrical stimulation experiments (Fig. 4E). Together, these results show that the complex dynamics of 5-HT release in the DRN can be faithfully captured by a SRP model.

**Figure 4:**
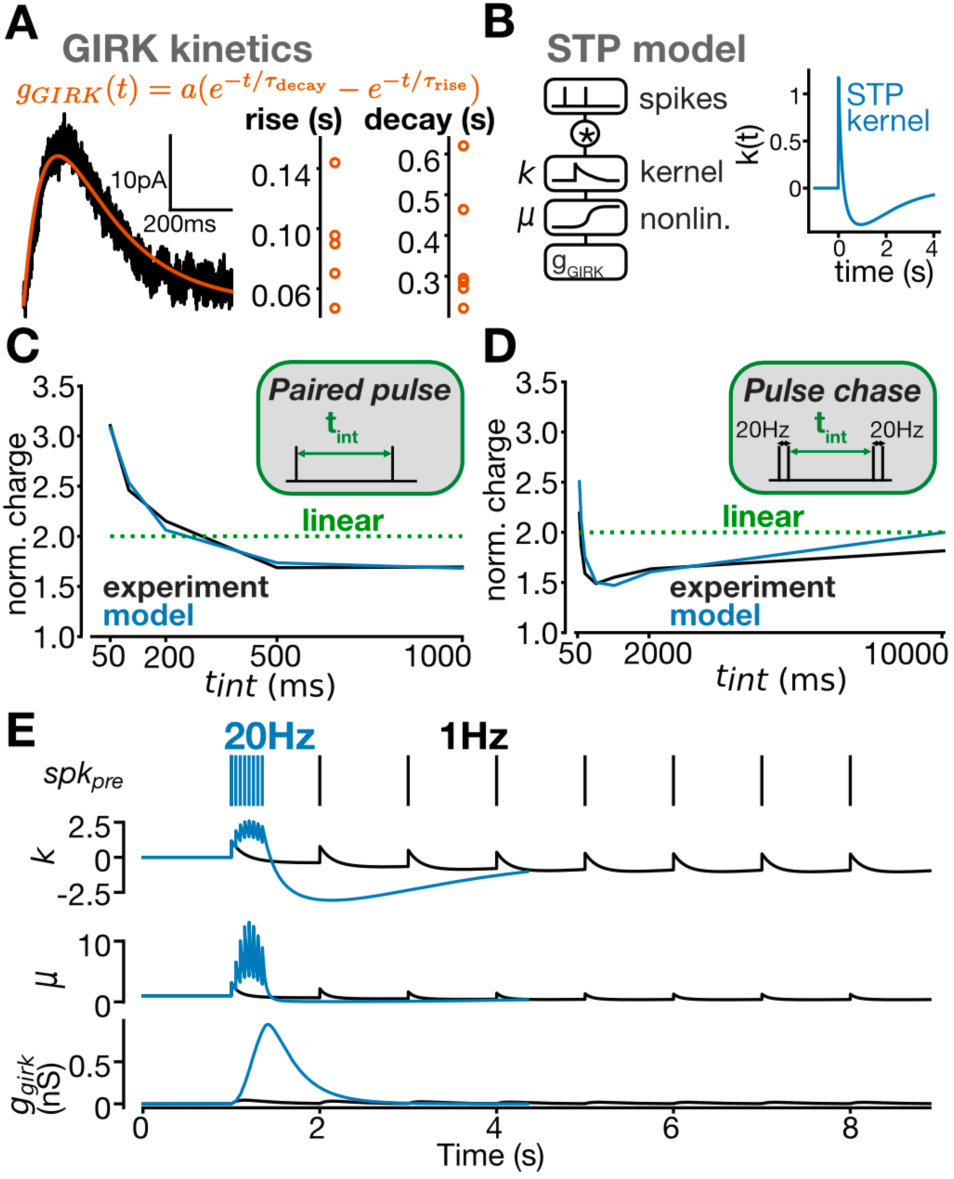
A quantitative model of short-term release dynamics captures patterned inputs. **(A)** Representative trace of kinetics fitted to an averaged electrically evoked GIRK current, and parameter distribution across the population (n=6 neurons). **(B)** Schematic depicting the linear-nonlinear short-term plasticity model and the plasticity kernel fitted using a least-squares approach to paired-pulse data. **(C)** Post-stimulus charge as a function of paired-pulse time interval, depicting experimental mean (black) and model predictions with fitted kernel and nonlinearity (blue). **(D)** Post-stimulus charge as a function of pulse-chase time interval, depicting experimental mean (black) and model predictions (blue). Note that model was fitted to paired-pulse, not pulse-chase dataset. **(E)** Simulations showing responses to 1Hz (black) and 20Hz (blue) presynaptic input, the model variables, and the resulting GIRK conductance.

### Synaptic facilitation dynamics and the spatial structure of 5-HT release in DRN

We next explored spatial features of 5-HT release in e*x vivo* slices of the DRN using two-photon imaging of the genetically encoded 5-HT sensor GRAB5-HT2M expressed selectively in 5-HT neurons (Fig. 5A). Bath application of 5-HT (50 µM) generated a large increase in GRAB_5-HT_ fluorescence in both neurites and cell bodies (Fig. 5B), acting as a positive control for GRAB_5-HT_ expression and overall function. We next electrically stimulated 5-HT connections in DRN slices at various stimulation frequencies (5-20Hz; while fixing pulse number at 8, similar to above; see Fig. 3 and Methods). We manually annotated responding regions and observed a strikingly nonlinear increase in fluorescence response as electrical stimulation frequency increased (Fig. 5C-D). GRAB_5-HT_-mediated fluorescence was sensitive to TTX application, demonstrating that electrically induced release events were action potential-driven (Fig. S11). By imaging 5-HT release directly, these data present a separate line of evidence showing facilitation dynamics of 5-HT release as revealed earlier by whole-cell recordings (Fig. 3), and additionally support a presynaptic locus for this phenomenon.

**Figure 5:**
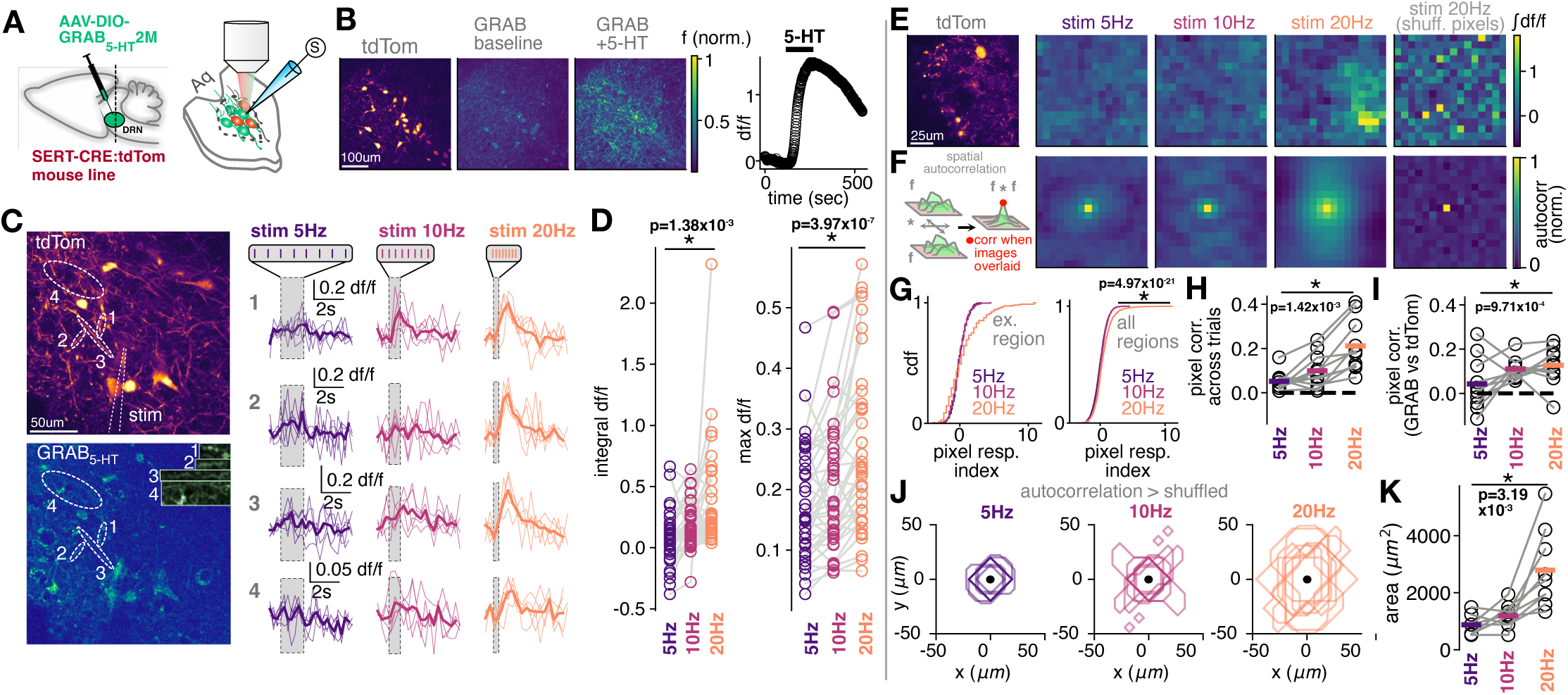
Synaptic facilitation dynamics and spatial structure of release revealed by GRAB5-HT serotonin sensor imaging in DRN. **(A)** Experimental schematic demonstrating 2P GRAB5-HT imaging approach. **(B)** Bath application of 5-HT generates an increase in GRAB signal. *Left*: fluorescence max-projection images from an example field of view demonstrating tdTomato signal, baseline GRAB signal before 5-HT application, and GRAB signal during 5-HT application. *Right*: quantification of whole-frame GRAB fluorescence during timecourse of 5-HT washon. **(C)** GRAB fluorescence responses are frequency-dependent. *Left*: Example region showing max-projection tdTomato and GRAB fluorescence images, as well as manually identified responsive ROIs (dotted ovals and inset). *Right*: GRAB sensor responses to electrical stimulation at 5, 10 and 20Hz (shaded regions) for the ROIs depicted on the left. Thin traces represent individual trials and thick traces represent means across all trials. **(D)** Frequency dependence of GRAB responses across ROIs (n=39), calculated as response integral (left) or response maximum (right; paired t-test between 5 and 20Hz data). **(E)** Example whole-frame tdTomato fluorescence (left) and binned GRAB responses to 5, 10 and 20Hz electrical stimulation (8 pulses), including 20Hz shuffled pixel control (right). **(F)** Spatial autocorrelation plots for the frames shown in E. Autocorrelation values are normalized from 0 to 1 for presentation. **(G)** Left: Pixelwise cumulative distribution of GRAB responses at 5, 10 and 20Hz for example region shown in E (pixel response index is expressed as a z-score vs 5Hz data). Right: Same as left, but across all regions (n=11; Kolmogorov-Smirnov test for 5Hz vs 20Hz) **(H)** Mean pixelwise Pearson R correlation of GRAB response across trials of repeated electrical stimulation at 5, 10 and 20Hz (n=11 regions; paired T-test, 5Hz vs 20Hz). **(I)** Mean pixelwise Pearson R correlation of GRAB response during stimulation, and tdTomato fluorescence (n=11 regions; paired T-test, 5Hz vs 20Hz). **(J)** Spatial clustering of GRAB5-HT signal is frequency-dependent. Shown are spatial masks where autocorrelation of GRAB signal significantly exceeds autocorrelation on shuffled frames (p=0.0001), with all regions overlaid (n=11 regions). **(K)** The total area of spatial masks in J at 5, 10 and 20Hz (n=11 regions; paired T-test, 5Hz vs 20Hz).

As stimulation frequency increased, GRAB_5-HT_ -mediated fluorescence increased in absolute intensity and became progressively more localized to hotspots within the imaged frame (Fig. 5E, left; Fig. 5G). To assess the reliability of the fluorescence signal, we computed the pairwise pixel correlation across repeated trials, and found that the response became more spatially reliable with higher stimulation frequencies (Fig. 5H), as expected for a probabilistic and facilitating population of synapses. GRAB_5-HT_ fluorescence displayed a weak overall correlation with tdTomato signal, likely reflecting a lack of strong somatic signal for the biosensor (Fig. 5I).

Our whole-frame analyses (Fig. 5E) suggested that GRAB_5-HT_ fluorescence responses were spatially restricted (*i.e.,* hotspots). To quantify the spatial features of these hotspots in a principled manner, we next computed the spatial autocorrelation of the GRAB_5-HT_ whole-frame response for each stimulation frequency (Fig. 5F, left; see Methods). As frequency increased, the spatial autocorrelation function became broader, providing a metric of hotspot size (Fig. 5F). To illustrate the expected outcome of the autocorrelation function should no spatial organization of the GRAB_5-HT_ response be present, we performed spatial shuffling of the sensor response pixels at 20Hz (Fig. 5E, right). This produced an autocorrelation function that sharply peaked in the center (correlation when images are overlaid) (Fig. 5F, right). We constructed a null distribution of the autocorrelation values arising from these shuffled 5-HT sensor response pixels for each condition (Fig. S12A-E; see Methods). By comparing this shuffled distribution with the real distribution of stimulation-evoked autocorrelation values (Fig. S12E), we were able to delineate a mask of statistically significant autocorrelation values for each region and stimulation condition (Fig. S12F). These masks, representing a metric of hotspot size of the GRAB_5-HT_ fluorescence signal, showed greater area as the stimulation frequency increased (Fig. 5J-K). Together, these data indicate that GRAB_5-HT_ fluorescence exhibits spatially localized hotspots that scale in both area and intensity with stimulation frequency, consistent with facilitating 5-HT release.

### Computations performed by a recurrent inhibitory network under excitatory drive

To identify the network computations in DRN that arise from recurrent connections between 5-HT neurons, dominated by short-term facilitation, we constructed a network model that combined our mathematical description of the short-term plasticity of 5-HT synapses (Fig. 4) with previously described excitability parameters for 5-HT neurons (Fig. 6A; Harkin et al., 2023; see Methods). This model consisted of a population of leaky-integrate-and-fire 5-HT neurons, with sparse recurrent connections imbued with the described short-term recurrent plasticity rules. 5-HT neurons received background depolarizing Poisson input, and a subset of 5-HT neurons received input from long-range targets (see Methods).

**Figure 6:**
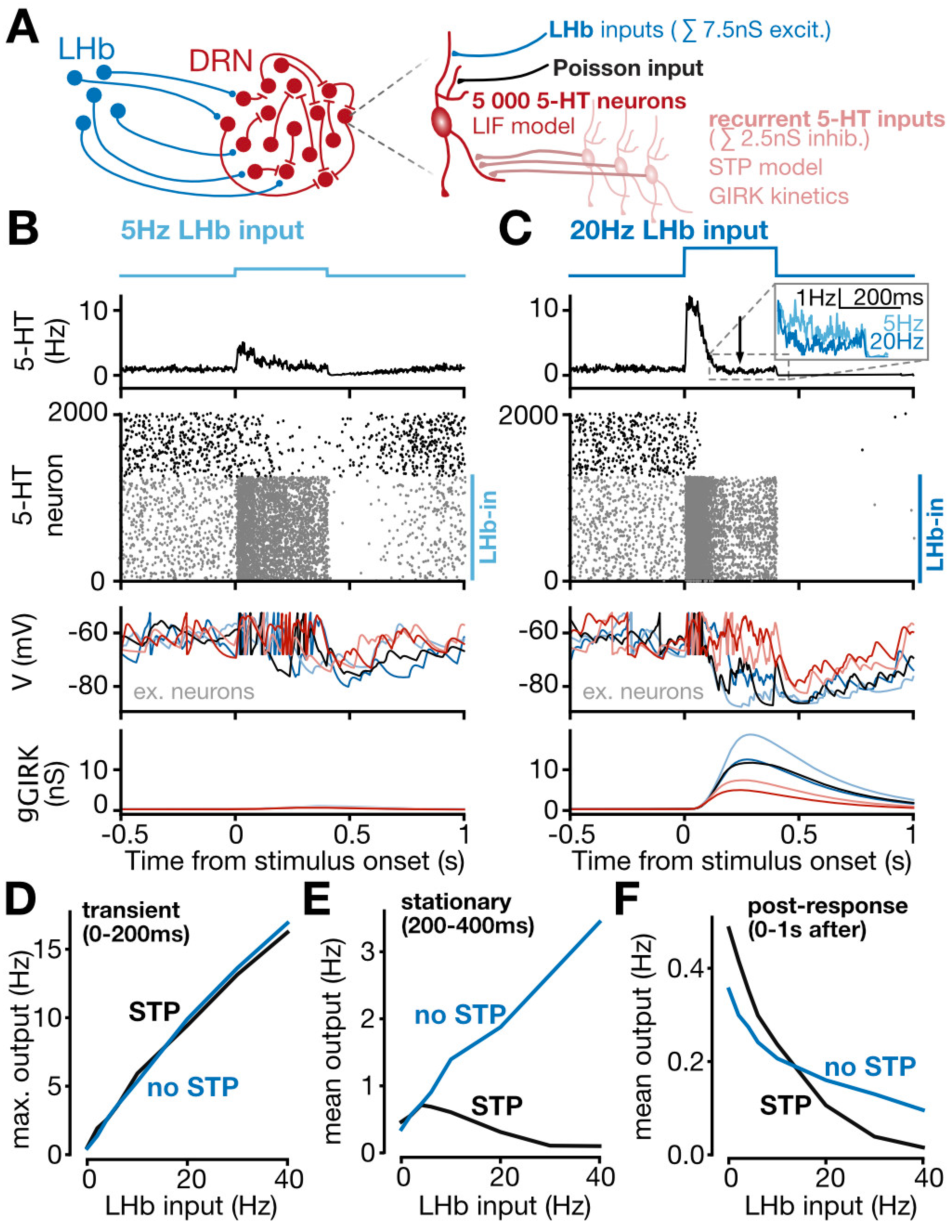
Habenula input to raphe generates a nonmonotonic network response through recurrent short-term plasticity. **(A)** Schematic of the LIF network model depicting direct excitation from LHb (contacting 25% of neurons) and recurrent inhibition between 5-HT neurons. **(B)** Results of a simulation where a 5Hz step input from LHb was delivered to a subset of 5-HT neurons. From top, depicted are the LHb input frequency; the mean firing rates of all 5-HT neurons; a raster plot depicting firing of individual 5-HT neurons over time (grey neurons represent those receiving direct LHb input; black neurons represent those not receiving direct LHb input; raster is limited to the first 2000 of 5000 5-HT neurons for clarity); representative simulated voltage traces of 10 5-HT neurons, each in a distinct color; and representative simulated GIRK conductances from recurrent inhibition of the same 5-HT neurons, each in a distinct color. **(C)** Results of a simulation where a 20Hz step input from LHb was delivered to a subset of 5-HT neurons. Same panels as described in B. **(D)** Input-output transformation for the transient (0-200ms) response phase of all 5-HT neurons in the network, with (black) and without (blue) STP rules at recurrent serotonergic connections. **(E)** Input-output transformation for the stationary (200-400ms) response phase of all 5-HT neurons, with (black) and without (blue) STP rules at recurrent serotonergic connections. **(F)** Input-output transformation for the post-response (0-1s after stimulus) phase of all 5-HT neurons, with (black) and without (blue) STP rules at recurrent serotonergic connections.

First, we considered how recurrence shaped the basic input-output properties of the network. Responses to sustained 5 Hz or 20 Hz LHb activity yielded three periods of the DRN population response (Fig. 6B-C): an early phasic period (0-200 ms), followed by a relaxation to a steady-state response (200-400 ms), and finally a post-stimulus period (up to 1000ms after the end of the stimulus). The early period resulted from 5-HT neurons responding to direct LHb excitation before 5-HT_1A_R -mediated feedback inhibition could modulate the response.

Consequently, the early period output rate was a rectified linear function of the input frequency (Fig. 6D). In contrast, the steady-state response was non-monotonic, displaying an inverted-U shape which peaked at input frequencies around 5 Hz and progressively decreased with higher LHb input frequencies (Fig. 6E). This peculiar non-monotonic function, essentially representing excitation-driven net inhibition, critically depended on the potent short-term facilitation at 5-HT synapses, as *in silico* removal of these short-term dynamics returned a linear, monotonic, input-output function (Fig. 6E). In contrast, the early or post-stimulus period were not appreciably affected by the removal of short-term recurrent dynamics (Fig. 6D-F). This unusual non-monotonic relationship held across a wide range of network connectivity parameters, including changes to recurrent connection probability between 5-HT neurons (Fig S13), and changes to the fraction of 5-HT neurons innervated by long-range input (Fig. S14). The latter manipulation, inspired by previous work describing anatomical heterogeneity within DRN (Zhou et al., 2017; Ren et al., 2018; Dorocic et al., 2014, Weissbourd et al., 2014), unexpectedly demonstrated that the shape and inversion point of this unique input-output relationship was preserved across a wide range of input fractions, from 10% to 50% of 5-HT neurons (Fig. S14A-C). The recurrent drive of the network scaled with the frequency of inputs, but not their precise innervation fraction (Fig. S14D-E). Removing the short-term facilitation dynamics resulted in recurrent drive tracking a combination of input frequency and input innervation (Fig. S14F-J). Together, these simulations suggest that short-term facilitation dynamics uniquely bestows on 5-HT output the ability to track input frequency while normalizing for differences in fine connectivity features. Because the core of this computation emerged largely independently of fine details of connectivity features, it may generalize to other long-range inputs to the DRN.

Next, we considered whether recurrent inhibition, operating over behavioral timescales, could gate interaction between functionally separate submodules or cellular assemblies previously reported within DRN (Ren et al., 2018; Huang et al., 2019; Paquelet et al., 2022). We extended the formalism developed above by implementing two segregated submodules that received excitatory inputs from either LHb (Module 1) or a second generic area termed A (Module 2; Fig. 7A), and asked how input from LHb would disrupt ongoing A-driven activity. Low-frequency (5Hz) LHb input did not appreciably modulate the activity of 5-HT neurons receiving input from area A, as expected by the functional segregation of these submodules (Fig. 7B-C, left). However, high-intensity LHb drive (20 Hz) sharply reduced the activity of A-recipient 5-HT neurons (Fig. 7C-D, right) despite these LHb inputs not directly contacting A-recipient 5-HT neurons. These interactions between submodules again critically depended on the short-term dynamics between 5-HT neurons, as demonstrated by *in silico* removal of these dynamics (Fig. 7E-G). Thus, strong LHb inputs to DRN have the ability to suppress anatomically separate submodules through recurrent inhibition, in essence implementing a winner-take-all computation.

**Figure 7:**
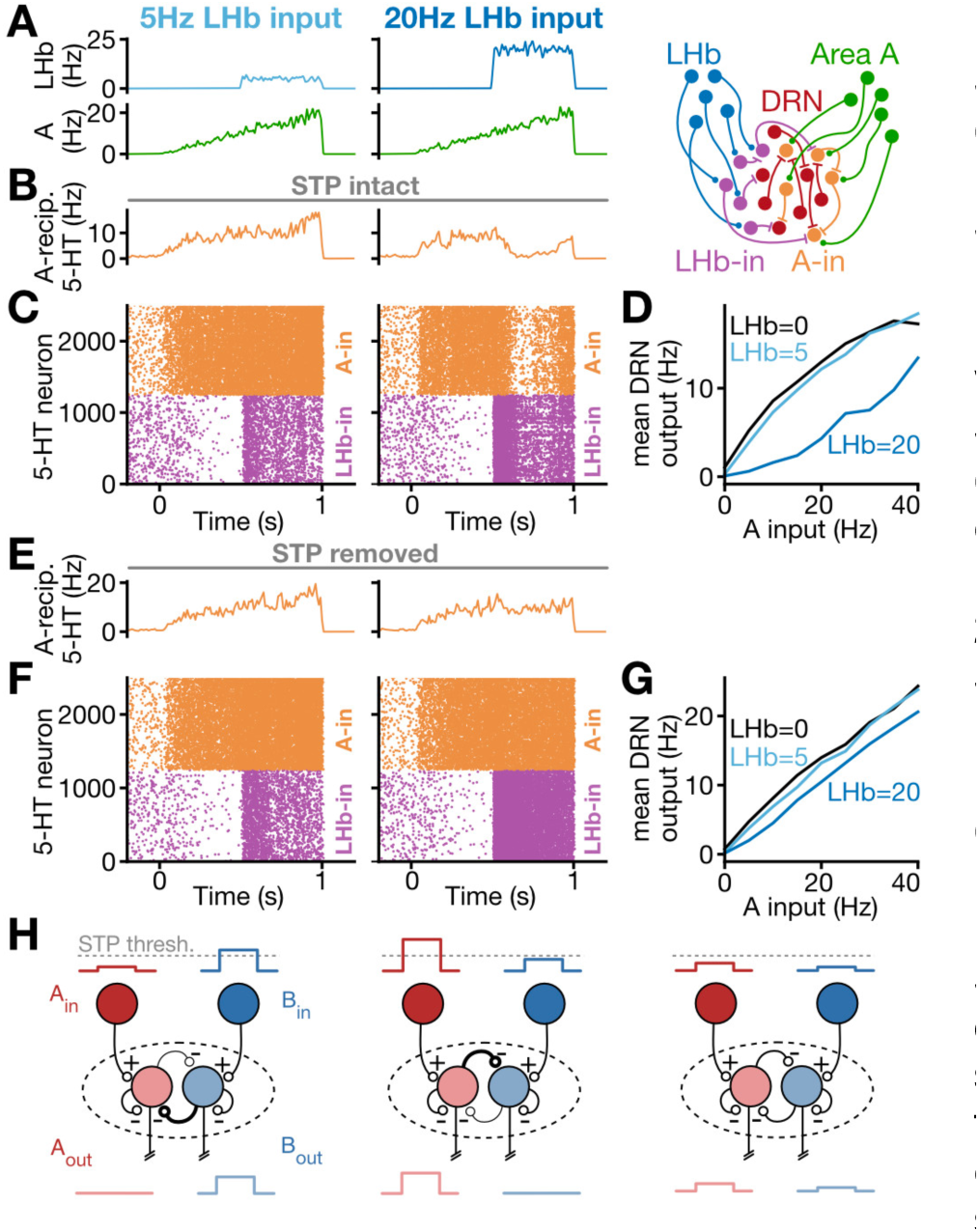
Winner-take-all dynamics generated by recurrent short-term plasticity. **(A)** Simulations with LHb inputs of either 5Hz (left) and 20Hz (middle) were delivered to 5-HT neurons during ramp input from area A. Schematic of LIF network model with two inputs (right). **(B)** Mean responses of 5-HT neurons to 5Hz (left) and 20Hz (right) input with STP rules intact. **(C)** Raster plot showing 5-HT neuron responses to 5Hz (left) and 20Hz (right) input with STP rules intact. **(D)** Input-output function, for input A, with STP rules intact and coincident LHb input at 0, 5 or 20Hz. **(E)** Mean responses of 5-HT neurons to 5Hz (left) and 20Hz (right) input with STP rules removed. **(F)** Raster plot showing 5-HT neuron responses to 5Hz (left) and 20Hz (right) input with STP rules removed. **(G)** Input-output function, for input A, with STP rules removed and coincident LHb input at 0, 5 or 20Hz. **(H)** Schematic of winner-take-all computation with recurrently connected submodules. When one input crosses an STP threshold, output from that submodule dominates while other submodules are suppressed through recurrent inhibition (left and centre). When neither input crosses an STP threshold, outputs from both submodules persist (right).

Further simulations revealed that this winner-take-all computation was not confined to a narrow range of network connectivity parameters. Decreasing the recurrent connection probability tenfold yielded qualitatively similar winner-take-all interactions (Fig. S15). We additionally explored changes in the spatial profile of recurrent connectivity (local vs global), revealing that winner-take-all dynamics were remarkably consistent across different spatial connectivity profiles (Fig. S16). In additional simulations, we explored how spatial overlap of LHb and A inputs within DRN affected winner-take-all dynamics (Fig. S17), demonstrating that strong winner-take-all dynamics emerged even in cases with high input overlap, enhanced by the presence of short-term plasticity at recurrent connections. Finally, to better understand how competing, graded long-range inputs are processed, we quantified the strength of the winner-take-all effect as the input frequency of both LHb and A was varied systematically. This mapping indicated that the winner-take-all effect scaled with the magnitude of both LHb and A over a variety of connectivity parameter choices (Fig. S16E; Fig. S17F), and this scaling depended on the presence of short-term recurrent facilitation (Fig. S16G; Fig. S17I). In summary, the winner-take-all computation between subregions of DRN is present independent of fine connectivity details, and is aided by short-term facilitation of 5-HT release in the DRN.

We next explored the biological plausibility of these winner-take-all rules through additional slice physiology experiments, where either habenula input or local recurrent connections were optogenetically stimulated to gate spiking output. First, a sparse ChETA expression paradigm was employed in DRN as described earlier (Fig. 2; Fig. S18A). In ChETA-negative 5-HT neurons, optical stimulation elicited outward 5-HT_1A_R currents (Fig. S18B). To directly test whether recurrent connections could gate spiking output of (ChETA-negative) 5-HT neurons, we examined the ability of high-frequency (20 Hz) optical stimulation of DRN ChETA-expressing neurons to perturb the persistent spiking induced by direct current injection of 5-HT neurons (Fig. S18C-7D). Consistent with predictions from our computational model, photostimulation triggered a significant reduction in spiking (Fig. S18D-E) which was mediated by 5-HT_1A_Rs (Fig. S18F-L). These data indicate that within DRN, recurrent inhibition can allow populations of 5-HT neurons to significantly shape spiking output of separate 5-HT populations. To ensure that the plausibility of these results extend to activation of long-range synaptic input, we found that high-frequency optical stimulation of LHb afferents in slice reliably inhibited spiking output of 5-HT neurons through 5-HT1ARs (Fig. S19).

Together, these results support a model wherein short-term facilitation at recurrent 5-HT connections generates highly nonlinear network computations. Long-range inputs lead to either excitation at low frequencies, or protracted network suppression at high frequencies; winner-take-all dynamics additionally dominate between functionally distinct submodules at high frequencies. *In silico* manipulations revealed that the short-term dynamics at these recurrent inhibitory connections does not only critically contribute to this core computation, but also to the robustness of these computations to meta-parameter changes in network architecture - possibly allowing these computations to be stable in the face of the heterogeneous connectivity parameters across DRN (described in Fig. S7; Zhou et al., 2017).

### Learned cue-reward associations are conditionally modulated by habenular drive to the DRN

As a final step, we set out to identify a behavioral manifestation of the non-linear computations triggered in the DRN by activating the habenulo-raphe pathway at distinct input frequencies. An internal state of reward anticipation can lead to measurable motor outputs, such as post-cue anticipatory licking in a classical condition task; these licking dynamics are known to be modulated by 5-HT activity (Matias et al., 2017). We thus developed a head-fixed classical conditioning task where we could optogenetically manipulate LHb inputs to DRN while monitoring licking behavior (Fig. 8A-B, see Methods). Mice underwent water deprivation before being trained on an auditory classical conditioning paradigm with three possible reward outcomes (Fig. 8B; 0uL, 5uL, 10uL). Trained mice displayed robust cue-driven anticipatory licking, a conditioned response which indicates expression of a learned stimulus-reward association (Fig. 8C; Cohen et al., 2012; see Methods for training criteria).

**Figure 8:**
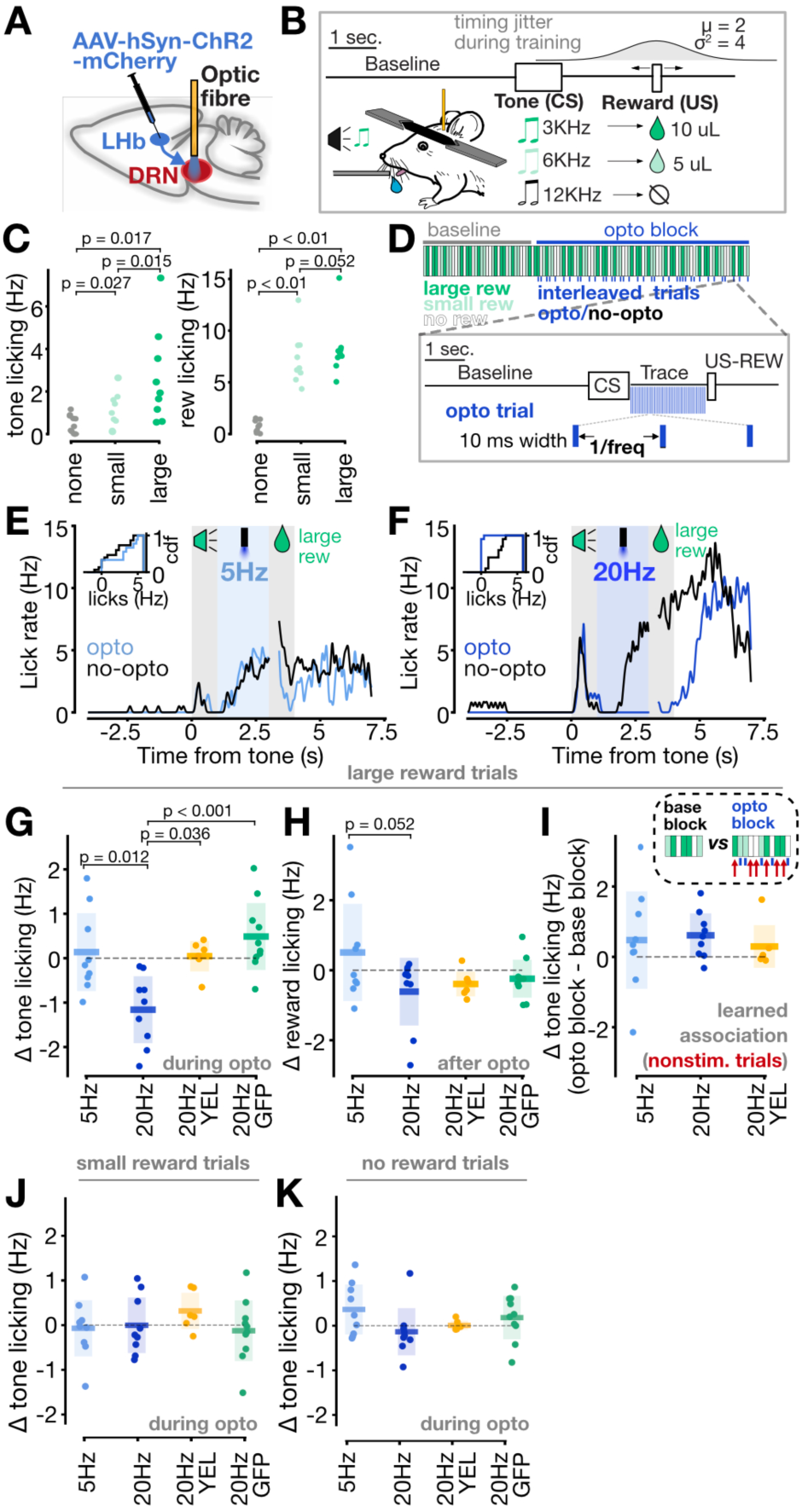
Habenula input to raphe disrupts the expression of learned reward associations in a value- and frequency-dependent manner. **(A)** Schematic illustrating experimental setup (left) and confocal image of LHb injection site (right). **(B)** Schematic of head-fixed classical conditioning task. **(C)** Mice learned to lick in response to the small reward-predicting and large reward-predicting cues (left), and displayed elevated licking after reward delivery (right) (n=9 mice). **(D)** Schematic of trial structure for optogenetic manipulations. **(E)** Mean licking across all large reward trials for a single mouse depicting control trials (black) or 5Hz optogenetic stimulation trials (blue). Inset shows cumulative distribution of licking for all large-reward trials. **(F)** Mean licking across all large reward trials for a single mouse depicting control trials (black) or 20Hz optogenetic stimulation trials (blue). Inset shows cumulative distribution of licking for all large-reward trials. **(G-I)** Summary statistics for optogenetic manipulation during post-tone period of large-reward trials. **(G)** Change in conditioned licking response during tone presentation, between control and optogenetic stimulation trials during opto block (paired two-sided t-tests). **(H)** Mean reward licking change between control and optogenetic stimulation trials (paired two-sided t-tests). **(I)** Change in post-tone licking for nonstimulated trials between baseline block and optogenetic stimulation block. **(J)** Change in post-tone licking between control and optogenetic stimulation trials (small reward trials). **(K)** Change in post-tone licking between control and optogenetic stimulation trials (no reward trials).

We optogenetically stimulated the habenulo-raphe pathway during the post-cue period of interleaved trials, at variable frequencies where physiology and modeling work strongly indicated that direct excitation (low frequency stimulation) or recurrent inhibition (triggered by high frequency stimulation) would dominate within DRN (see Methods). During trials with a large reward outcome, low-frequency (5Hz) optogenetic stimulation had no effect on anticipatory licking, while high-frequency (20Hz) photostimulation induced a striking and rapid decrease in anticipatory licking which recovered rapidly after stimulation ceased (Fig. 8E-G). In contrast, during trials with a small reward outcome, neither 20Hz or 5Hz photostimulation modulated anticipatory licking (Fig. 8J; Fig. S20B-C); during trials with no reward outcome, photostimulating at 5Hz after the auditory tone triggered a slight, but at times clearly noticeable, increase in licking which was not seen during 20Hz photostimulation (Fig. 8K; Fig. S20D-E). As controls for visual artefacts and viral expression, we photostimulated with yellow light in the same cohort (Fig. 8G, yellow; Miyazaki et al., 2018; see Methods), or photostimulated with blue light in a separate cohort with a control viral vector (Fig. 8G; green; see Methods), producing no changes in reward anticipation in either case.

As serotonergic circuits can alter conditioned responses or reward consumption over longer timescales (Matias et al., 2017; Zhong et al., 2017), we asked whether our post-cue optogenetic stimulation had an effect during the subsequent reward period, finding that licking during this post-reward period was not significantly modulated (Fig. 8H). We additionally asked whether post-cue photostimulation drove longer-timescale changes in licking behavior occurring throughout the session. Our analysis, consisting of comparing early trials with nonstimulated late trials occurring during the optogenetic stimulation block, failed to reveal significant long-term alterations in reward anticipation (Fig. 8I; see Methods) consistent with a lack of modulation of the underlying reward association itself. There was a positive correlation between baseline anticipatory licking and optogenetic effect size for high-frequency photostimulation (Fig. S20H), suggesting a divisive rather than subtractive inhibitory effect of pathway activation on reward anticipation. Finally, the effect size of 20Hz and 5Hz stimulation on anticipatory licking showed no relationship (Fig. S20I). To rule out nonspecific stimulation-induced motor effects, we photostimulated at high frequencies during the reward consumption phase instead of in the post-cue period, observing no significant modulation of licking in these conditions (Fig. S20F-G).

Taken together, these results support a model whereby habenular inputs to the raphe can contextually modulate the expression of a conditioned reward response in a temporally precise manner, providing a putative behavioral manifestation of recurrent inhibitory dynamics in this circuit.

## Discussion

Here, using a combination of optogenetic, electrophysiological, computational and behavioral approaches, we provide evidence that DRN is organized in an unsuspected recurrent inhibitory network, characterized by a broad connectivity within this nucleus, and strongly facilitating release. We outlined a core set of computations supported by this connectivity motif, including excitation-driven inhibition of network response at high input frequencies, and a robust winner-take-all computation where anatomically independent raphe subsystems can be antagonistically coupled via recurrent inhibition. These results add to the reported role of 5-HT in modulating cognitive processing, not direct appetitive or aversive proximal behaviors (Fonseca et al., 2015; Miyazaki et al., 2018) and links a local circuit’s network architecture, and associated synaptic dynamics, to a core set of ethologically relevant computations.

We present several lines of experimental evidence supporting a connectivity model whereby 5-HT neurons in the DRN are organized in a local recurrent inhibitory network. Our use of sparse optogenetic stimulation experiments represents in effect an optogenetic equivalent of paired recordings from synaptically-connected neurons, which have provided decisive functional validation of the recurrent connectivity nature of several networks in the brain (Miles & Wong, 1986; Mason et al., 1991; Markram et al., 1997; Debanne et al., 2008). These experiments, unambiguously showing functional connections between 5-HT neurons, supports a model which is largely inconsistent with the widely and long-held idea that 5-HT neurons are regulated in an autocrine fashion by dendritically-released 5-HT acting on so-called 5-HT_1A_R somatodendritic autoreceptors (Gozlan et al., 1983; Weissmann-Nanopoulos et al., 1985; Pineyro & Blier, 1999). This autoreceptor hypothesis makes several experimental predictions which we directly tested and failed to find support for. While our results do not formally allow to distinguish whether interconnected 5-HT neurons communicate through dendro-dendritic or canonical axo-dendritic functional contacts, or whether release of 5-HT is spatially confined or consists of more spatially diffuse volume transmission (Maxwell et al., 1983; Ridet et al., 1994; Bunin & Wightmann, 1998), we nonetheless note that classic axonal release provides the most parsimonious and credible model accounting for our results. While a subset of 5-HT dendrites contains 5-HT containing vesicles (Descarries et al., 1982; Chazal and Ralston, 1987), 5-HT containing axon terminals are nonetheless widespread in the DRN and are structurally indistinguishable from those found elsewhere in the brain (Chazal and Ralston, 1987). Importantly, this view does not preclude the presence of NMDAR-dependent dendritic release of 5-HT that may operate in distinct physiological conditions (Colgan et al., 2012). Furthermore, the facilitation of the short-term 5-HT release stands in stark contrast to the robustly depressing dopamine release dynamics recently described in striatum (Liu et al., 2018, Liu et al., 2021).

Previous experimental and theoretical work has identified a number of possible computational functions of network recurrence, including winner-take-all competition (Coultrip et al., 1992), signal amplification and selectivity (Douglas et al., 1995) and probabilistic inference (Echeveste et al., 2020). Our results show that the recurrent inhibitory dynamics lead to a non-monotonic input-output relationship in the DRN by supporting a strong excitation-driven inhibition, where larger excitatory inputs generate greater network suppression. A natural question is how this computation compares with gain control associated with classical feedforward inhibition motif. A range of distinct gain control regimes have been described with feedforward inhibition such as subtractive and divisive inhibition (Ayaz & Chance, 2009) and even non-monotonic inhibition (Mejias et al., 2014; Lewis et al., 2007). However, the recurrent inhibition motif described here is unique in that the phasic network response is dominated by direct excitation and linearly reports input intensity, while the delayed steady-state response exhibits non-monotonic behavior with increasing excitation. In principle, these temporally distinguishable epochs of 5-HT output dynamics could support multiplexed codes such as valence encoding over multiple timescales (Cohen et al., 2015). Our network model additionally uncovered an emergent winner-take-all-like computation in the DRN, where strong activation of connected modules of 5-HT neurons can inhibit activity of others. This builds on previous work describing anatomically segregated DRN subdomains (Ren et al., 2018), and implies that a winner-take-all computation governs state selection between submodules. While these simulations uncovered general algorithmic principles which were tested with a range of connectivity profiles in our network models, more precise quantitative information on the recurrent connectivity between submodules (Ren et al., 2018) or on the additional recruitment of feedforward GABAergic inhibition by some inputs (Zhou et al., 2017; Geddes et al., 2016) should bring about refinement of these general principles.

More broadly, our behavioral results provide evidence that LHb inputs to the DRN can rapidly and reversibly alter behavioral state, which is consistent with previous findings on the habenulo-raphe pathway in other model organisms (Amo et al., 2014; Marquez et al., 2020) and may represent an important role for this pathway in dynamic state selection. Indeed, cue-reward learning has been shown to lead to a gradual increase in the population activity of 5-HT neurons during reward anticipation, peaking around reward delivery (Zhong et al., 2017), and current theories postulate that 5-HT neurons represent valence information during associative learning (Liu et al., 2017). Thus, the DRN microcircuit has many of the core ingredients necessary to perform such contextual behavioral state transitions.

Animals often exhibit sharp thresholds when transitioning between ethologically relevant behavioral states such as exploitation and exploration (Marques et al., 2020). These state transitions require brain circuits which continually evaluate graded representations of either internal or external cues, enacting thresholding operations on behavioral timescales. Building on recent theories (Liu et al., 2020; Luo et al., 2016; Cohen et al., 2015; Marques et al., 2020), the winner-take-all computation between ensembles of recurrently connected 5-HT neurons, operating over protracted timescales, satisfies many of the conditions required for the largely exclusive, non-overlapping implementation of distinct naturalistic behaviors (Ren et al., 2018).

## Acknowledgements

This work was supported by grants from the Canadian Institute for Health Research (JCB; RN), the uOttawa Brain and Mind Institute and the Canadian Heart and Stroke Foundation (JCB), Brain Canada (JCB), the Krembil Foundation (JCB) and the Canadian Foundation for Innovation (JCB). We thank members of the Béïque, Naud and Maler lab for helpful discussions. We would also like to thank Drs Salah El Mestikawy and Sylvain Williams for providing us with the SERT-Cre mouse line.

## Author contributions

Conceptualization, M.B.L., S.G., S.H-D., R.N., and J.-C.B.; Methodology, M.B.L., S.G. ; Software, M.B.L.; Formal Analysis, M.B.L.; Investigation, M.B.L., S.G., M.C., E.H., S.M., E.H.-G., ; Resources, M.B.L., E.H. and R.N. ; Writing – Original Draft, M.B.L.; Writing – Review & Editing, M.B.L., R.N., and J.-C.B.; Supervision, J.-C.B.; Funding Acquisition, J.-C.B.

## Data availability statement

The datasets and code generated during this study are available from the corresponding author upon reasonable request.

## Methods

### Viral Constructs and Retrograde Tracer

| Construct | Source | Titre (IU/μl) |
| --- | --- | --- |
| AAV9.CAG.hChR2(H134R)-mCherry.WPRE.SV40 | (Penn Vector Core, AV-9-20938M) | $\sim 10^{12}$ |
| AAV9.Ef1a.DIO.ChETA(E123T/H134R)-eYFP.WPRE.hGH | (Penn Vector Core, AV-9-26968P) | $\sim 10^{12}$ |
| rAAV2-retro-hSyn1-cre | (Janelia Farm Research Campus) | $\sim 10^{13}$ |
| AAV2/9.emCBA.eGFP | Canadian Neuor Photonics Platform, Université Laval | $\sim 10^{13}$ |
| AAV-EF1a-DIO-5-HT2M (Serotype: 2/9) | Canadian Neuor Photonics Platform, Université Laval | $\sim 10^{12}$ |

#### Animal use

All experiments and procedures were performed in accordance with approved procedures and guidelines set forth by the University of Ottawa Animal Care and Veterinary Services. For this study, SERT-Cre mice (B6.Cg-Tg(Slc6a4-cre)ET33Gsat/Mmcd) were crossed with the R26R line (Rosa26R-tdTomato fluorescent protein; tdTom) to generate a SERT-Cre::Rosa-tdTomato mouse line (referred to in the manuscript as SERT-Cre::tdTom), and SOM-Cre mice tm2.1(cre)Zjh/J) were crossed with the R26R line to generate a SOM-Cre::Rosa-tdTom mouse line (referred to in the manuscript as SOM-Cre::tdTom). Mice were kept on a 12:12 hour light/dark cycle, with access to food and water ad libitum unless otherwise noted. A mix of male and female SERT-Cre::tdTom and SOM-Cre::tdTom mice (1-3 months for slice physiology experiments, 2-6 months for behavior experiments) were used. A mix of male and female SOM-Cre::Rosa-tdTom mice were used (1-3 months for slice physiology experiments).

For the behavior portion of the study, mice with a C57BL/6 background were utilized, either originating as mice with a wild-type genotype originating from the SERT-Cre:: tdTomato line, or as mice purchased from Charles River Labs. Mice were kept on a 12:12 hour reverse light/dark cycle and were water restricted to 85-90% of their initial weight during behavioral testing.

### Slice physiology and anatomy

#### Stereotaxic Injections

All experiments and procedures were performed in accordance with approved procedures and guidelines set forth by the University of Ottawa Animal Care and Veterinary Services. For stereotaxic injections, mice were injected with 0.05 mg/kg buprenorphine and anesthetized by inhalation of isoflurane. Injections were performed using a 10 μl Hamilton syringe with a 33 gauge needle, or with a Nanoject III programmable injector with a pulled glass micropipette. 0.4 - 0.5 μl was injected per coordinate, injected at a rate of 0.1 – 0.2 ul per minute.

Stereotaxic co-ordinates are as follows: LHb (from Bregma skull landmark, in mm: AP −1.7, ML +/- 0.4, DV −3.0) and DRN (from Lambda skull landmark, in mm, 20 degree angle: AP −0.5, ML +1.0, DV −3.0; or 0 degree angle: AP −0.5, ML +0, DV-3.2, −3, −2.8, sequential injections to tile DV extent of DRN with total volume split between all 3 coordinates. Both injection types were utilized). For retrograde tracing, 0.4 μl of rAAV-retro-TdTomato was injected. For ChR2 (H134R) expression in LHb, 0.4 μl of AAV9.CAG.hChR2(H134R)-mCherry.WPRE.SV40 (titre ∼10^12^ GC/mL) was injected bilaterally and animals were left to recover for 3 to 4 weeks prior to ex-vivo electrophysiological recordings. In anterograde tracing, AAV-GFP was injected in the LHb and animals were perfused 2-3 weeks post-injections.

For experiments requiring sparsely labeled cells in the DRN with ChETA-YFP, we further diluted virus in a 20 % mannitol/saline solution at a ratio of 2:1 (mannitol solution: virus solution). In these experiments, the virus solution was AAV9.Ef1a.DIO.ChETA(E123T/H134R)- eYFP.WPRE.hGH. The mixed mannitol and virus solution was then injected in the DRN at the coordinates above.

For experiments involving expression of ChETA in SOM interneurons in DRN, utilizing the SOM-Cre::Rosa-tdTom mouse line, we injected a total of 0.5uL of AAV9.Ef1a.DIO.ChETA(E123T/H134R)-eYFP.WPRE.hGH at the DRN coordinates above.

#### Slice preparation

LHb and DRN containing brain slices were prepared from 8 to 10 week old mice (3 to 4 weeks post-stereotaxic injections). For electrical stimulation experiments, brainstem slices were prepared from P21-P30 mice. Acute slices were prepared similarly to previously described methods (Geddes et al., 2016). In brief, mice were anesthetized by isoflurane inhalation (Baxter Corporation, Canada) and sacrificed by decapitation. The brain was removed, and brain slices were sectioned while immersed in ice-cold choline chloride-based cutting solution of the following composition (in mM): 119 choline-Cl, 2.5 KCl, 1 CaCl2, 4.3 MgSO4-7H2O, 1 NaH2PO4, 1.3 sodium L-ascorbate, 26.2 NaHCO3, and 11 glucose, and equilibrated with 95% O2, 5% CO2. Slices were recovered in a chamber containing standard Ringer’s solution of the following composition (in mM): 119 NaCl, 2.5 CaCl2, 1.3 MgSO4-7H2O, 1 NaH2PO4, 26.2 NaHCO3, and 11 glucose, at a temperature of 37°C, continuously bubbled with a mixture of 95% O2, 5% CO2. Slices were left to recover for 1 hour in a recovery chamber and equilibrate to a temperature of approximately 21°C until recordings were performed.

#### Whole-Cell Electrophysiology

DRN neurons were visualized using an upright microscope: (1) Examiner D1 equipped with Dodt-contrast or differential-interference contrast (DIC) (Zeiss, Oberkochen, Germany; 40x/0.75NA objective) or, (2) BX51W1 equipped with DIC optics (Olympus, Center Valley, PA; 40x/0.80NA objective). 5-HT neurons were visually identified by TdTomato signal driven by the SERT promoter under EPI-fluorescence illumination (xCite Series 120pc). Whole-cell recordings were carried out using an Axon Multiclamp 700B amplifier and electrical signals were low-pass filtered at 2 kHz and sampled at 10 kHz with an Axon Digidata 1440A (or 1550) digitizer. Whole-cell recordings were performed using borosilicate glass patch electrodes (3-6 MΩ; World Precision Instruments, Florida) pulled on a Narishige PC-10 pipette puller (Narishige, Japan). Slices were constantly bathed in Ringer solution containing (in mM): 119 NaCl, 2.5 KCl, 1.3 MgSO4, 2.5 CaCl2, 1.0 NaH2PO4, 11 glucose, 26.2 NaHCO3, and 0.1 L-Tryptophan saturated with 95% O2 and 5% CO2 (pH = 7.3; 295-310 mOsm/L). In the experiments described in Fig. S4 (exploration of feedforward inhibition) and Fig. S10 (electrical stimulation of 5-HT afferents, and optogenetic stimulation of long-range LHB input to 5-HT neurons), the Ringer solution was identical to that described above except that 1.5mM CaCl2 was used in place of 2.5mM CaCl2 (as described in the caption).

Three intracellular recording solutions were used. For voltage-clamp experiments involving solely excitatory post-synaptic currents (EPSCs) recordings, we used an intracellular solution of the following composition: 115 mM cesium methane-sulfonate, 5 mM tetraethylammonium-Cl, 10 mM sodium phosphocreatine, 20 mM HEPES, 2.8 mM NaCl, 5 mM QX-314, 0.4 mM EGTA, 3 mM ATP(Mg2+), and 0.5 mM GTP, pH 7.25 (adjusted with CsOH; osmolarity, 280–290 mOsmol/L). For recordings involving 5HT1AR-mediated current or potential, we used the either of the following internal solutions: 1) 115 mM potassium gluconate, 20 mM KCl, 10 mM sodium phosphocreatine, 10 mM HEPES, 4 mM ATP(Mg2+), and 0.5 mM GTP, pH 7.25 (adjusted with KOH; osmolarity, 280–290 mOsmol/L): or 2) 115 mM KMeSO4; 5 mM KCl; 5 mM NaCl; 1 mM MgCl2; 0.020 mM EGTA; 10mM HEPES; 4 mM ATP; 0.4 mM GTP; and 10mM Na2phoshocreatine, pH7.4 (adjusted with KOH; osmolarity 280-290 mOsmol/L). For all voltage clamp recordings, access resistance was continuously monitored using a 125 ms, 2 mV hyperpolarizing pulse, at least 245 ms prior to electrical or optogenetic stimulation. Recordings were discarded when the access resistance changed by >30%. Liquid junction potential was not compensated for.

The slice physiology experiments described in Figures 1 and 2 were conducted at room temperature (20-24C). The electrical stimulation experiments described in Figures 3 and 4, as well as Figures S4 (exploration of feedforward inhibition) and S10 (replication of main results of Figure 3), were conducted nominally at 30-32°C through the use of an in-line bath heater and stage heater.

#### LED Photostimulation

LED photostimulation was delivered using a PlexBright LED module and controller (465 nm; Plexon, Texas). Light was delivered through a 200 μm patch fiber cable with a bare fiber tip with ∼ 1 cm of glass exposed. This fiber was held at a ∼45 degree angle to the slice, with the tip at a distance of approximately 1cm from the surface of the acute brain slice. The tip of the fiber emitted light in a narrow, approximately conical shape with a measured irradiance of 792 mW/mm^2^ at the slice. The fiber’s position was manually calibrated such that all of the emitted light was hitting the slice, focused on the portion of the slice where the physiologically probed neuron was. For EPSC/EPSP stimulation, unless otherwise stated, optically evoked-PSCs were stimulated at a frequency of 0.1 or 0.067 Hz with light pulses of 1-3 ms in length and an irradiance of 792 mW/mm2 measured. In driving 5-HT1AR-mediated currents, optical stimulation was accomplished using 10 ms pulses at 20 Hz for 400 ms.

#### Electrical stimulation of 5-HT afferents

Electrical stimuli were delivered through a bipolar concentric stimulating electrode located >100 um from the soma of the recorded neuron. Stimuli were 0.1 ms in duration, and consisted of frequencies provided in the text. For all electrical stimulation experiments, we bath-applied a cocktail of synaptic blockers to isolate the 5-HT1AR-mediated responses (i.e., 10 µM NBQX, 100 µM picrotoxin, 200 nM CGP 55845). All 5-HT electrical stimulation experiments were performed at near-physiological temperature (30-32 °C).

#### Immunohistochemistry

Mice were perfused using a minimally invasive method developed by Kenneth D. Eichenbaum and colleague in 2005. Briefly, mice were put under general anaesthesia with 1.5% isoflurane. The animal was then placed on its back for injection (1 mL PFA4% in 0.1 M PBS). The injection was done with a 1 mL syringes equipped with a 26G needle over a 30 seconds course. The correct position of the needle was confirmed by the appearance of blood in the needle hub. Brains were then put in 10 mL of PFA4% overnight and then transferred in a 15 mL tube with 10 mL of PBS (0.1 M PBS, 0.1% Sodium azide). The fixed brains were then cut in 50 um thick slices with a vibrotome (Vibrotome,1500 sectioning system).

Free-floating sections were blocked in 1% BSA + 0.03% Triton X-100 (TX-100) for 45-60 min, and incubated in primary antibody in blocking solution at 4 °C. On the following morning, slices were washed three times, for 5 min each time, with 0.1% Tx-100 before incubation in the secondary antibody goat anti-rabbit–conjugated Alexa-647 (Life technologies, A21245) for 2 h at room temperature. Slices were washed with 0.1% Tx-100 three times, for 5 min each time, before mounting onto slides with Fluoromount-G (Southern Biothech, 0100-01). The primary antibody used for this paper were Tryptophan hydroxylase 2 (1:1,000; Millipore, ABN60), NeuN (1:1000; EnCor Biotechnology, RPCA-Fox3-AP) and Nissl (1:1000; NeuroTrace® 435/455 Blue Fluorescent Nissl Stain, Life technologies, N21479).

#### Fluorescent tracer injections in DRN

To examine the local connectivity and network structure in DRN, we performed visually guided injections of fluorescent tracer in DRN in acute slices (described fully in Trinh et al., 2015). Briefly, in these experiments, SERT-CRE-tdTomato animals (P30-P40) were sacrificed and acute slices prepared as described above (*slice preparation*). We utilized the following fluorescent dextrans interchangeably, observing similar expression and transport between them: Cascade Blue; Alexa Fluor 488; Alexa Fluor 647 (all suspended at 10 mg/mL in 0.1M phosphate-buffered saline (PBS) pH 7.2). At 5x-10x magnification with a Zeiss upright microscope with an epifluorescent lamp, fluorescent dextran (described above) was injected in either dorsal DRN or ventral DRN (all locations given in Fig. S7D, inset) by iontophoresis (five to ten 400 ms pulses of ∼ –100 V via a 7 mOhm electrode). After a fluorescent sphere of ∼100 µm diameter was observed, injection was stopped. Slices were incubated overnight in oxygenated ACSF to allow anterograde and retrograde transport, fixed in 4% paraformaldehyde and rendered transparent using the SeeDB procedure (see Trinh et al., 2015 for details). Slices were imaged with a Zeiss confocal microscope (LSM800 AxioObserver Z1 microscope; Zen software) with the appropriate filters for resolving the utilized fluorescent dextrans.

#### Data Analysis

All electrophysiological recordings were analyzed offline using Clampfit 10.4 (Molecular Devices), Origin 8.5 software (OriginLab) and the Numpy and Scipy technical computing libraries within Python. All data are presented as means +/- SEM. n refers to the number of cells recorded. Unless otherwise stated, statistical significances were derived using a standard Student’s two-sided t-test (p < 0.05). P values of less than 0.05 were considered statistically significant and are indicated with an asterisk(*). Unless otherwise stated, all measurements were taken from distinct neurons.

All anatomy experiments (fluorescent tracer injections, described above) were analyzed using a combination of IMARIS and custom Python scripts. After acquiring images (described above in *Fluorescent tracer injections in DRN*), we performed semi-automated cell identification using the IMARIS image analysis software package (Oxford Instruments), employing the spots function. This allowed us to identify neurons dual-labeled with tdTomato and the fluorescent dextran dye, and extract their coordinates in three dimensions (X,Y,Z) relative to the centre of the injection. Only neurons that were clearly outside of the extent of the dextran injection sphere were considered. We additionally extracted the locations of all neurons that were tdTomato positive (using Imaris) to act as a null distribution to compare with the co-expressing neurons. The validity of all identified neurons was manually checked against the raw fluorescence images. Together, this analysis yielded, for each individual dextran injection, a dataset of 3-dimensional coordinates for dual-labeled (tdTomato and fluorescence dextran) 5-HT neurons and single-labeled (tdTomato) 5-HT neurons.

Custom Python scripts employing the Numpy and Scipy technical computing libraries were utilized to perform analysis on this dataset. To compare the observed probability distribution over distance of dual-labeled neurons with the null model of unstructured connectivity, we first constructed a probability distribution of all single-labeled (tdTomato) 5-HT neurons over distance. We computed the difference, at each distance bin, between the probability distribution for dual-labeled neurons and the probability distribution for single-labeled neurons, yielding a delta probability distribution over distance. To construct a polar plot, we computed the angle of each cell relative to the injection centre and computed a probability distribution over these angles, calculating a delta probability distribution over angle similar to above.

#### Drugs and Chemicals

Picrotoxin, 2,3-Dioxo-6-nitro-1,2,3,4-tetrahydrobenzo[f]quinoxaline-7-sulfonamide di-sodium salt (NBQX), DL-2-Amino-5-phosphonopentanoic acid (D,L-APV), picrotoxin, CGP 55845 and WAY 100-635 were purchased from Abcam (Cambridge, MA). Tetrodotoxin (TTX) was purchased from Tocris Bioscience (Ellisville, MO). 4-Aminopuridine (4-AP) was purchased from Sigma Aldrich (St. Louis, MO).

### Two-photon imaging of a serotonin biosensor

#### Stereotaxic Injections

Mice were injected with 0.05 mg/kg buprenorphine and anesthetized by inhalation of isoflurane. Injections were performed using a Nanoject III programmable injector with a pulled glass micropipette. 0.4μl was injected at a rate of 0.1μl per minute. Stereotaxic co-ordinates are as follows: DRN (from Lambda skull landmark, in mm, 0 degree needle angle: AP +0.4, ML +0, DV-3.2, −3, −2.8, sequential injections to tile DV extent of DRN with total volume split between all 3 coordinates.) 0.4 μl of AAV-EF1a-DIO-5-HT2M (Serotype:2/9; titre ∼4.3*10^12^ GC/mL) was injected and animals were left to recover for 3 to 4 weeks prior to 2P imaging experiments.

#### Two-photon imaging

Brain slices were prepared using the method described above in *Slice physiology and anatomy,* subsection *Slice preparation.* Slices were positioned in a recording chamber for imaging, and received constant bath application of Ringer solution containing (in mM): 119 NaCl, 2.5 KCl, 1.3 MgSO4, 2.0 CaCl2, 1.0 NaH2PO4, 11 glucose, 26.2 NaHCO3, and 0.1 L-Tryptophan saturated with 95% O2 and 5% CO2 (pH = 7.3; 295-310 mOsm/L). All imaging was conducted at a measured temperature of 30-32C through the use of an inline heater and stage heater. Two-photon imaging was performed using an ultrafast Ti:Sapphire pulsed laser (InSightX3, Spectra Physics, Milpitas, California) tuned to 920nm to visualize both the GRAB5-HT2M (virally expressed) and tdTomato (5-HT neuron marker) signal. Laser intensity was controlled using an acousto-optic modulator coupled to a galvanometer scanning system (Zeiss 810; Germany). Images were acquired using a 40x/0.8NA objective.

#### Electrical stimulation of 5-HT afferents

Electrical stimuli were delivered through a patch pipette filled with Ringer solution (described above) located within the field of view. The pipette was filled with Alexa 594 for visualization during imaging experiments. Stimuli were 0.1 ms in duration, and consisted of frequencies provided in the text.

#### Drugs and chemicals

In some experiments, we bath applied either serotonin hydrochloride (Tocris Bioscience (Ellisville, MO; 50mM) or TTX (Tocris Bioscience (Ellisville, MO), 0.5uM).

#### Data analysis

ROIs were manually annotated and raw fluorescence was extracted using Suite2P (Pachitariu et al., 2016). All analysis was performed with custom Python scripts employing the Numpy and Scipy technical computing libraries. To compute *ΔF*/*F*_0_traces, *F*_0_was calculated as the mean fluorescence from the last three pre-stimulus images. To compute ∫ *ΔF*/*F*_0_, in order to capture the full serotonin biosensor response, we integrated the trial-averaged *ΔF*/*F*_0_ response from stimulus onset to 5 seconds post-stimulus. To compute *max*(*ΔF*/*F*_0_), we computed the value of the trial-averaged *ΔF*/*F*_0_ response for the same time period as described above.

For full-frame analysis of GRAB sensor responses, to decrease noise, we adopted a spatial pooling approach, first downsampling each 512×512 frame of GRAB sensor responses by a factor of 32 to yield a resolution of 16×16 pixels. Next, for each of these spatially pooled pixels, we computed a *ΔF*/*F*_0_value at each time point as described above for ROI analysis. This yielded an image describing the spatial layout of stimulus responses for each stimulus type. To compute the pixel response index, for each frame we first considered the distribution of the ∫ *ΔF*/*F*_0_ responses across all pixels for the 5Hz electrical stimulation condition. From this 5Hz distribution we computed a standard deviation, *σ*_5*Hz*_, and mean, *μ*_5*Hz*_. These 5Hz parameters were used to calculate a normalized Z-score metric for the other conditions: ***z*** = (***r*** − *μ*_5*Hz*_)/*σ*_5*Hz*_, where ***r*** is a vector across all pixels of ∫ *ΔF*/*F*_0_ values for a particular electrical stimulation frequency.

We computed the spatial autocorrelation of these images using the signal.correlate2d function from the Scipy technical computing library, yielding a 2-dimensional autocorrelation function of the same shape as the original stimulus response image, with maximal intensity value in the centre.

To compute spatial masks associated with the 2-dimensional autocorrelation function, we randomly shuffled the spatial location of each response pixel and computed a 2-dimensional autocorrelation function on this shuffled image. We repeated this shuffling 100 times for each image to compute a distribution of null autocorrelation responses across all pixels (excluding the central pixel corresponding to the two images fully aligned). Utilizing this null distribution of autocorrelation values, we assigned a threshold of p=0.0001 for which experimental autocorrelation values would have to exceed to be considered significant (ie greater than 99.99% of null autocorrelation values). The number and spatial location of these statistically significant autocorrelation values were used to compute the spatial mask and area of autocorrelation.

The pixel correlation across trials was computed by calculating the Pearson correlation coefficient across all pixels for each unique pair of trials (typically 4-6 trials were presented), and then taking the mean of this Pearson r value. The pixel correlation (GRAB vs tdTomato) was computed similarly, but using the GRAB sensor response and the tdTomato signal as input.

### Modeling of synaptic dynamics and network simulations

#### GIRK conductance

The GIRK conductance kinetics were modelled as a sum of exponentials:

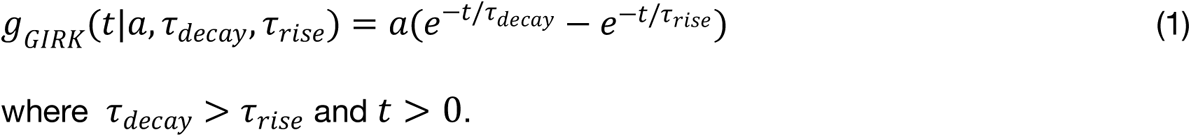

The parameters *a*, *τ*_*decay*_, *τ*_*rise*_ were fit to experimentally-determined 5-HT1AR-mediated GIRK conductances obtained following single-pulse electrical stimulation data (n=6 neurons) using a least-squares approach as follows: We first fit a monoexponential decay function *f*(*t*|*τ*_*decay*_) = *e*^−*t*/*τ_decay_*^ to the normalized post-peak decay phase of 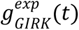. This yielded a constrained estimate of the GIRK decay, *τ*_*decay*_. Next, fixing *τ*_*decay*_, we found the values of *a*, *τ*_*rise*_ that minimized the mean-squared error over the whole observation period. Each estimate was averaged across the population (n=6 neurons):

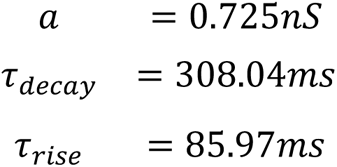

#### Quantifying 5-HT release dynamics through optogenetic stimulation

In some experiments, we optogenetically stimulated CHETA-expressing 5-HT neurons in slice, while recording 5-HT1AR outward currents from identified 5-HT neurons. By photostimulating at 20Hz and varying the optical pulse number, we could infer basic properties of release dynamics at these 5-HT releasing synapses.

We assume that the current generated by a single pulse is a random variable following a binomial distribution *B*(*n*, *p*), which reflects the activity of *n* independent 5-HT releasing synapses with release probability *p*. For a train of successive pulses, we assume *n* is fixed, and that each pulse is independent. Then *Q*(*n*_*pulse*_ = *a*), the distribution of currents given by *a* pulses, is distributed as a sum of *a* binomial distributions:

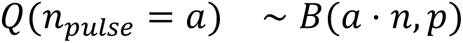

Accordingly, the variance of Q as the pulse number *a* varies is as follows (Sigworth, 1980):

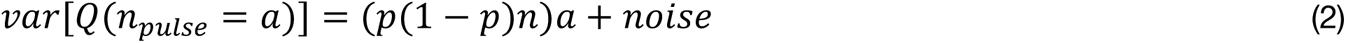

Since we observed that the variance of the current did not change as a function of pulse number (Fig. S6), we conclude that the release probability of 5-HT in these conditions can be approximated as being deterministic (i.e., close to *p* = 1). This conclusion nonetheless probably reflects the artefactual increase in release probability obtained when optogenetically activating synaptic terminals (Zhang & Oertner, 2007; see the section below).

#### Quantifying 5-HT release dynamics through electrical stimulation

We performed experiments where 5-HT afferents were electrically stimulated in acute slice preparations while recording from identified 5-HT neurons. Electrical stimulation intensities were low (in the perithreshold response range), and stimulation frequency was 1Hz. To quantify successes and failures, we computed the total event charge triggered by each stimuli (by integrating current over time), which formed an *event* distribution containing both putative successes and failures. We constructed a surrogate *noise* distribution by performing an identical analysis on a nonstimulated portion of the same recording for each neuron (*i.e.,* integrating equivalent durations of membrane noise where no stimuli were presented). Next, for each neuron we transformed the *event* and *noise* charge distributions into z-scored distributions (where the z-score was computed from that neuron’s *noise* distribution). Finally, we pooled z-scored distributions across neurons. This allows us to directly compare electrically evoked responses; responses with a z-score > 3 likely represent successes, while responses with a z-score near 0 probably represent release failures. For electrical stimulation at higher frequencies (1-20Hz), we constructed surrogate noise distributions and event distributions in a manner identical to that described above.

For paired-pulse and pulse-chase stimuli, we quantified responses by first computing total charge elicited over the duration of each of the stimuli (integrating current over time). We next computed the total charge elicited by a single electrical stimulation delivered in isolation, and used this to calculate an expected value for purely linear summation of two pulses (*i.e.,* by multiplying the single stimulation-induced charge by 2). Finally, for each neuron, we normalized the stimulus-evoked charges in the paired-pulse or pulse-chase condition by the expected value for linear summation of two individual pulses.

#### Short-term plasticity

Short-term plasticity dynamics were modeled using a linear-nonlinear approach called the spike-response plasticity (SRP; Rossbroich et al., 2021). It utilizes a possibly complex efficacy kernel to capture time-varying, spike-triggered changes in release probability (i.e., short-term plasticity). Namely, each presynaptic spike is convolved with the following efficacy kernel:

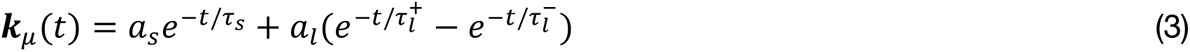

To capture the complexity of the dynamics, we chose to implement a kernel consisting of multiple timescales of responses. The short component is controlled by *τ*_*s*_ and *a*_*s*_, and dictates the rapid short-term facilitation dynamics. The long component is controlled by 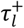 and *a_l_*, and dictates the more prolonged short-term depression dynamics. 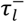 is a shaping term that reduces the effect of short-term depression in the early post-spike phases where short-term facilitation dominates. Since the kernel consists of a sum of exponential functions, it can be conveniently written as a dynamical system. Let *k* reflects the current efficacy state of the synapse (*k* = ***k***_*μ*_ ∗ *S*) and is transformed through a nonlinear operation into a spike efficacy, *μ*. In turn, *μ* dictates the magnitude of the postsynaptic response given a presynaptic spike, and is implemented as a nonlinear readout of k:

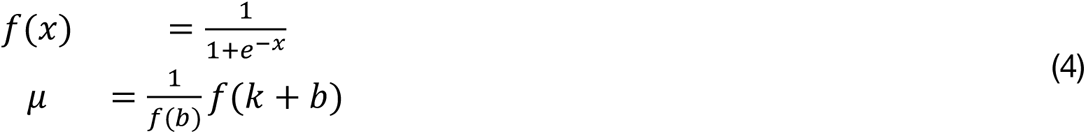

Here, *f*(*x*) is the nonlinear function and *b* is a baseline term that controls whether the STP dynamics evolve sublinearly or supralinearly.

#### Fitting short-term plasticity dynamics to experimental data

We start with a set of paired-pulse electrical stimulation experiments where the time interval ranges from 50ms to 1sec. For our experimental dataset, we had computed, for each time interval *t*_*i*_, a ratio of the total charge transfer for both pulses *q*_1_ and *q*_2_ to the charge transfer for a single pulse delivered separately:

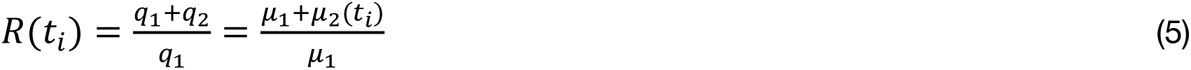

Where *μ*_1_ and *μ*_2_ relative postsynaptic response amplitudes for the first and second pulse given by the SRP model. We wish to find optimal parameters for the efficacy kernel, 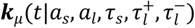, such that we arrive at a local minimum in a cost function comparing the experimental and theoretical charges for various delays *τ*_*i*_ of the paired-pulse stimulation.

We start by deriving an analytical expression for (5) in terms of the underlying efficacy kernel.

The first spike’s efficacy is taken by evaluating the efficacy kernel at time 0:

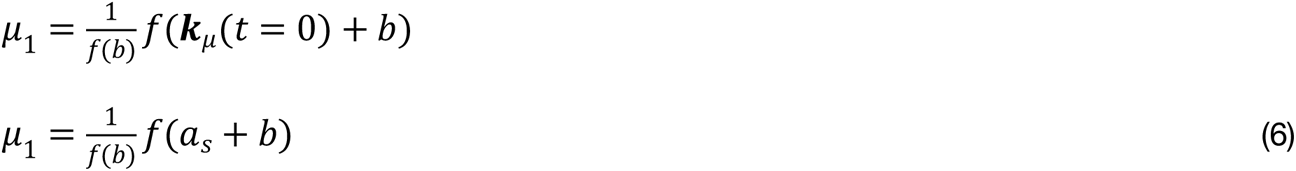

The second spike’s efficacy is taken by evaluating the sum of an efficacy kernel at time *t*_*i*_ (efficacy from first spike at the time of spike two) and an efficacy kernel at time 0 (second spike’s efficacy).

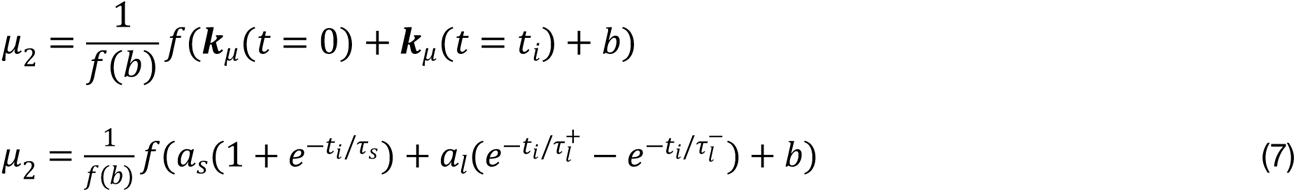

Substituting (6) and (7) into (5):

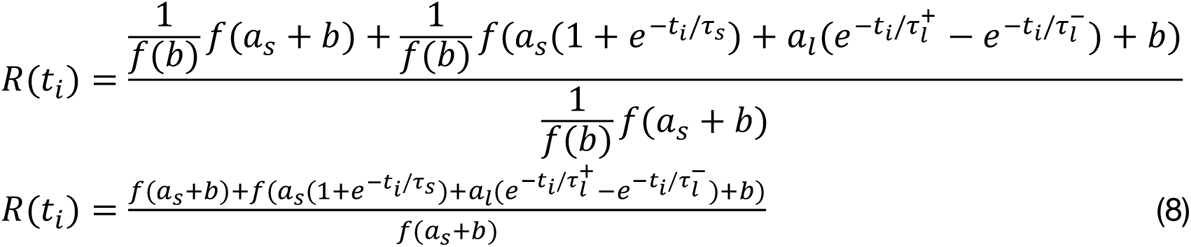

Equation (8) and the corresponding experimental results (5) are used to perform least-squares optimization on the parameters 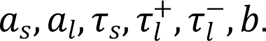 This yields the following parameter estimates:

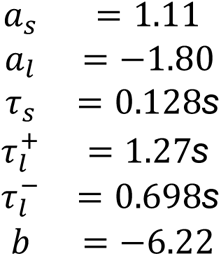

#### A general expression for spike efficacy given past spiking history

One can also write a solution to the generalized case of calculating the efficacy, *μ*_*n*_, of a certain presynaptic spike *n* occurring at time *t*_*n*_, given a past history of spikes with times *t*_1_, *t*_2_, *t*_3_, … *t*_*n*−1_:

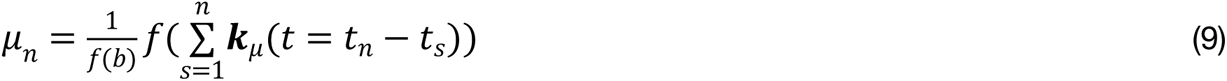

This equation is used to calculate the model’s predictions for the matched pulse-chase experimental data.

#### Network model of recurrently connected 5-HT neurons receiving LHb input

To construct a simplified model of the DRN network, we took a modular approach where each individual component was fitted to experimental data where possible. Our model consisted of 5000 conductance-based leaky-integrate-and-fire 5-HT neurons, building on recent simplified single-cell modeling of 5-HT neurons (Harkin et al., 2023). 25% of 5-HT neurons received excitatory input from LHb (∑7.5*nS*), while all 5-HT neurons received background Poisson excitation as well as 5-HT1AR-mediated inhibition from other 5-HT neurons (∑2.5*nS*). The 5-HT synapse of recurrent inhibitory network were imparted with both the experimentally-fitted event kinetics and the short-term dynamics described above (i.e., from the SRP model).

Individual 5-HT neurons were fit with a simplified leaky-integrate-and-fire model. The fitting was performed experimentally according to previously described methods (Pozzorini et al., 2015). Briefly, identified serotonin neurons from SERT-CRE-tdTomato mice were recorded in whole-cell current-clamp configuration, and a noisy input stimulus was presented through multiple test and training repetitions. These repetitions were used to train a generalized integrate-and-fire model to each individual serotonin neuron by maximizing the likelihood of the model (for full details, see Harkin et al., 2023).

In this work we focused on basic extracted parameters from this model:*g*_*leak*_, *E*_*leak*_, *C*_*m*_, *t*_*refractory*_, *V*_*thresh*_. While Harkin et al., (2023) utilized a current-based generalized-integrate-and-fire model to represent serotonin neuron firing, here we consider a simplified, conductance-based, leaky-integrate-and-fire model of individual serotonin neurons:

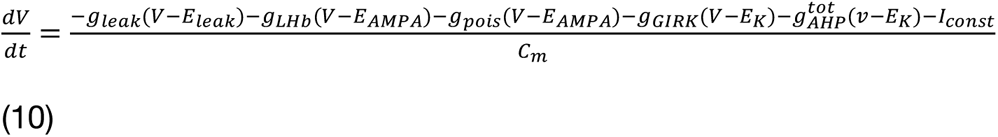

With the fitted parameters:

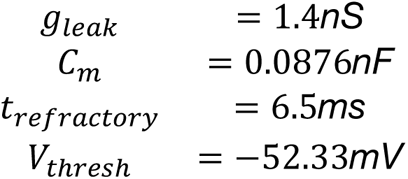

And the chosen ionic reversal potentials, which, for the purposes of this LIF model, are considered as fixed parameters:

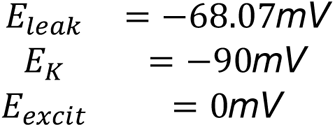

Conductances for *g*_*LHb*_, *g*_*pois*_, *g*_*GIRK*_, *g*_*AHP*_ were considered with the following dynamics, where *s* is a synapse state variable which instantaneously increments by 1 whenever a presynaptic neuron fires, and exponentially decays to 0 with a characteristic time-constant:

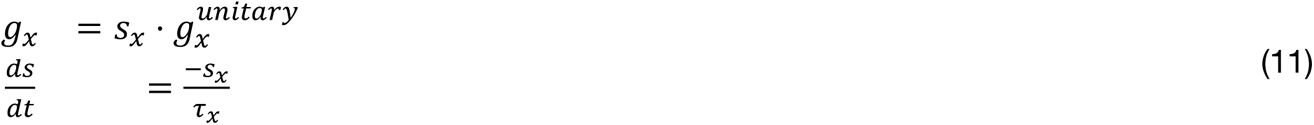

where *x* represents conductances of either *LHb*, *pois*, *GIRK* or *AHP*.

The AHP conductance is expressed as a sum of three unitary conductances with distinct timescales.

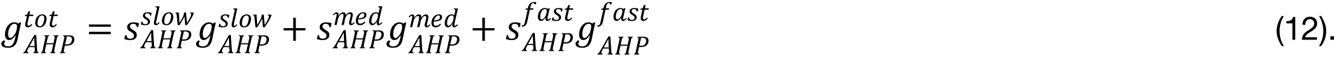

The following values were chosen for individual parameters of the model. Note that excitatory decays were chosen to reflect a mixture of both AMPA and NMDA currents. Note that these reflect unitary conductance values, and neurons receive multiple LHb (*g*_*LHb*_) and recurrent GIRK (*g*_*GIRK*_) inputs, as well as a constant background rate of Poisson inputs (*g*_*pois*_).

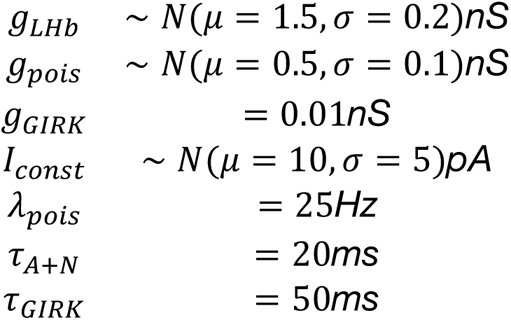

For the AHP dynamics, the following values were extracted from fits on identified 5-HT neurons using a previously described protocol (see Pozzorini et al., 2015; Harkin et al. 2023):

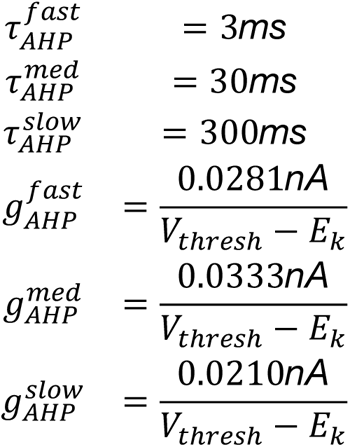

Note that since the generalized integrate-and-fire model in Pozzorini et al. (2015) extracts a spike-triggered current for the AHP, we convert this to a conductance by dividing by the driving force at spike threshold.

The DRN network model consisted of 5000 5-HT neurons with the LIF dynamics described above. 5-HT neurons were recurrently connected in an unstructured manner with 5% random connection probability and 0.01 nS unitary GIRK synaptic conductance (∑2.5*nS* net GIRK inhibition received by each neuron before facilitation/depression dynamics are considered). Short-term plasticity between 5-HT neuron connections was modelled using the linear-nonlinear description fit to data, described in the preceding section. 25% of DRN 5-HT neurons receive long-range input from a cluster of 500 LHb neurons emitting Poisson spike trains at rates specified in the text, where each LHb neuron and 5-HT neuron is randomly connected with 1% probability (*i.e.,* each 5-HT neuron receives an average of 5 LHb synapses) with synapse strength specified above. This yields a mean LHb conductance of ∑7.5*nS* for each 5-HT neuron.

#### Network model of recurrently connected 5-HT neurons receiving LHb and A input (Winner-take-all computation)

In the simulations outlined in Fig. 7, we considered a DRN network where 25% of neurons received LHb input, and 25% of neurons received input from a generic area termed Area A. In these simulations, the basic neural architecture and dynamics were identical to that described above, except that 25% of 5-HT neurons now received input from Area A (non-overlapping with the 25% of neurons already receiving LHb input). Area A consisted of 500 neurons emitting Poisson spikes at rates specified in the text, where each A neuron and 5-HT neuron is randomly connected with 1% probability (*ie* each 5-HT neuron receives an average of 5 A synapses). This yields a mean A conductance of ∑7.5*nS* for each 5-HT neuron. Synapse strength is identical to that of LHb-5-HT synapses, as are decay kinetics:

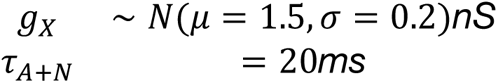

#### Exploration of changes in network connectivity parameters

In some simulations, we varied the fraction of 5-HT neurons innervated by long-range LHb input, from 10-50% (Fig. S14). We sought to compute several network-wide metrics corresponding to recurrent drive, in order to assess whether these metrics were tracking input frequency, or alternately input fraction. The intuition behind these analyses is that the magnitude of the inhibitory synaptic conductance *g*_*GIRK*_, associated with recurrent connections, provides a population-level metric of the strength of recurrence. First, for each simulated 5-HT neuron, we computed the maximum post-stimulus *g*_*GIRK*_ conductance, and we then constructed a cumulative distribution of this *g*_*GIRK*_ variable across the population (Fig. S14D, S14I). Second, we computed the population mean of *g*_*GIRK*_ for each combination of input fraction and input rate (Fig. S14E, S14J).

In other simulations, we varied the spatial structure of recurrent connectivity (local vs global recurrence) to assess its impact on winner-take-all computations (Fig. S16). The probability of recurrent connections was considered as a Gaussian distribution with a peak amplitude of the maximum connectivity (p=0.05), with standard deviations that varied according to the spatial structure of recurrent connectivity. We computed a simple metric relating the standard deviation of the connectivity (in units of neurons) to the total number of neurons of the network, *c* = *σ*/*n*_5–*HT*_, which we report in the text. In these network simulations and simulations in Fig. S17, we sought to additionally compute a metric of the LHb-driven change in the response patterns of A-recipient 5-HT neurons. We compared the rates of these A-recipient neurons when receiving LHb input at a given rate, versus receiving no LHb input. This was computed as 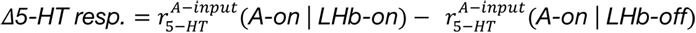. We interpret this metric as quantifying the strength of the winner-take-all interaction and utilize it to explore the parameter space of combinations of rates of A and LHb.

We additionally varied the spatial overlap of inputs from LHb and A (Fig. S17). In these simulations, each input innervated a total of 25% of 5-HT neurons, but the fraction of neurons receiving overlapping input from both LHb and A, *o*, is varied as a connectivity parameter across simulations. All other connectivity parameters are identical to in the main simulations. We computed the metric quantifying the strength of winner-take-all interactions as described above.

Finally, in many simulations we removed the short-term plasticity rules at recurrent connections. This was implemented by altering the plasticity kernel to be a flat line over time, such that any presynaptic input would produce a consistent postsynaptic response that was history-independent. All other parameters, including recurrent connectivity, were not altered.

### Auditory classical conditioning task

#### Surgery

All experiments and procedures were performed in accordance with approved procedures and guidelines set forth by the University of Ottawa Animal Care and Veterinary Services. For stereotaxic injections, mice were injected with 0.05 mg/kg buprenorphine 1 hour prior to surgery and anesthetized by inhalation of isoflurane (4% for induction and 1.5-2% for maintenance). The head was shaved, ointment was applied to each eye, and an incision was made above the skull. A dental drill was used to perform skull craniotomies at locations above the coordinates for LHb and DRN described below.

For stereotaxic injections in LHb, we utilized the virus AAV9.CAG.hChR2(H134R)- mCherry.WPRE.SV40 at a concentration of ∼10^-12^ GC/mL. Injections were performed using a 10 μl Hamilton Neuros syringe with point style 4 (12° beveled needle point). 0.4 - 0.5 μl was injected per coordinate at a rate of 0.1 – 0.2 μl per minute. Stereotaxic coordinates for injection are as follows for LHb, from Bregma skull landmark, in mm: AP +1.7, ML +/- 0.4, DV −3.1 to - 3.3. Following stereotaxic injections, we affixed a custom stainless steel or 3D-printed Acrylonitrile butadiene styrene (ABS) headbar to the anterior surface of the skull with C&B Metabond Quick Adhesive Cement (Parkell).

For optic fiber implantation in DRN, each animal received a fiber (Doric Mono Fibre-Optic Cannula, 4mm length, part number: MFC_100/125-0.37_4mm_ZF1.25_C45) which was implanted at the following coordinates, from Bregma skull landmark in mm: AP-4.6, ML 0, DV - 2.9. In some animals, we used an angled injection to avoid the superior saggital sinus (20° angle, from Bregma skull landmark in mm: AP-4.6, ML +1, DV −3.25). Optic fibers were slowly lowered to the desired dorsoventral location, and affixed with C&B Metabond Quick Adhesive Cement (Parkell) followed by Jet Denture repair cement (Lang) to provide structural stability to the implant.

After surgery, animals received topical bupivicaine (2%) and intraperitoneal (IP) carprofen (20 mg/kg). Post-operative care consisted of a further bupivicaine treatment at 4-6h post-surgery, and IP carprofen injection 4-6h post-surgery and for the next two days. Animals were allowed to recover from surgery for one week before water restriction was started. All animals were housed separately.

#### Behavior, water restriction and optogenetic stimulation

Animals were placed on water restriction for 7 days prior to the start of the task. During this time, they were habituated to the experimenters and the apparatus through handling and brief periods of head fixation while daily water (0.8 - 1 mL) was provided. After habituation, for 2-3 days each mouse was head-fixed for short bouts and 5-10 uL apple juice was delivered to a lickport every 1 to 8 s without preceding auditory tones. Following this, mice underwent 1-2 weeks of training on the primary auditory classical conditioning task, described below.

The auditory classical conditioning task consisted of 120 randomly interleaved trials presented to each mouse per day, giving a total reward value of ∼0.6 - 0.8 mL per mouse per day. Inter-trial intervals were drawn from an exponential distribution with *λ* = 5*s*, where 2*s* < *ITI* < 14*s* to prevent excessively long or short ITIs. Exponentially distributed ITIs give the convenient property that trials follow a Poisson distribution in time, and therefore are memoryless (the mice cannot predict trial onset times). To prevent the mice from licking continuously during the baseline periods, new trial starts were contingent on baseline mouse licking of <10% (fractional time). Three trial types were randomly interleaved: a *large-reward trial*, consisting of a 3 kHz tone and a 10uL apple juice reward, a *small-reward trial*, consisting of a 6 kHz tone and 5 uL apple juice reward, and a *no-reward trial*, consisting of a 12 kHz tone and 0 uL apple juice reward. For each trial, the tone lasted 1 s, the trace period between tone and reward delivery was drawn from a Gaussian distribution with *μ* = 2*s*, *σ* = 4*s* (values of *t*_*trace*_ < 0.5*s* were prohibited), and the apple juice reward was concomitantly delivered with a 500 ms white noise tone to aid learning. Following reward delivery, licking was monitored for 4 s before the trial was terminated. This auditory classical conditioning task was presented daily to mice for 1-2 weeks until criterion performance had been achieved for >2 consecutive days. Criterion performance was defined as a mean lickrate of >0.5 Hz during the trace period of large-reward trials. While many mice displayed mean lickrates which were significantly higher than this (Fig. 8C), this was a convenient lower threshold for criterion performance.

After criterion performance was reached, mice experienced the auditory classical conditioning paradigm while we optogenetically activated the habenulo-raphe pathway in some trials. The trial structure consisted of 120 trials of interleaved large-reward, small-reward and no-reward trials as before. Each trial was composed of the same tone and reward associations and durations as described above, except the period between tone and reward was fixed at 2 s.

For the first 40 trials, no optogenetic stimulation was delivered (*baseline block*). For the final 80 trials, optogenetic stimulation was delivered on 40 randomly chosen trials (*optogenetic stimulation block*). Optogenetic stimulation was composed of pulses of 10 ms width, at frequencies noted in the text (5Hz or 20Hz), delivered during the period between tone and reward. Optogenetic stimulation was delivered intracranially via an optic fibre implanted in DRN, to which a patch cable was connected before the session.

The behavioral apparatus consisted of a sound-insulated box containing a Raspberry Pi 3B, custom steel head fixation apparatus and equipment for delivering stimuli and acquiring signals. Lick signals were acquired via a custom 3D-printed combined infrared lickometer and stainless steel lickport. Lickport placement was adjusted daily for each mouse to ensure optimal performance and signal acquisition. A solenoid valve (NResearch 2-way NC isolation valve) gated apple juice delivery, and flow rate was calibrated weekly. Speakers connected to the Raspberry Pi (Logitech Z150) delivered auditory stimuli. We visualized animal behavior with an infrared video camera connected to the Raspberry Pi (Raspberry Pi Camera Module 2 NoIR). In vivo optogenetic stimulation was delivered using a PlexBright Compact LED module and controller (either blue light, 465; or yellow light, 590 nm; as noted in the manuscript; Plexon, Texas) and a PlexBright optical patch cable. A custom script (Python 3.8) on the Raspberry Pi registered lickometer data at 200 Hz while controlling trial structure, auditory stimulus delivery, reward delivery and optogenetic stimulation. Data was stored in hdf5 format.

After behavioral testing was completed for individual mice, they were perfused transcardially with 4% paraformaldehyde, and brains were cut in 50-100 um sections. Immunohistochemistry was performed as described above, and the injection sites and optic fiber tracts were verified.

#### Data analysis

Data was analyzed with custom Python 3.8 scripts, employing the numpy (van der Walt et al., 2011), scipy (Jones et al., 2001), h5py and matplotlib technical computing libraries. Briefly, raw licking data was binarized and converted into lick onset times. For visualizations (Fig. 8E-F), licking signals were calculated by convolving lick onset times with a gaussian kernel (*σ* = 50*ms*) and averaging across all trials of that type. Optogenetic analyses are described below.

Optogenetically evoked changes in anticipatory (trace) licking were calculated as follows: we considered trials of a given type during the optogenetic stimulation block (last 80 trials), and calculated the mean licking frequency in the period between the tone and reward (anticipatory licking) for each trial. To calculate an optogenetically evoked change in anticipatory licking, the mean anticipatory licking frequency for optogenetically stimulated trials was subtracted from the mean anticipatory licking frequency for non-stimulated interleaved trials during the optogenetic stimulation block. Optogenetically evoked changes in reward licking were calculated in a similar way, but the period considered was the post-reward epoch.

We additionally considered whether optogenetic stimulation modulated behavior beyond the stimulated period (Fig. 8I). For these analyses, we compared nonstimulated trials during the baseline block with nonstimulated interleaved trials during the optogenetic stimulation block. For each trial of a given type, we first computed mean anticipatory licking frequency as described above. Next, to quantify long-term optogenetically evoked changes in anticipatory licking, we subtracted mean anticipatory licking frequency for nonstimulated trials during the optogenetic stimulation block from mean anticipatory licking frequency for trials during the baseline block.

---

**Figure S1:**
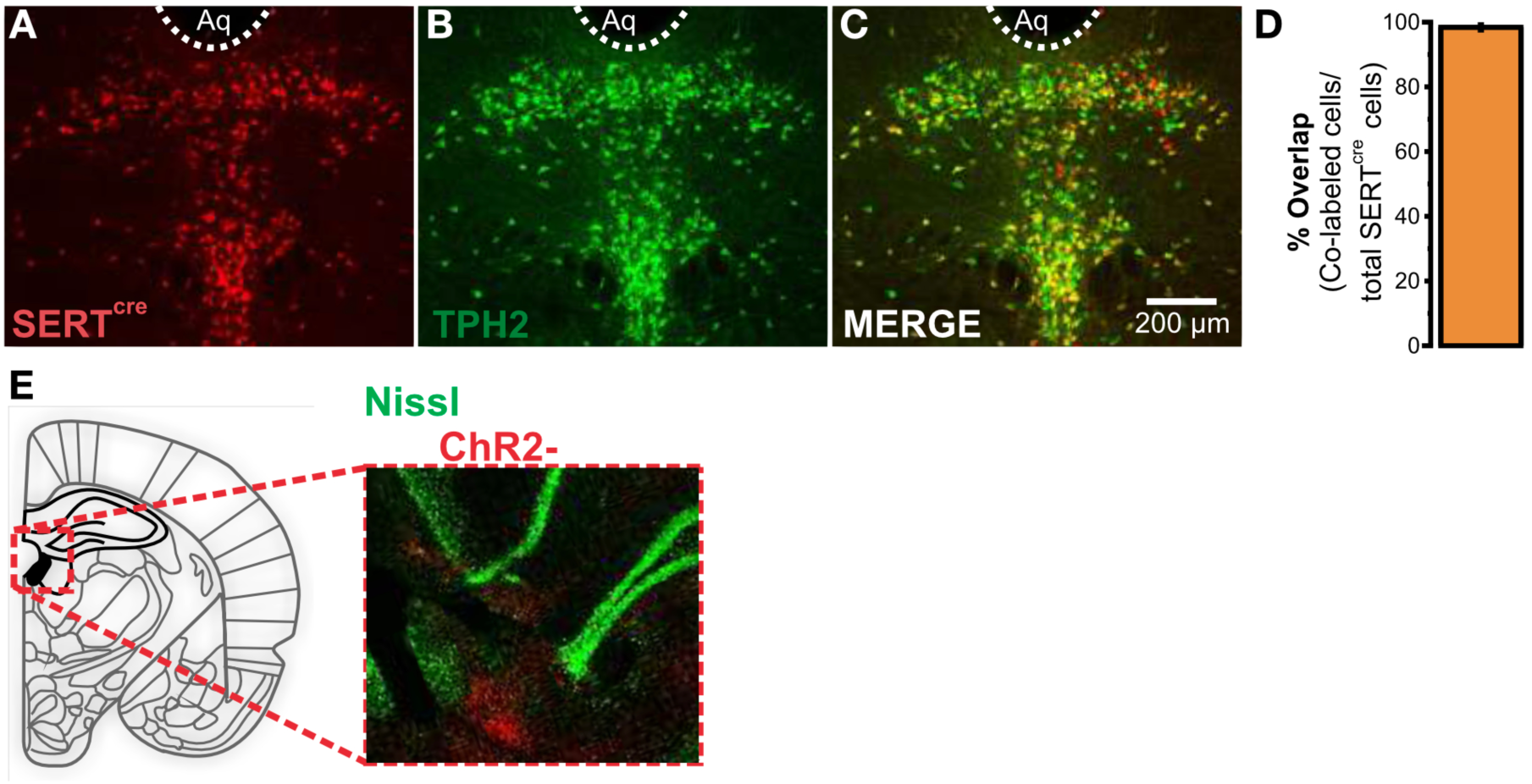
Validation of SERT-CRE-tdTomato mouse line and LHb injection site. Confocal image of SERTcre **(A)** and TPH2-expressing **(B)** 5-HT neurons in the DRN. **(C)** Merge confocal image of SERTcre and TPH2 channels. **(D)** Bar plot of overlap between SERT-cre and TPH2-expressing 5-HT neurons in the DRN (98.38 +/- 0.8; n = 3, mean +- SE). **(E)** Representative image of LHb injection site (AAV-ChR2-mCherry).

**Figure S2:**
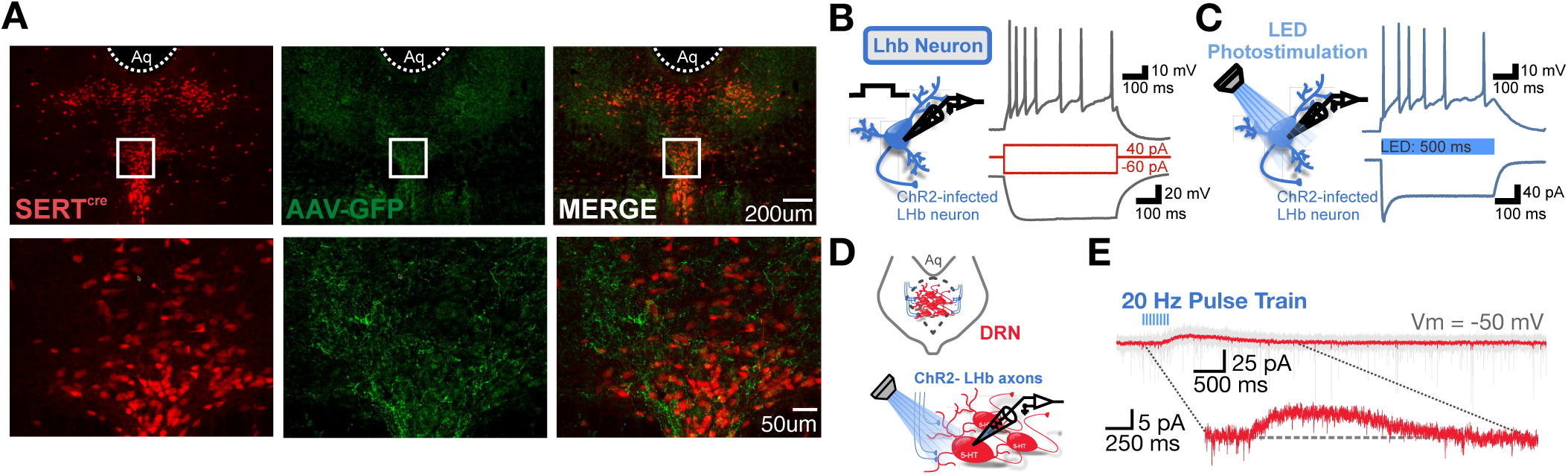
LHb sends axonal projections to the DRN. **(A)** Confocal image of GFP-expressing LHb axons in the DRN with 5-HT neurons identified with SERT-TdT. **(B)** ChR2-infected LHb neuron can be induced to spike with current injection. **(C)** LED photostimulation produces qualitatively similar spiking patterns to those induced by current injection. **(D)** Experimental schematic for long-range optogenetic interrogation. **(E)** Example serotonin neuron which does not receive direct LHb input, but displays a delayed heterosynaptic inhibitory current.

**Figure S3:**
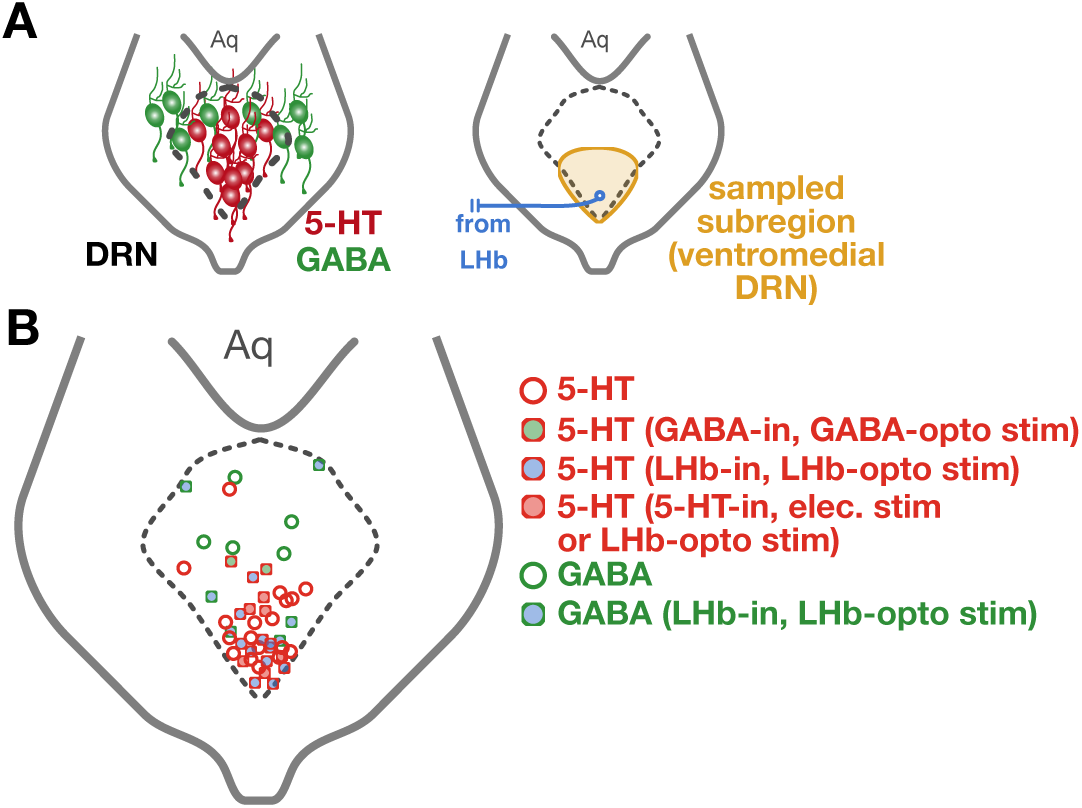
Anatomical location of 5-HT and GABA neurons recorded in the whole-cell configuration from the dorsal raphe nucleus. **(A)** Left, schematic of DRN and the location of cell types within DRN. Right, schematic of the targeted subregion of DRN for electrophysiological experiments. **(B)** Representative subsample of the anatomical locations of 5-HT and GABAergic neurons, projected onto the dorsoventral and mediolateral axes (n=48 neurons). Images were taken using a 5x objective after electrophysiological recording and were registered manually. Circles with shading denote a response as described in the legend, related to electrophysiology experiments described in Figures 1, 2 and 3, as well as Figure S4 and S10. GABA-in, LHb-in and 5-HT-in refer to the observed presence of GABAergic conductances, direct excitatory conductances from long-range LHb activation, and indirect serotonergic inhibitory conductances, respectively. LHb-opto stim and GABA-opto stim refer to optogenetic stimulation experiments described in Fig S10H-K, Fig. 1 and Fig. S4, while elec.-stim refers to electrical stimulation experiments described in Fig. 3 and Fig. S10A-G.

**Figure S4:**
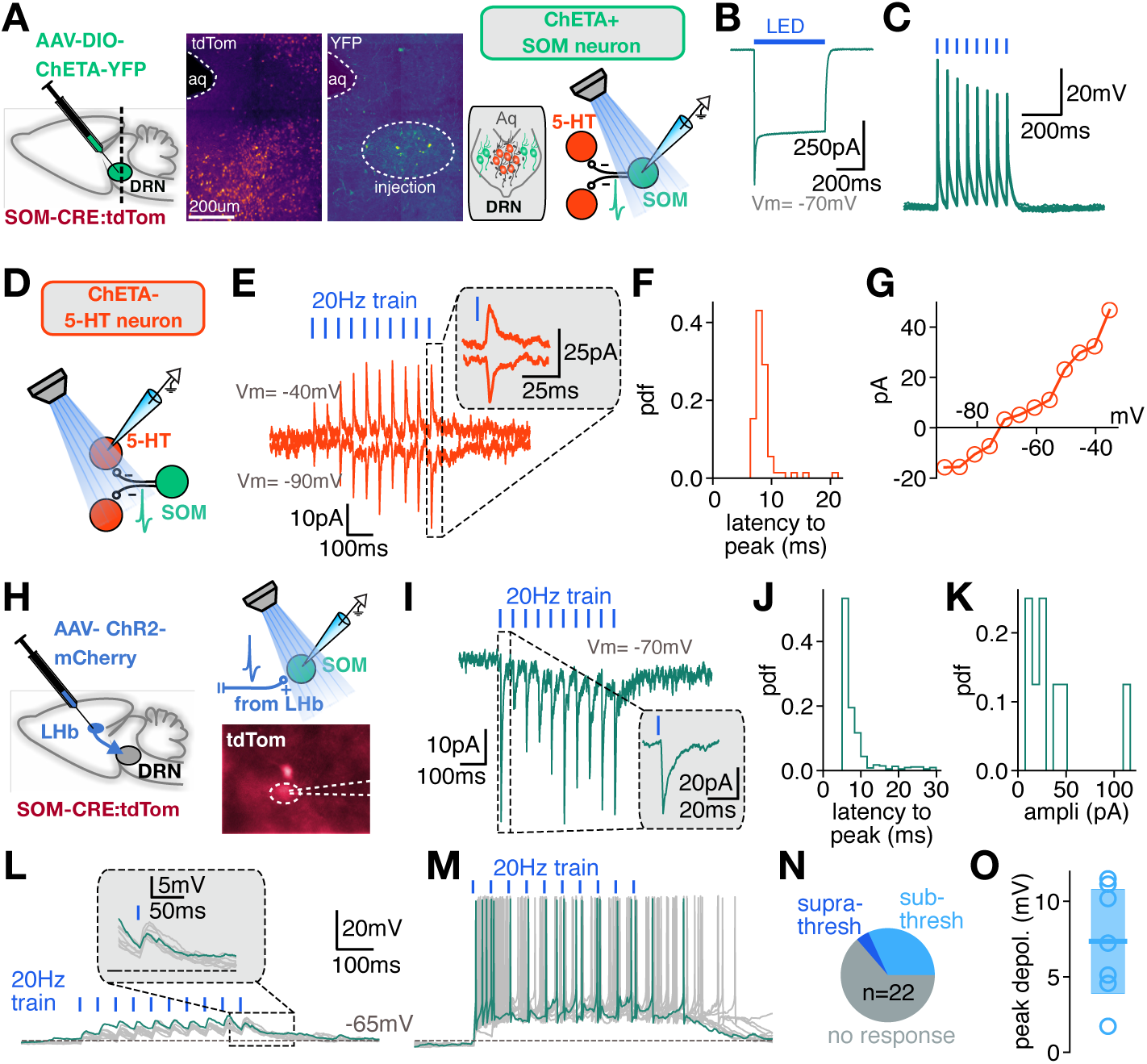
Lateral habenula input fails to recruit strong disynaptic inhibition onto 5-HT neurons located in the ventromedial portion of DRN. **(A)** Experimental setup for panels B-G. Schematic showing viral expression strategy (left), confocal images showing CHETA-YFP and tdTomato expression in DRN (middle), and schematic of whole-cell patch clamp recording setup from SOM neurons, related to panels B and C (right). **(B)** Example trace from CHETA-YFP-expressing 5-HT neuron recorded in voltage clamp, demonstrating photostimulation-evoked inward current. **(C)** Example trace from CHETA-YFP-expressing 5-HT neuron recorded in current clamp, demonstrating photostimulation-evoked spiketrains at 20Hz. **(D)** Schematic of whole-cell patch-clamp recording setup from 5-HT neurons, related to panels E-G. **(E)** Trial-averaged traces from an example 5-HT neuron, recorded in voltage clamp, illustrating photostimulation-driven currents at two holding potentials. Inset depicts zoomed region. **(F)** Latency to peak outward current response at Vm=-40mV across all photostimulation events (n=120 events) for one example neuron. **(G)** I-V curve for photostimulation-driven currents from one example neuron. **(H)** Experimental setup for panels I-O. **(I)** Trial-averaged trace from an example SOM neuron, recorded in voltage clamp, illustrating photostimulation-driven inward currents. Inset depicts zoomed region. **(J)** Latency to peak inward current response across all photostimulation events (n=615 events, 8 neurons). **(K)** Amplitude of peak current response across all photostimulation events (n=615 events, 8 neurons). **(L-M)** Current-clamp traces from two example SOM neurons showing subthreshold responses to photostimulation **(L)** and suprathreshold responses to photostimulation **(M)**. **(N)** Classification of SOM responses to photostimulation across the population of recorded neurons. **(O)** Peak depolarization from baseline for SOM neurons exhibiting subthreshold responses (n=7). Line and shaded area express mean +-standard deviation.

**Figure S5:**
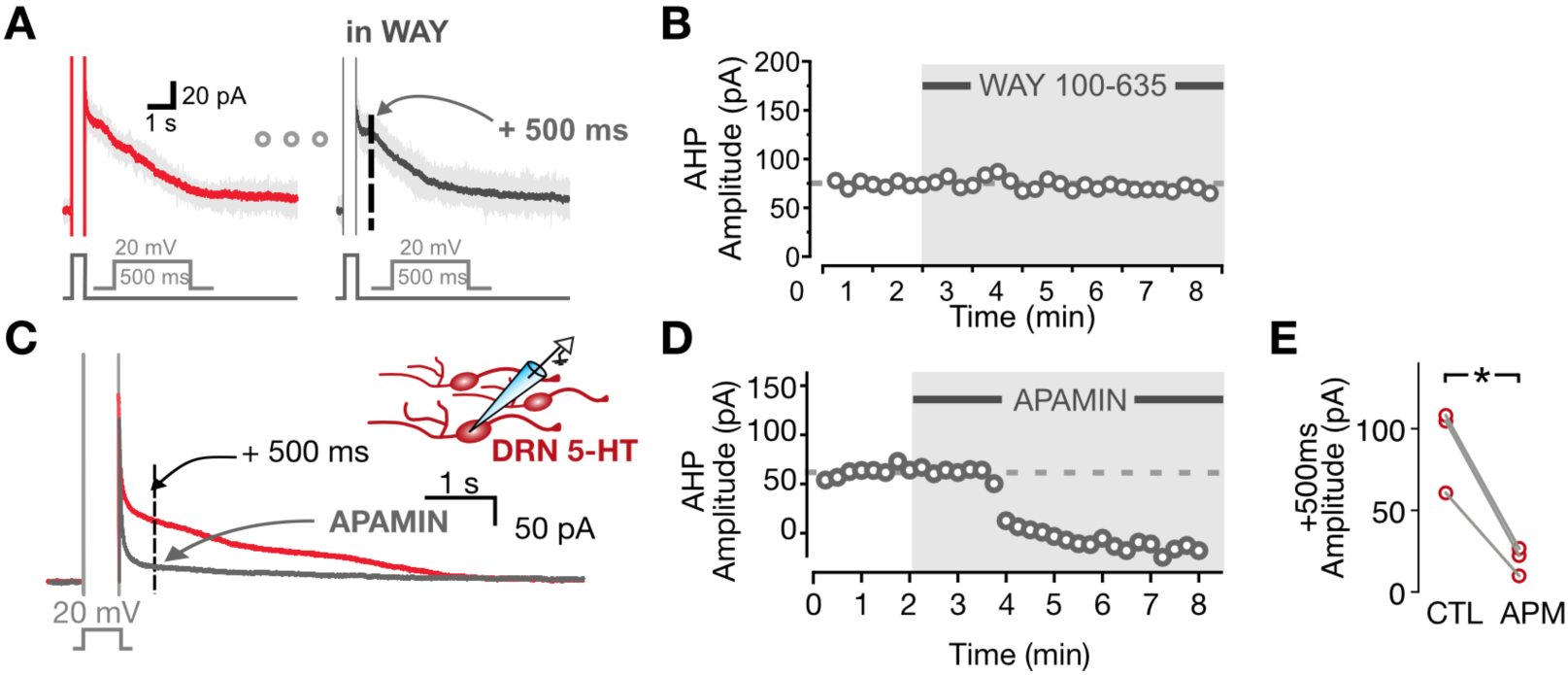
Voltage-clamp experiments provide evidence that the afterhyperpolarization of serotonin neurons is mediated by SK channels, not 5-HT1ARs. **(A)** Representative traces of AHP current (IAHP) driven by a 500 ms depolarization step to 20 mV in voltage clamp. **(B)** Representative scatter plot showing no differences in IAHP amplitude following WAY 100-635 application. **(C)** Representative IAHP traces before and after APAMIN application (300 nM). **(D)** Representative scatter plot showing the IAHP +500 ms amplitudes in response to APAMIN. **(E)** +500 ms IAHP amplitudes before and after APAMIN application (CTL, 91.1 +/- 15.3; APM, 19.6 +/- 5.0; n = 3; p = 0.02, paired two-sided t-test).

**Figure S6:**
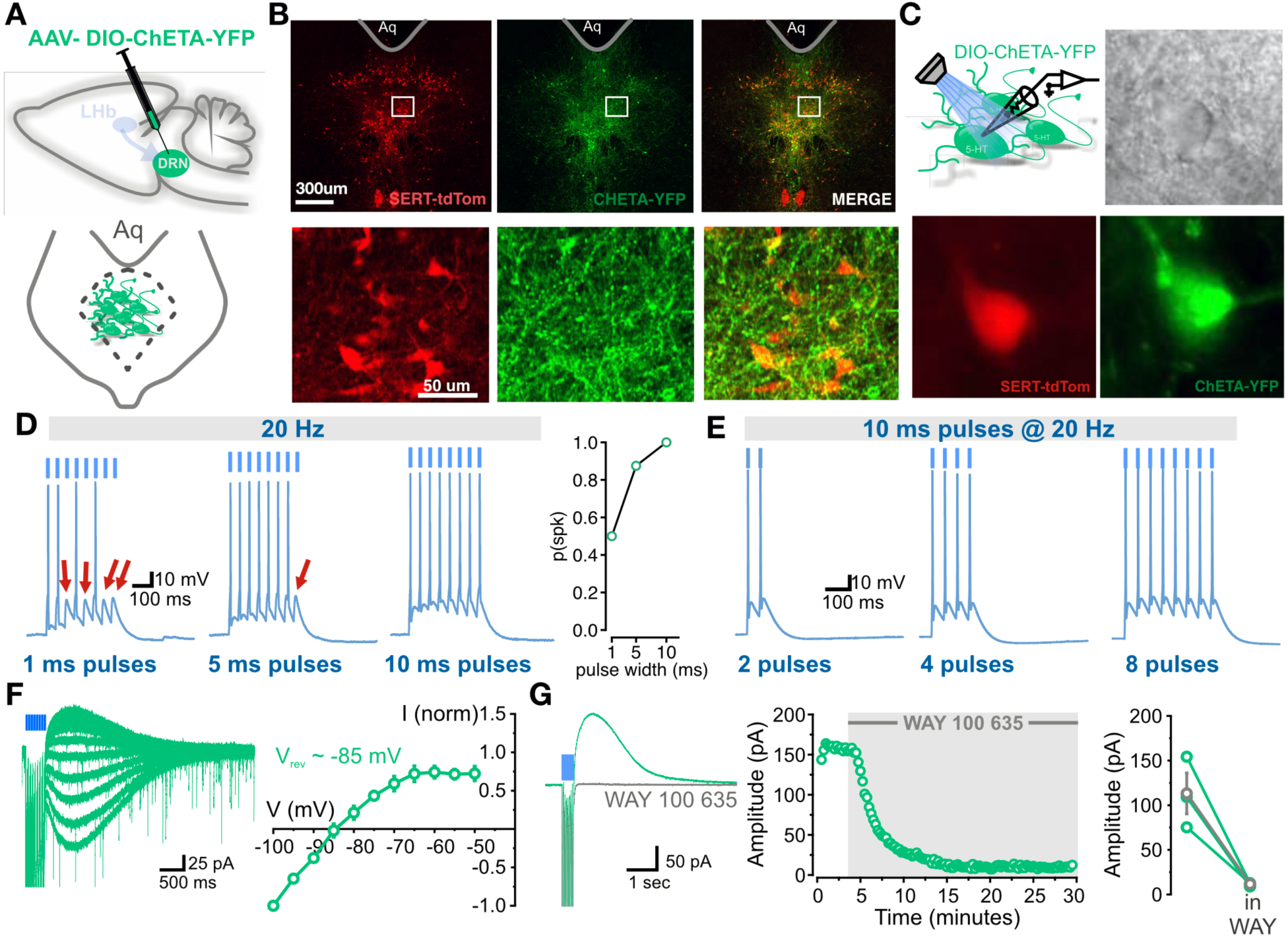
Validation of local CHETA optogenetic strategy in DRN. **(A)** Schematic depicting AAV-DIO-ChETA-YFP injection into the DRN of SERTcre-TdT mice. **(B)** Confocal image of ChETA-YFP expression in tdTomato-positive neurons in the DRN. Above, low magnification (scale, 300 μm) ; below, high magnification (scale, 50 μm). **(C)** Schematic and EPI-flourescent image of whole-cell recordings from ChETA-YFP-expressing SERT-TdT neurons. **(D)** Left, LED stimulation of 5-HT neuron spiking with different pulse durations at frequency of 20 Hz. Right, quantification of the probability of photostimulation eliciting a spike as a function of pulse width (n=8 pulses). **(E)** LED stimulation of 5-HT neuron spiking in response to varying number of 10 ms pulses at 20 Hz. **(F)** Representative IV relationship traces and IV relationship (n = 5, mean +- SE). **(G)** Representative traces and scatter plot showing complete blockade of outward current by WAY 100-635. Right, population data is displayed (Pre-WAY, 112.9 +/- 22.9; Post-WAY, 11.5 +/- 1.1; n = 3; p = 0.02, paired two-sided t-test).

**Figure S7:**
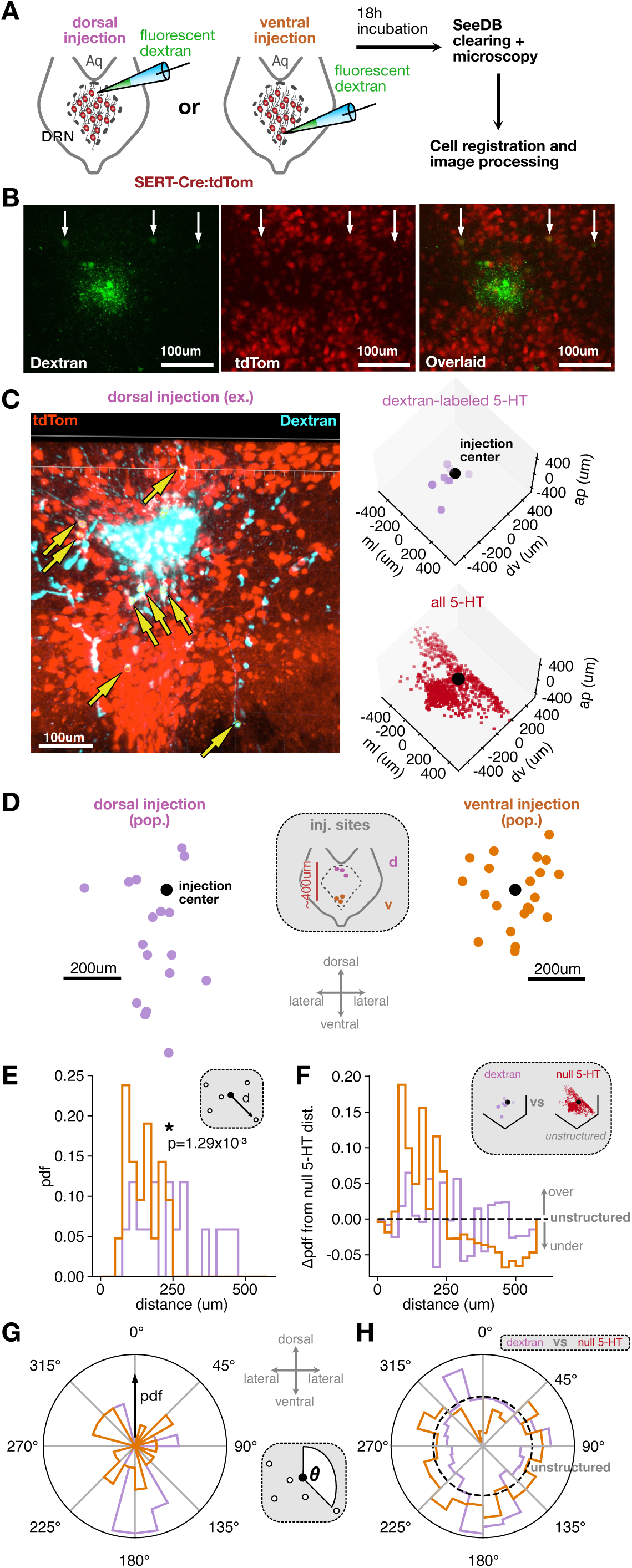
Spatial scale and anatomical features of recurrent connectivity. **(A)** Experimental schematic demonstrating fluorescent dextran labeling approach. **(B)** Representative confocal images of DRN showing dextran injection site and labeled neurons (left), tdTomato neurons (middle) and overlaid image showing colocalization (right, arrows). **(C)** Representative confocal image (left) depicting dextran injection in dorsal DRN (cyan), tdTomato-positive 5-HT neurons (red), and neurons exhibiting colocalization (arrows). Extracted coordinates are shown for colabeled 5-HT neurons (middle) and all 5-HT neurons (right). **(D)** Locations of all colabeled 5-HT neurons from injection center for dorsal injection (left; n=17 neurons, 3 injections) and ventral injection (right; n=21 neurons, 3 injections). Inset depicts anatomical locations of each injection site within DRN. **(E)** Probability density of the distance from colabeled 5-HT neurons to injection center, for dorsal (purple) and ventral (orange) injections (K-S test for statistical comparison). **(F)** Probability density of colabeled neuron distance, expressed as a difference in probability from the distribution of all 5-HT neurons (unstructured, dotted line). **(G)** Polar plot depicting anatomical location of colabeled 5-HT neurons from injection site, for dorsal (purple) and ventral (orange) injections. **(H)** Polar plot of colabeled neuron location as in G, but expressed as a difference in probability from the distribution of all 5-HT neurons (unstructured, dotted line).

**Figure S8:**
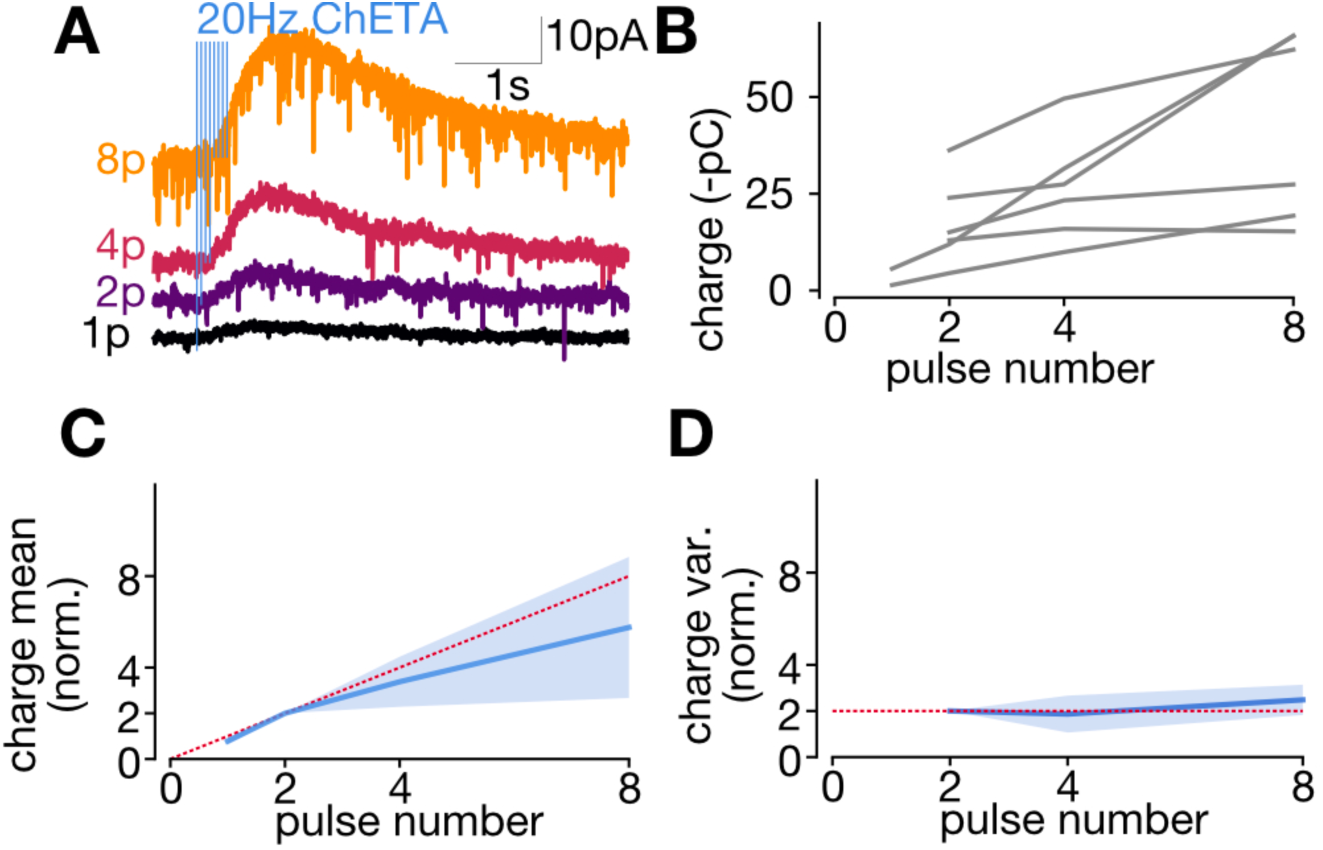
Postsynaptic dynamics of recurrent 5-HT1A connections indicate non-desensitizing receptors. **(A)** Representative averaged traces from a single non-CHETA-expressing 5-HT neuron depicting responses to 20Hz optogenetic stimulation of DRN for various pulse numbers. **(B)** Event charge for each neuron as a function of pulse number at 20Hz (n=6). **(C)** Charge mean as a function of pulse number, normalized to the mean at two pulses (n=6, mean +-SD). Dotted red line represents a linear relationship between pulse number and evoked charge. **(D)** Charge variance as a function of pulse number, normalized to the variance at two pulses (n=3, mean +-SD). Dotted red line represents predicted variance for release probability p=1.

**Figure S9.**
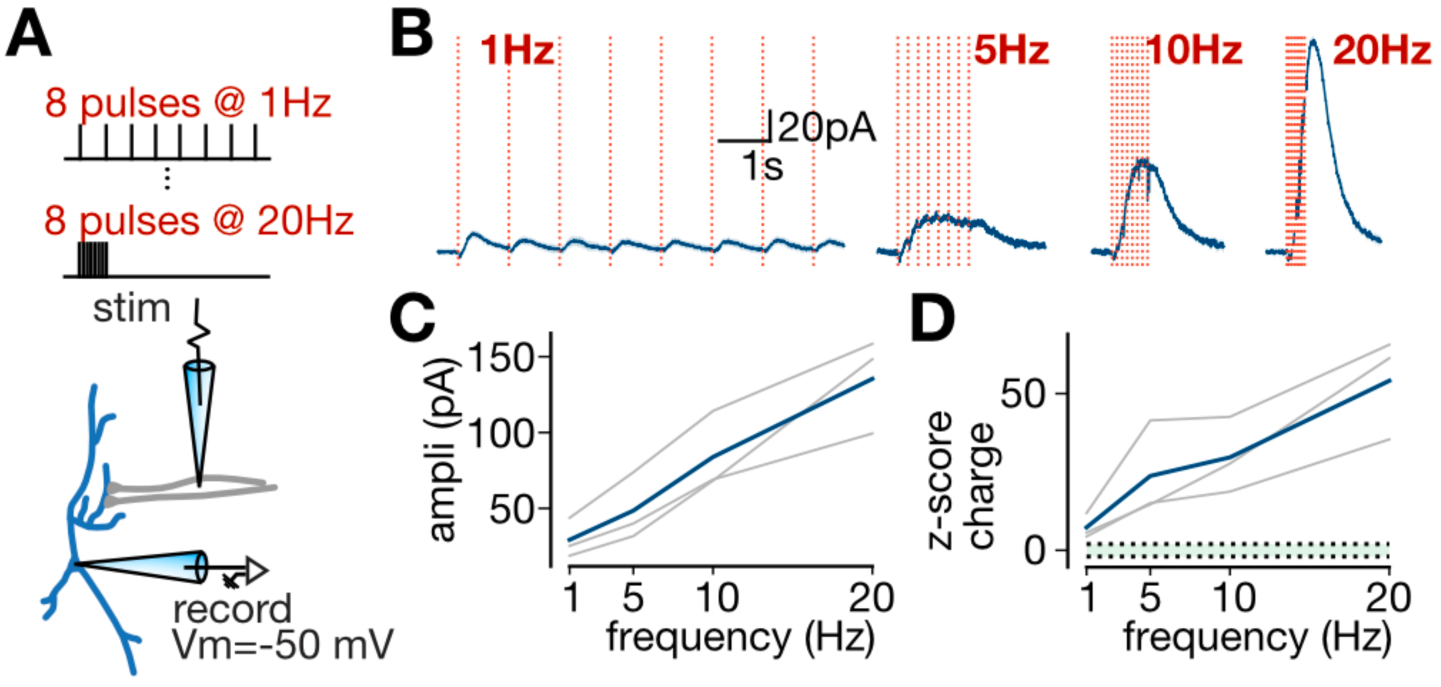
Frequency dependence of 5-HT1A connections at suprathreshold stimulation frequencies. **(A)** Experimental setup depicting electrical stimulation of 5-HT1A synapses while recording from an identified 5-HT neuron in the whole-cell configuration. **(B)** Representative averaged traces from a single neuron depicting responses to electrical stimulation at various frequencies, with pulse number held constant. **(C)** IPSC amplitude as a function of frequency, for individual neurons (grey) and the mean of all neurons (blue, n=3). **(D)** Event charge, expressed as a z-score relative to a noise distribution, as a function of frequency. Depicted are individual neurons (grey) and the mean of all neurons (blue, n=3). Green shaded area represents 95% confidence interval for noise distribution.

**Figure S10.**
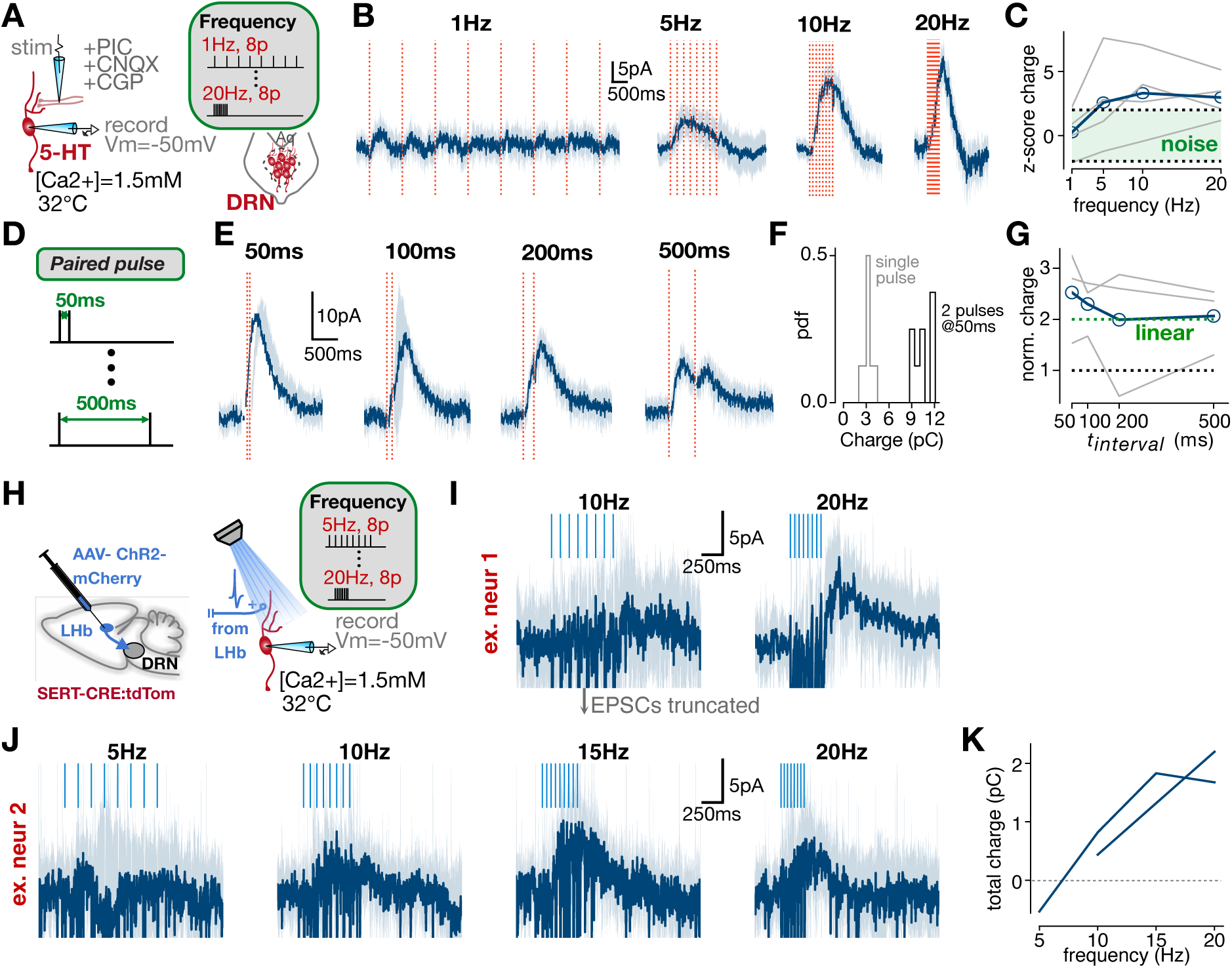
Frequency-dependent facilitation of serotonin release under low extracellular calcium conditions. Related to Figure 3. **(A)**: Experimental setup for panels B-C depicting whole-cell patch clamp recording from genetically identified 5-HT neuron with electrically evoked 5-HT release, under physiological recording conditions (extracellular calcium and temperature). Inset depicts electrical stimulation parameters for frequency-varying experiments. **(B)** Representative averaged traces from a single neuron depicting responses to electrical stimulation at various frequencies. Red dotted lines denote electrical stimulation, and the shaded area denotes standard deviation across trials. **(C)** Charge z-score as a function of electrical stimulation frequency, with 95% noise C.I shown in green (n=4 neurons). **(D)** Schematic showing electrical stimulation parameters for paired-pulse experiments (E-G). **(E)** Representative averaged traces from a single neuron depicting responses to the paired-pulse electrical stimulation protocol. **(F)** Charge histograms for single-pulse responses and paired-pulse responses at 50ms for one example neuron. **(G)** Post-stimulus charge as a function of paired-pulse time interval (n=4 neurons). Green line depicts expected charge from linear summation. Blue line depicts population average and grey lines are responses of individual neurons. **(H)** Experimental setup for panels I-K depicting channelrhodopsin injection in LHb (left), and whole-cell patch clamp recording from genetically identified 5-HT neuron with electrically evoked 5-HT release, under physiological recording conditions (right). **(I)** Representative averaged traces from one example neuron depicting current responses to optogenetic stimulation of LHb afferents at two frequencies. Note that EPSCs are truncated for clarity to focus on direct comparison of the outward GIRK conductance. **(J)** Representative averaged traces from another example neuron depicting current responses to optogenetic stimulation of LHb afferents at four frequencies. **(K)** Total charge evoked by LHb stimulation as a functino of optogenetic stimulation frequency. Each line depicts one neuron.

**Figure S11:**
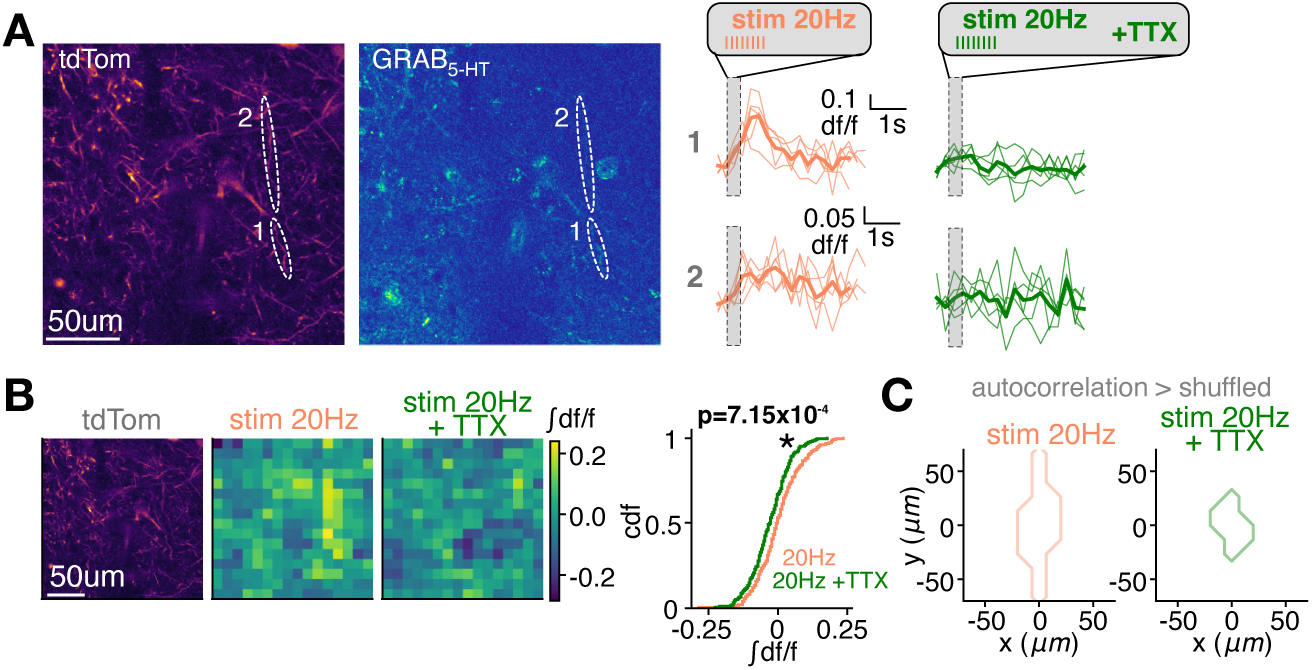
GRAB serotonin biosensor imaging reveals that 5-HT release is action potential-dependent. Related to Figure 5. **(A)** Left: Max-projection images of whole-frame tdTomato fluorescence and GRAB fluorescence during 20Hz electrical stimulation (8 pulses). Right: GRAB sensor responses to electrical stimulation at 20Hz before and after TTX washon for the ROIs depicted on the left. Thin traces represent individual trials and thick traces represent means across all trials. **(B)** Example whole-frame tdTomato fluorescence and binned GRAB responses to 20Hz electrical stimulation before and after TTX washon (left), and cumulative distribution of GRAB responses across pixels during 20Hz electrical stimulation before and after TTX washon (right; Kolmogorov-Smirnov test displayed). **(C)** Spatial autocorrelation plots for the GRAB fluorescence frames shown in B. Autocorrelation values are normalized from 0 to 1 for presentation.

**Figure S12.**
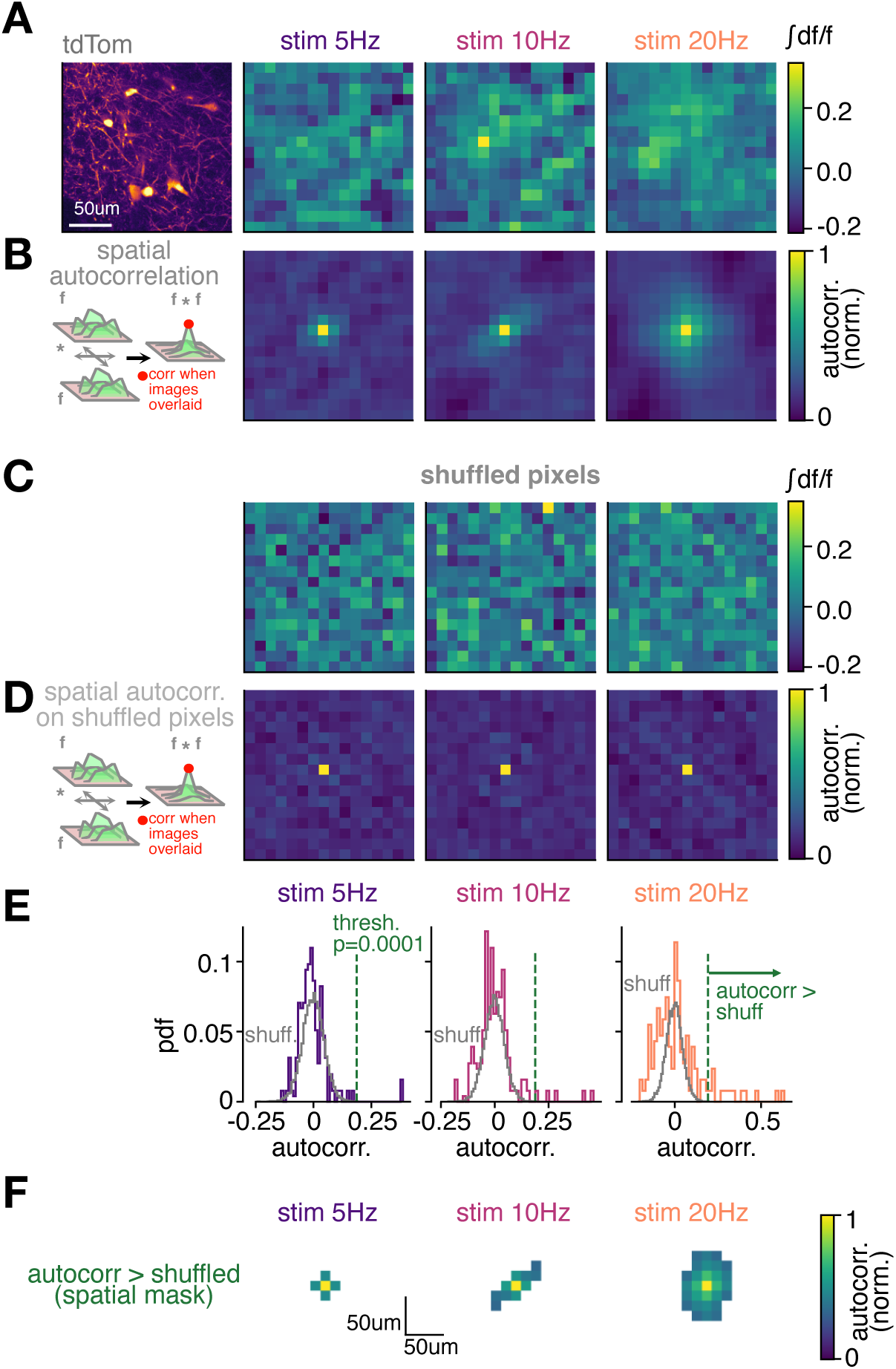
Generating a mask for statistically significant autocorrelation regions with a pixel shuffling approach. Related to Figure 5. **(A)** Example region showing whole-frame tdTomato fluorescence (left) and binned GRAB responses to 5, 10 and 20Hz electrical stimulation (8 pulses; right). **(B)** Spatial autocorrelation plots for the frames shown in A. Autocorrelation values are normalized from 0 to 1 for presentation. **(C)** Same as A, but the spatial location of pixels are shuffled. **(D)** Spatial autocorrelation plots for the pixel-shuffled frames shown in C. Autocorrelation values are normalized from 0 to 1 for presentation. **(E)** Probability distribution function of raw autocorrelation values for all pixels, for the spatial autocorrelations performed on pixel-shuffled GRAB responses (grey; computed from D) and for the spatial autocorrelations performed on GRAB responses (colors correspond to labeled electrical stimulation frequencies; computed from B). Pixels to the right of the green threshold have autocorrelation values that statistically exceed those of the shuffled distribution (p=0.0001). **(F)** Spatial mask for statistically significant autocorrelation values in B, where only pixels that exceed the significance threshold compared to the pixel-shuffled noise distribution (p=0.0001, as computed in E) are shown.

**Figure S13:**
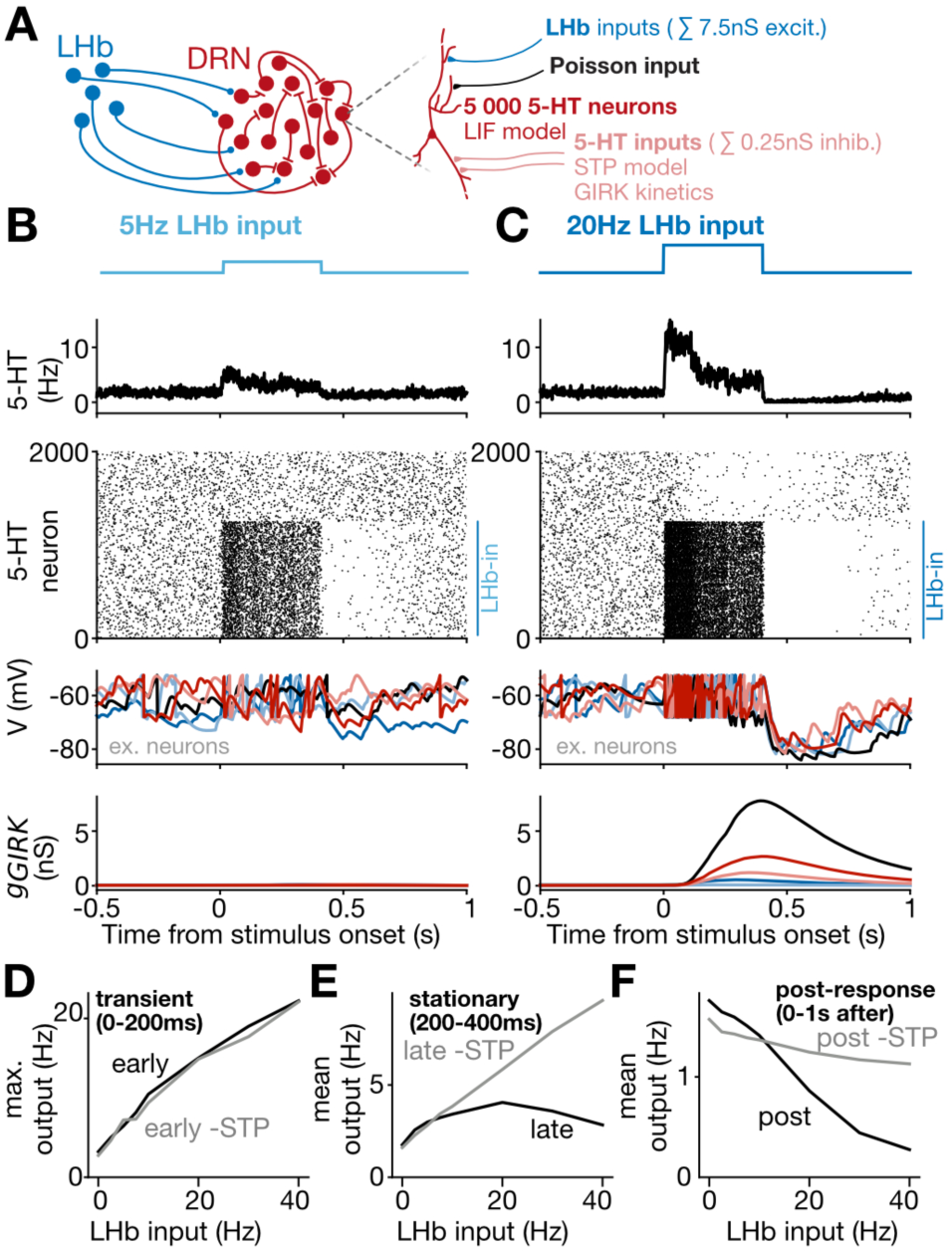
Nonlinear transformation of habenulo-raphe afferents generated by recurrent plasticity rules, with tenfold decrease in recurrent inhibitory connection probability (sum 0.25nS conductance received by each serotonin neuron). Related to Figure 6. **(A)** Schematic of the LIF network model. **(B)** Results of a simulation where a 5Hz step input from LHb was delivered to a subset of serotonin neurons. **(C)** Results of a simulation where a 20Hz step input from LHb was delivered to a subset of serotonin neurons. **(D)** Input-output transformation for the transient (0-200ms) response phase, with (black) and without (grey) STP rules at recurrent serotonergic connections. **(E)** Input-output transformation for the stationary (200-400ms) response phase, with (black) and without (grey) STP rules at recurrent serotonergic connections. **(F)** Input-output transformation for the post-response (0-1s after stimulus) phase, with (black) and without (grey) STP rules at recurrent serotonergic connections.

**Figure S14.**
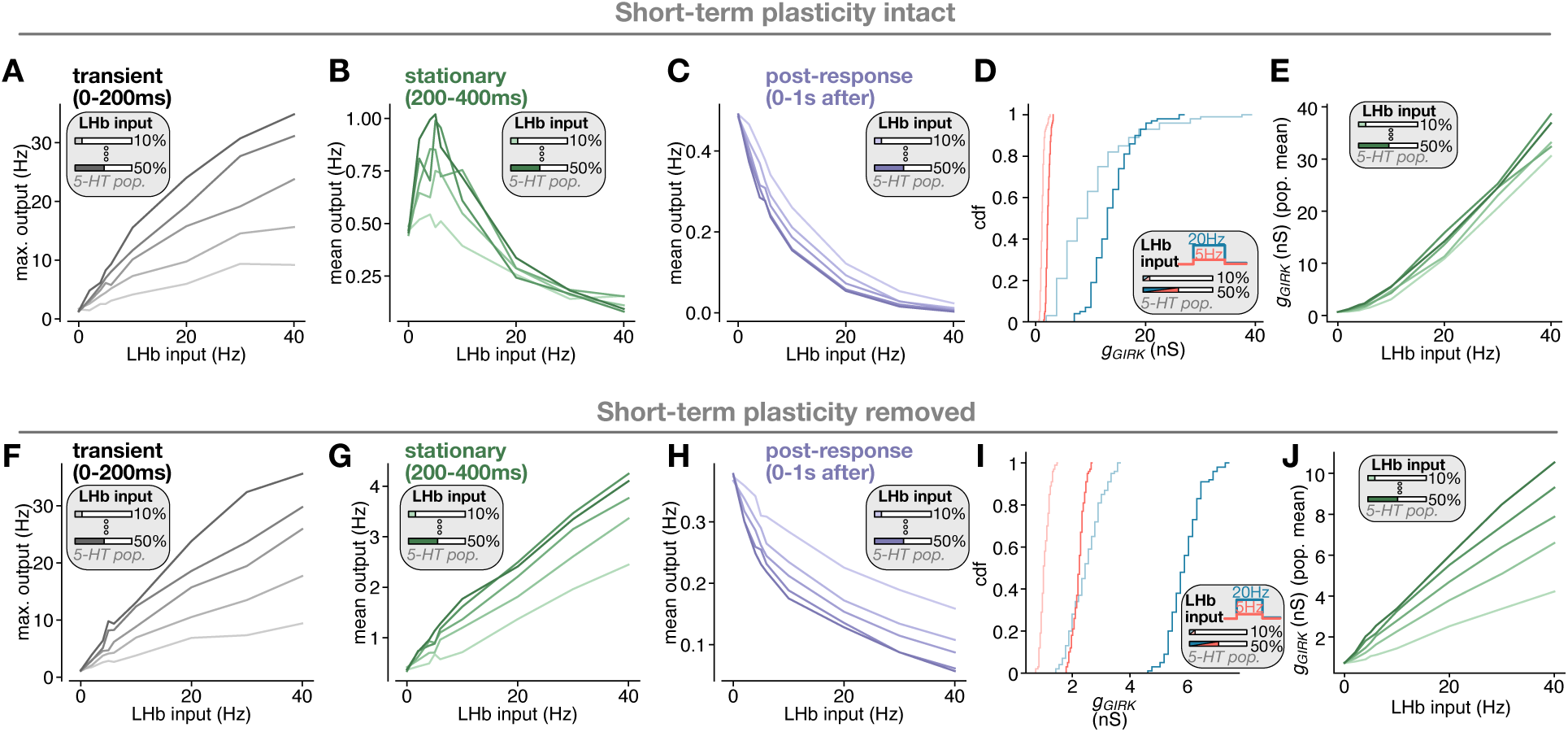
The input-output function of the habenulo-raphe pathway is largely invariant to the fraction of 5-HT neurons innervated by LHb, and this invariance depends on recurrent short-term plasticity. Related to network simulations depicted in Figure 6. Panels A-E depict results of simulations where short-term plasticity at 5-HT-5-HT connections is intact. **(A)** Input-output relationship of 5-HT neurons during the transient (0-200ms) phase of LHb input. Colors denote different fractions of 5-HT neurons receiving LHb input (shown in inset; 10-50%). **(B)** Same as A, but during stationary phase of LHb input (200-400ms). **(C)** Same as A, but during the post-response period of LHb input (0-1s after). **(D)** Cumulative distribution function of GIRK conductance across simulated 5-HT neurons, for 5Hz LHb input (red) and 20Hz LHb input (blue), and for fractions of 5-HT neurons innervated by LHb at 10% (lighter shade) and 50% (darker shade) (all combinations depicted in inset). Note how GIRK conductance depends primarily on LHb input frequency, instead of on LHb input fraction. **(E)** Relationship between LHb input frequency and GIRK conductance (population mean) as LHb input fraction is varied (colors denote distinct input fractions, shown in inset), demonstrating that population GIRK conductance is largely invariant to LHb input fraction. Panels D-H depict results of simulations where short-term plasticity at 5-HT-5-HT connections has been removed (i.e. linearized responses). **(F)** Input-output relationship of 5-HT neurons during the transient (0-200ms) phase of LHb input. Colors denote different fractions of 5-HT neurons receiving LHb input (shown in inset; 10-50%). **(G)** Same as A, but during stationary phase of LHb input (200-400ms). **(H)** Same as A, but during the post-response period of LHb input (0-1s after). **(I)** Cumulative distribution function of GIRK conductance across simulated 5-HT neurons, for 5Hz LHb input (red) and 20Hz LHb input (blue), and for fractions of 5-HT neurons innervated by LHb at 10% (lighter shade) and 50% (darker shade) (all combinations depicted in inset). Note how without short-term plasticity, GIRK conductance depends on both LHb input frequency and LHb input fraction. **(J)** Relationship between LHb input frequency and GIRK conductance (population mean) as LHb input fraction is varied (colors denote distinct input fractions, shown in inset), demonstrating that without short-term plasticity dynamics, the population GIRK conductance is not invariant to LHb input fraction.

**Figure S15:**
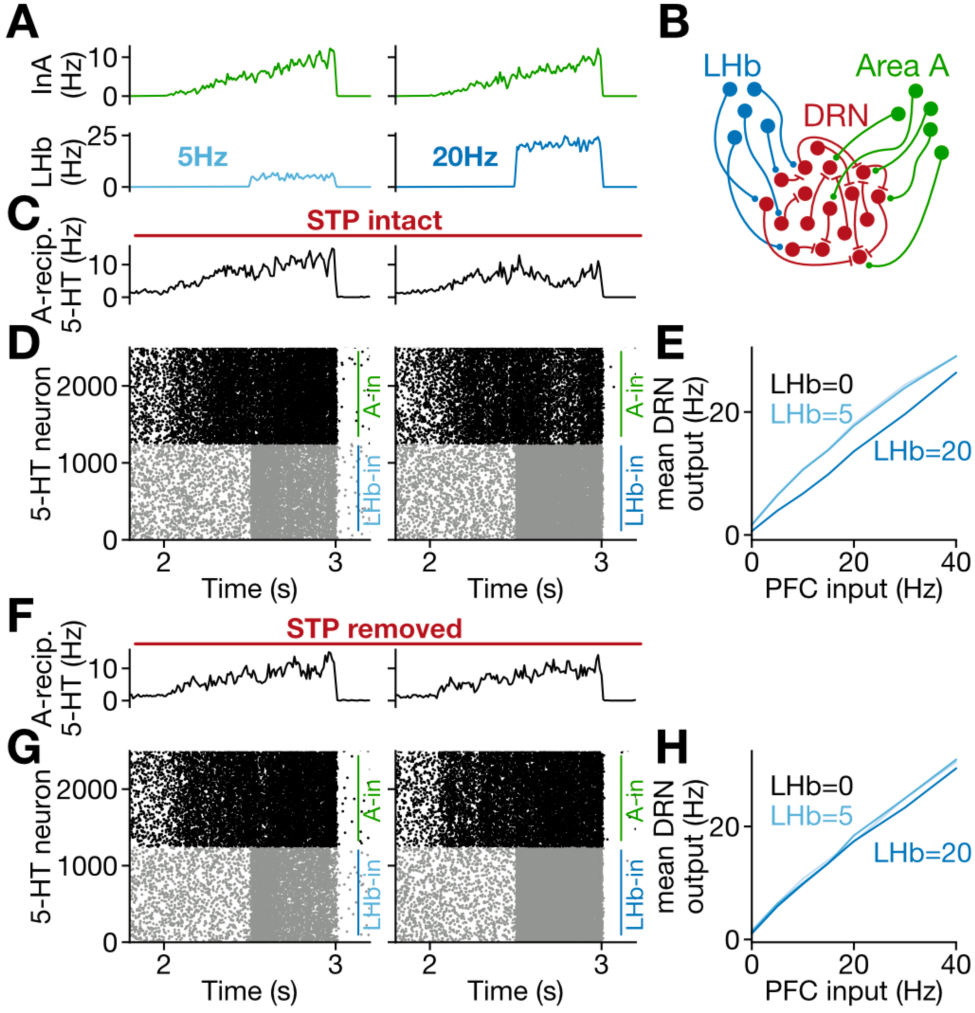
Plasticity-dependent winner-take-all dynamics in raphe, with tenfold decrease in recurrent inhibitory connection probability (sum 0.25nS conductance received by each serotonin neuron). **(A)** Simulations with LHb inputs of either 5Hz (left) and 20Hz (middle) were delivered to serotonin neurons during ramp input from area A. Schematic of LIF network model with two inputs (right). Related to Figure 7. **(B)** Mean responses of serotonin neurons to 5Hz (left) and 20Hz (right) input with STP rules intact. **(C)** Raster plot showing serotonin neuron responses to 5Hz (left) and 20Hz (right) input with STP rules intact. **(D)** Input-output function, for input A, with STP rules intact and coincident LHb input at 0, 5 or 20Hz. **(E)** Mean responses of serotonin neurons to 5Hz (left) and 20Hz (right) input with STP rules removed. **(F)** Raster plot showing serotonin neuron responses to 5Hz (left) and 20Hz (right) input with STP rules removed. **(G)** Input-output function, for input A, with STP rules removed and coincident LHb input at 0, 5 or 20Hz.

**Figure S16:**
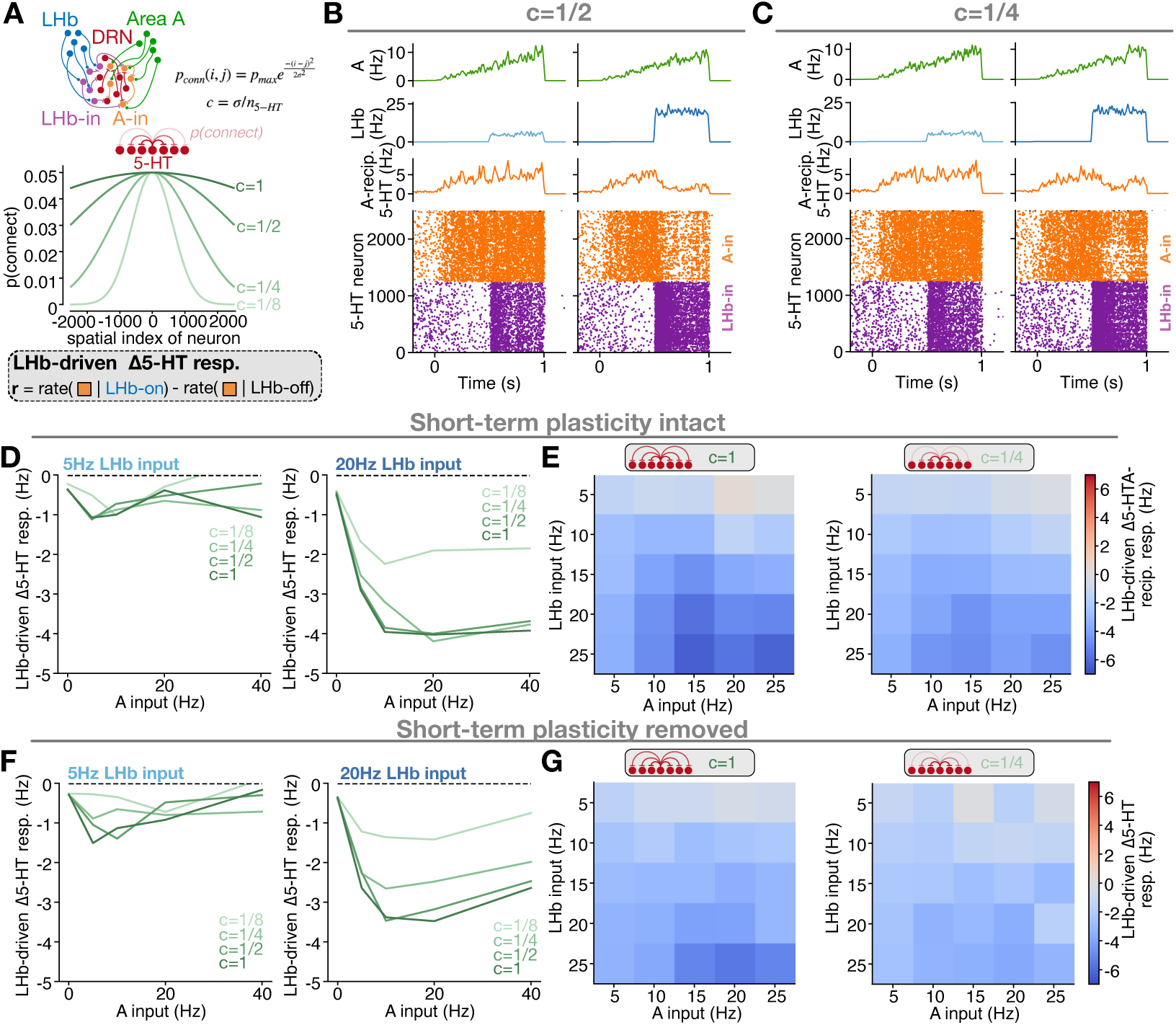
Winner-take-all dynamics in the DRN persist over a wide range of spatial connectivity profiles for recurrent inhibition. Related to network simulations shown in Figure 7. **(A)** Schematic of network simulation, with LHb and Area A providing input to 5-HT neurons (top). Plot demonstrating how 5-HT-5-HT recurrent connectivity probability varies with distance, as spatial connectivity profiles are parameterized (bottom). **(B)** Simulation with more global recurrent spatial connectivity of *c=*1/2. Shown are mean firing rates of A and LHb inputs (top); mean firing rates of A-recipient 5-HT neurons (middle); and raster plot depicting firing rates of 2500 5-HT neurons receiving either LHb input (purple) or A input (orange) (depicted is the 50% of 5-HT neurons receiving direct input from either LHb or PFC). **(C)** Simulation with more local recurrent spatial connectivity of *c=*1/4. Layout of panels is identical to in B. D-E quantify the strength of winner-take-all dynamics as a function of recurrent spatial connectivity profiles, while short-term plasticity at recurrent connections is intact. **(D)** LHb-evoked change in A-recipient 5-HT neuron responses, as a function of the rate of A input, for 5Hz (left) and 20Hz (right) LHb input. Spatial connectivity profile is varied (shown in legend), demonstrating that winner-take-all dynamics are stable across a wide range of spatial recurrent connectivity profiles. **(E)** Strength of winner-take-all effect (quantified as LHb-evoked change in A-recipient 5-HT neuron responses) as A input rate and LHb input rate are varied. Shown are recurrent spatial connectivity values of *c*=1 (left) and *c*=1/4 (right). F-G quantify the strength of winner-take-all dynamics as a function of recurrent spatial connectivity profiles, while short-term plasticity at recurrent connections has been removed (i.e. linearized responses). **(F)**: LHb-evoked change in A-recipient 5-HT neuron responses, as a function of the rate of A input, for 5Hz (left) and 20Hz (right) LHb input. Spatial connectivity profile is varied as depicted in the legend, demonstrating that with short-term plasticity removed, the winner-take-all dynamics vary over the tested range of spatial recurrent connectivity profiles. **(G)** Strength of winner-take-all effect (quantified as LHb-evoked change in A-recipient 5-HT neuron responses) as A input rate and LHb input rate are varied. Shown are recurrent spatial connectivity values of *c*=1 (left) and *c*=1/4 (right). Note that the winner-take-all effect is weaker than with short-term plasticity intact (E).

**Figure S17:**
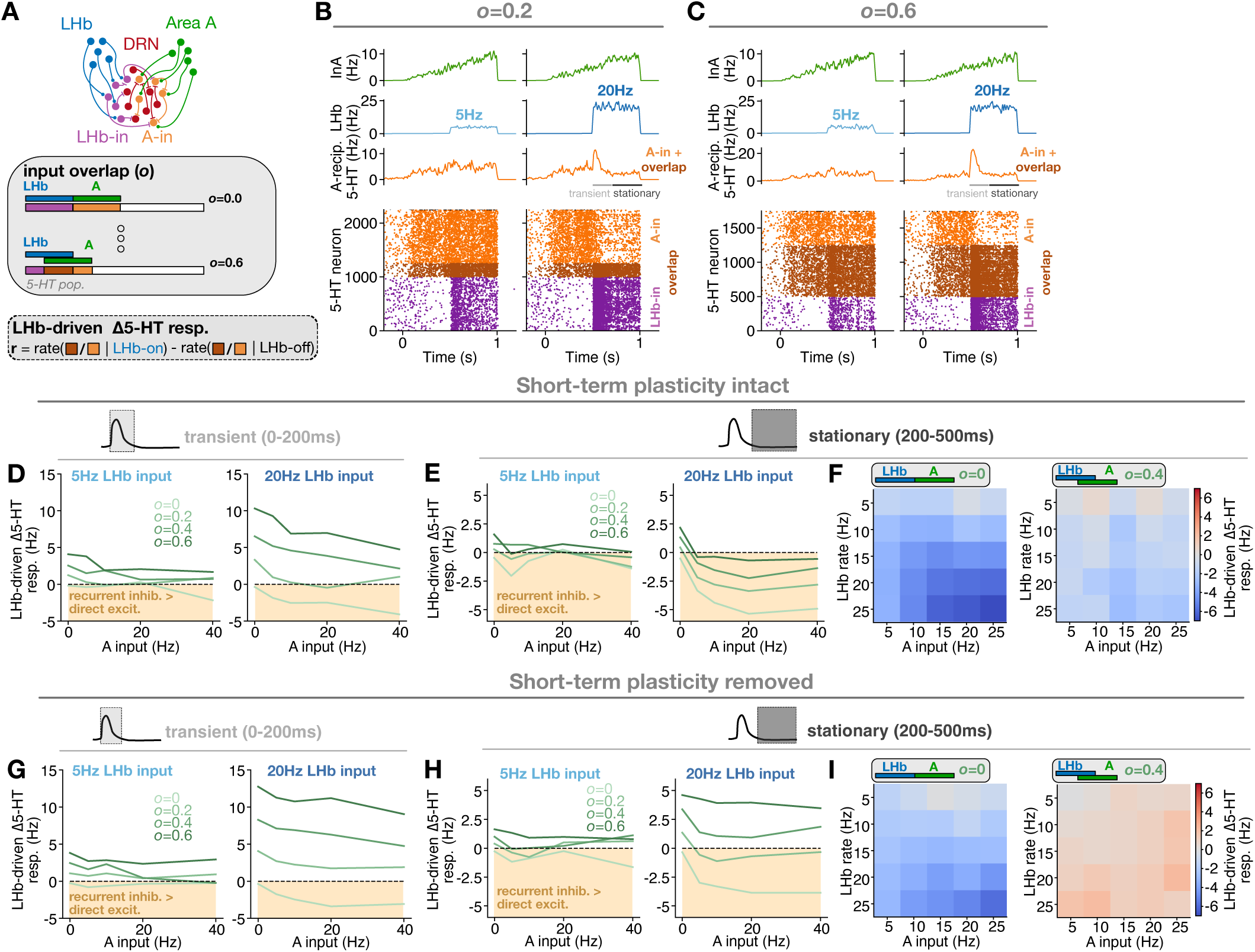
Winner-take-all dynamics in the DRN persist over a wide range of overlap fractions between competing inputs, a phenomenon which depends on recurrent short-term plasticity. Related to network simulations shown in Figure 7. **(A)** Schematic of network simulation, with LHb and Area A providing input to 5-HT neurons (top). Bottom contains schematic demonstrating how the overlap between LHb and A inputs onto 5-HT neurons is varied from 0-60%. **(B)** Simulation with small input overlap of *o=*0.2. Shown are mean firing rates of A and LHb inputs (top); mean firing rates of A-recipient 5-HT neurons (middle); and raster plot depicting firing rates of 2500 5-HT neurons receiving either LHb input (purple) or A input (orange) (depicted is the 50% of 5-HT neurons receiving direct input from either LHb or PFC). B Simulation with larger input overlap of *o=*0.6. Layout of panels is identical to in B. D-F quantify the strength of winner-take-all dynamics as a function of input overlap fraction, while short-term plasticity at recurrent connections is intact. **(D)** LHb-evoked change in A-recipient 5-HT neuron responses during the transient phase (0-200ms), as a function of the rate of A input, for 5Hz (left) and 20Hz (right) LHb input. Input overlap is varied (shown in legend). **(E)** LHb-evoked change in A-recipient 5-HT neuron responses during the stationary phase (200-500ms), as a function of the rate of A input, for 5Hz (left) and 20Hz (right) LHb input. Input overlap is varied (shown in legend). **(F)** Strength of winner-take-all effect (quantified as LHb-evoked change in A-recipient 5-HT neuron responses) as A input rate and LHb input rate are varied. Shown are input overlap values of *o*=0 (left) and *o*=0.4 (right), illustrating winner-take-all dynamics (ie negative 5-HT modulation) for these input fractions. G-I quantify the strength of winner-take-all dynamics as a function of input overlap fraction, while short-term plasticity at recurrent connections has been removed (i.e. linearized responses). **(G)** LHb-evoked change in A-recipient 5-HT neuron responses during the transient phase (0-200ms), as a function of the rate of A input, for 5Hz (left) and 20Hz (right) LHb input. Input overlap is varied (shown in legend). **(H)** LHb-evoked change in A-recipient 5-HT neuron responses during the stationary phase (200-500ms), as a function of the rate of A input, for 5Hz (left) and 20Hz (right) LHb input. Input overlap is varied (shown in legend). **(I)** Strength of winner-take-all effect (quantified as LHb-evoked change in A-recipient 5-HT neuron responses) as A input rate and LHb input rate are varied. Shown are input overlap values of *o*=0 (left) and *o*=0.4 (right), illustrating that winner-take-all dynamics are no longer present with moderate input overlap of *o*=0.4 when short-term plasticity is removed (compare with F).

**Figure S18:**
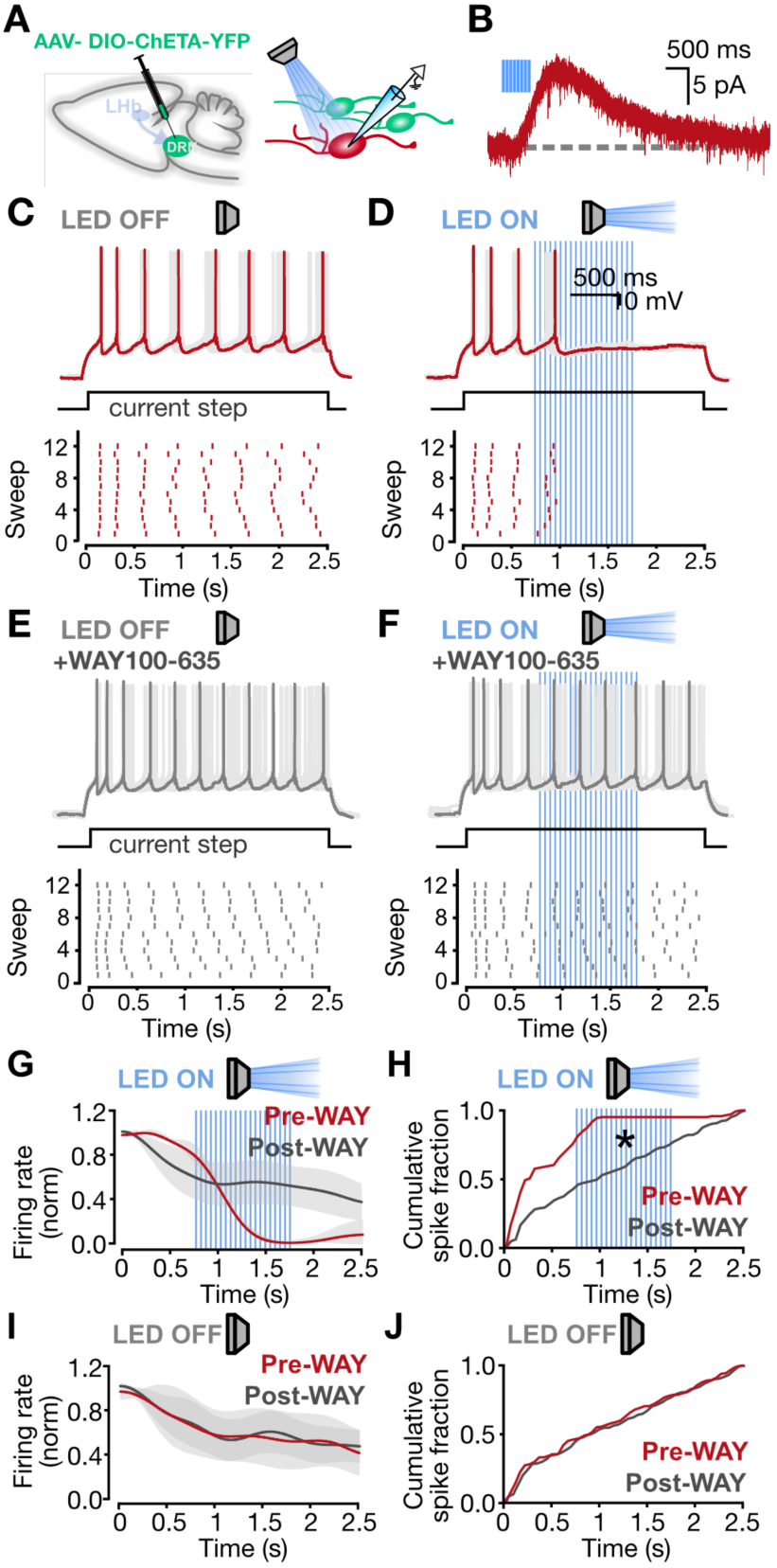
Recurrent inhibition gates spiking activity of 5-HT neurons. **(A)** Schematic of experimental design. Left, AAV-DIO-ChETA-YFP is injected into the DRN at a 2:1 ratio with Mannitol. Right, whole-cell recordings from ChETA-negative 5-HT neurons. **(B)** Recurrent 5HT1AR-mediated current in ChETA-negative 5-HT neuron. **(C)** Spiking of a 5-HT neuron in response to current injection. Below, raster plot showing the same neuron’s activity across trials. **(D)** Spiking of a 5-HT neuron in response to current injection and 20Hz optical activation of CHETA-positive 5-HT neurons. Below, raster plot showing the same neuron’s activity across trials.**(E)** Spiking of a 5-HT neuron in response to current injection in the presence of WAY100-635. Below, raster plot showing the same neuron’s activity across trials. **(F)** Spiking of a 5-HT neuron in response to current injection and 20Hz optical activation of CHETA-positive 5-HT neurons, in the presence of WAY100-635. Below, raster plot showing the same neuron’s activity across trials. **(G)** Smoothed firing rate (normalized) over time for pre- and post-WAY recordings with LED stimulation (mean +- SE, n=3 neurons). **(H)** Cumulative spike fraction in pre- and post-WAY recordings with LED stimulation (n=3 neurons, 2-sample Kolmogorov-Smirnov test: p = 7.4e-15). **(I)** Smoothed firing rate (normalized) over time for pre- and post-WAY recordings without LED stimulation (mean +- SE, n=3 neurons). **(J)** Cumulative spike fraction in pre- and post-WAY recordings with LED stimulation (3 neurons, 2-sample Kolmogorov-Smirnov test: p = 0.85).

**Figure S19:**
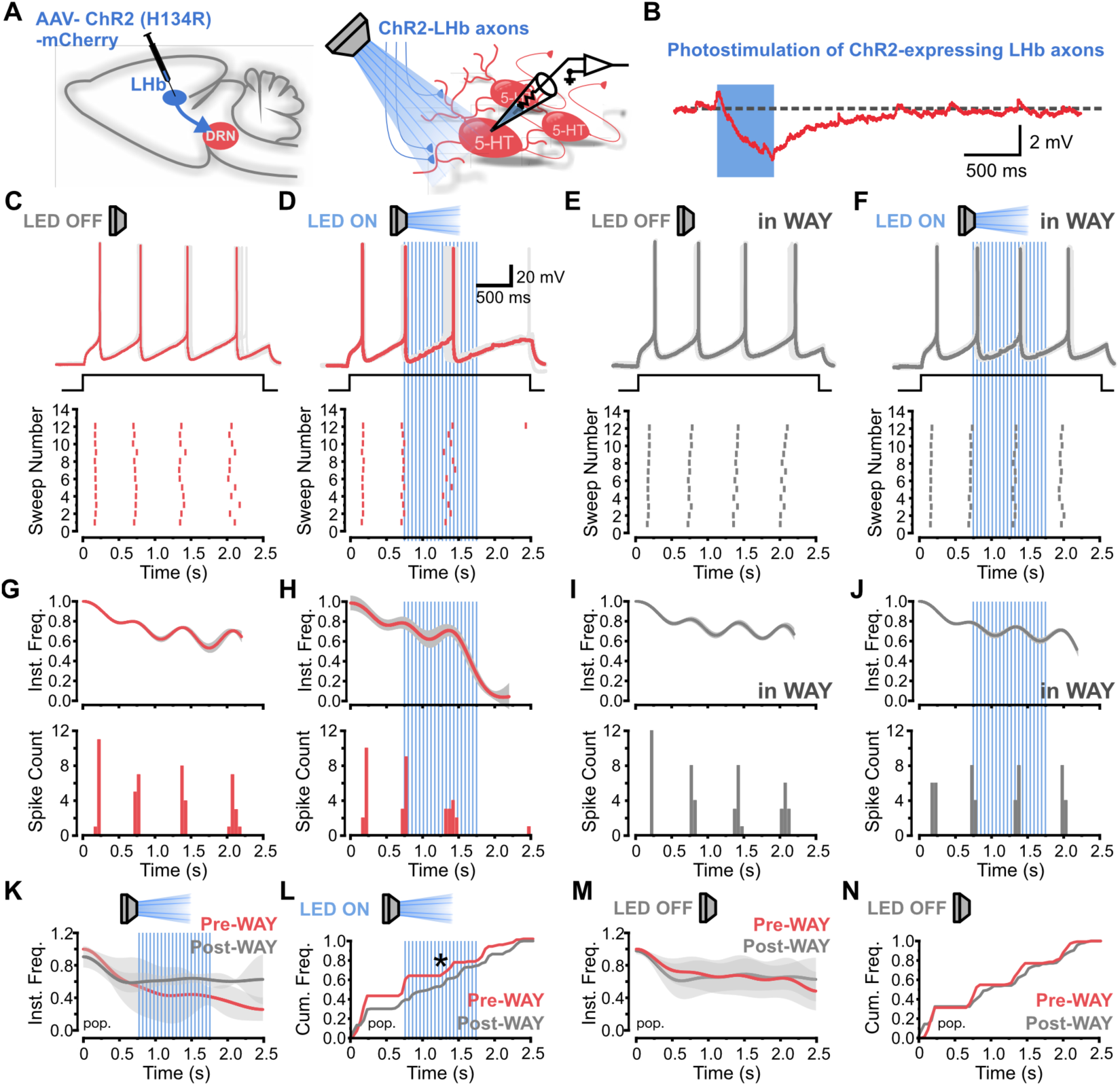
LHb-driven, 5HT1AR mediated, recurrent feedforward modulation of 5-HT firing activity in the DRN. **(A)** Schematic of experimental design. Left, AAV-ChR2(H134R)-mCherry is injected into the LHb. Right, whole-cell recordings from TdT-positive 5-HT neurons. **(B)** LHb-driven heterosynaptic 5HT1AR-mediated potential in SERT-TdT 5-HT neuron. **(C)** Spiking of a 5-HT neuron in response to a 2.5 s current step (scale, 20 mV, 500ms). Below, raster plot of the same 5-HT neuron spiking. **(D)** Spiking of a 5-HT neuron in response to a 2.5 s current step with LED activation of ChR2-expressing LHb terminals in the slice for 1 sec in the middle of the current step. Below, raster plot of the same 5-HT neuron spiking. **(E)** Spiking of a 5-HT neuron in response to a 2.5 s current step in the presence of WAY 100-635. Below, raster plot of the same 5-HT neuron spiking. **(F)** In the presence of WAY 100-635, spiking of a 5-HT neuron in response to a 2.5 s current step with LED activation of ChR2-expressing LHb terminals in the slice for 1 sec in the middle of the current step. Below, raster plot of the same 5-HT neuron spiking. **(G)** Instantaneous frequency line (top) and spike count histogram plots (below) from recording in **C**. **(H)** Instantaneous frequency line (top) and spike count histogram plots (below) from recording in **D**. **(I)** Instantaneous frequency line (top) and spike count histogram plots (below) from recording in **E**. **(J)** Instantaneous frequency line (top) and spike count histogram plots (below) from recording in **F**. **(K)** Instantaneous frequency population plot in pre- and post-WAY recordings with LED stimulation (n=3 neurons, mean +- SE). **(L)** Cumulative frequency population plot in pre- and post-WAY recordings with LED stimulation (3 cells, p = 0.0017, 2-sample Kolmogorov-Smirnov test). **(M)** Instantaneous frequency population plot in pre- and post-WAY recordings in the absence of LED stimulation (n=3 neurons, mean +- SE). **(N)** Cumulative frequency population plot in pre- and post-WAY recordings in the absence of LED stimulation (3 cells, p = 0.07, 2-sample Kolmogorov-Smirnov test).

**Figure S20:**
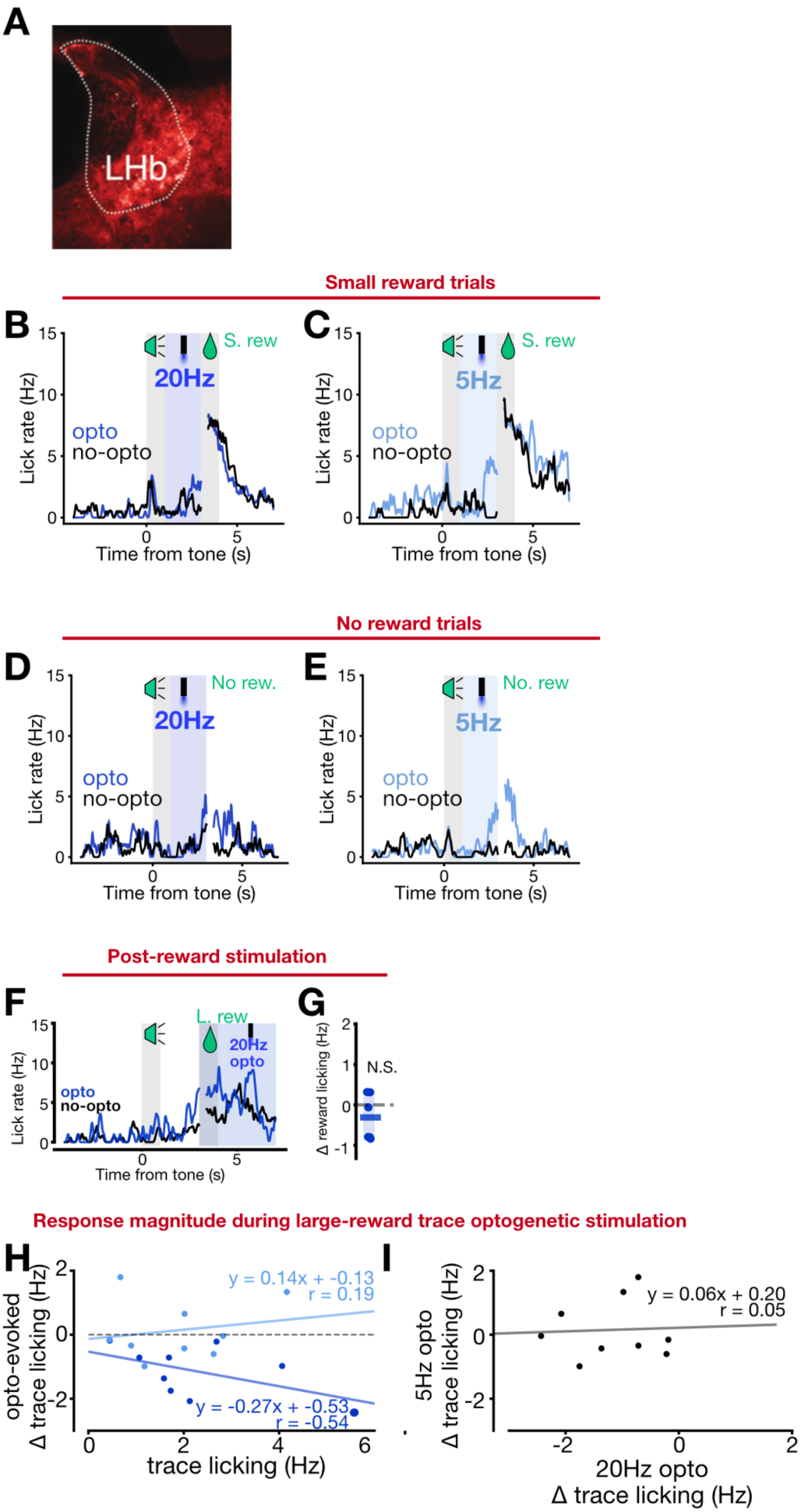
The habenulo-raphe pathway disrupts reward-conditioned responses in a value-dependent manner and does not affect unconditioned response. **(A)** Representative example of LHb injection site. **(B)** Mean licking across all small reward trials for a single mouse depicting control trials (black) or 20Hz optogenetic stimulation trials (blue). **(C)** Mean licking across all small reward trials for a single mouse depicting control trials (black) or 5Hz optogenetic stimulation trials (blue). **(D)** Mean licking across all no reward trials for a single mouse depicting control trials (black) or 20Hz optogenetic stimulation trials (blue). **(E)** Mean licking across all no reward trials for a single mouse depicting control trials (black) or 5Hz optogenetic stimulation trials (blue). **(F-G)** depict post-reward optogenetic stimulation experiments. **(F)** Mean licking across all large reward trials for a single mouse depicting control trials (black) or trials where 20Hz optogenetic stimulation was delivered during the post-reward period (blue). **(G)** Change in post-reward licking during optogenetic stimulation trials, compared to pre-stimulated baseline period (n=6 mice, paired two-sided t-test, unstimulated vs stimulated reward licking). (**H**) Correlation between mean trace licking (large reward trials) and optogenetic effect size for 20Hz stimulation (dark blue) and 5Hz stimulation (light blue) for trace stimulation during large reward trials. (**I**) Correlation between effect size for 20Hz stimulation and 5Hz stimulation for trace stimulation during large reward trials.

